# A Principled Statistical Framework for Analyzing Spatial Patterns in Spatially Resolved Multi-Omics

**DOI:** 10.64898/2026.08.04.742894

**Authors:** Jinpu Li, Mauminah Raina, Yiqing Wang, Shuai Zeng, Yang Yu, Xiufan Yu, Xiaojie Jin, Yuzhou Chang, Daniel Feliciano, Jonathan Himmelfarb, Ana C. Ricardo, Patrick H. Nachman, Miguel Vazquez, M. Luiza Caramori, Laura Barisoni, Matthias Kretzler, Sanjay Jain, Pierre C. Dagher, Tarek M. El-Achkar, Michael T Eadon, Human Biomolecular Atlas Program, Kidney Precision Medicine Project, Ricardo Melo Ferreira, Qin Ma, Juexin Wang, Dong Xu

## Abstract

Emerging spatial multi-omics technologies enable the profiling of molecular variation within its tissue context, yet existing methods for identifying spatially variable features lack principled approaches to experimental design and cross-sample inference. Here, we present STORM, a principled **S**tatistical **TO**ol for spatially **R**esolved **M**ulti-omics, for rigorously analyzing spatial patterns in spatial multi-omics research. STORM incorporates a robust and efficient nonparametric test that quantifies local deviations in molecular feature measurements to detect spatial dependence across transcriptomic and proteomic data. It further estimates an interpretable spatial effect size, supports power calculations for both spatial locations and biological replicates, and enables formal group-level comparisons. In several simulated and experimental spatial multi-omics case studies, STORM demonstrates reliable performance in detecting spatial structures while offering quantitative support for study design decisions. Overall, STORM provides a principled statistical framework that unifies spatial hypothesis testing, effect size estimation, power analysis, and experimental design for spatial multi-omics data.

## Introduction

The emergence of spatial multi-omics has substantially expanded the capabilities to characterize molecular heterogeneity within intact tissues. Spatially variable features (SVFs) encompass genes and other molecular modalities whose expression patterns are non-randomly distributed within tissue sections^1–3^. Identifying SVFs is critical for delineating functionally distinct tissue regions, characterizing cell-cell communication networks, and uncovering novel biomarkers for diseases such as cancer and neurodegenerative disorders^4–6^. Numerous statistical methods have been developed for this purpose, including regression-based approaches such as spatialDE^7^ and SPARK^8^, graph-based approaches such as BinSpect^9^ and scGCO^10^, and permutation-based methods such as Trendsceek^11^ and SpaGene^12^. Extensions such as SPARK-X^13^ and HEARTSVG^14^ combine multiple statistical strategies to improve detection performance. However, several critical gaps remain in methodological development.

Firstly, there is currently no widely adopted statistical framework that supports principled power and sample-size justification for the analysis of SVFs^15^. Existing SVF detection methods are primarily designed for *post hoc* hypothesis testing and often depend on pilot data that may not be available during study design. Even though some comparison works provide experience guidelines, investigators lack formal guidance on determining the minimum number of spots required within a sample or the number of biological replicates required across groups to achieve adequate statistical power^16^. This limitation directly affects resource allocation, experimental feasibility, and reproducibility.

Secondly, most available approaches focus on single-sample inference and provide limited support for analyses across biological replicates or conditions. A small number of methods address related comparison tasks, but their scope remains restricted. For example, DESpace2^17^ examines whether spatial patterns between conditions are identical, whereas the test is based on predefined spatial domains. Other recently developed methods, such as SPADE^18^, scHOT^19^, and STcompare^20^, are limited to pairwise comparisons between two genes or two samples, with only one sample representing each condition. Another commonly used alternative is to combine per-sample p-values through meta-analysis to assess whether spatial patterns are consistently present within a group. However, this approach does not support formal between-group comparisons because effect size estimates are typically unavailable. The absence of a unified framework for both individual-level and group-level spatial inference complicates the design and interpretation of case–control or multi-condition spatial studies.

Thirdly, current methods predominantly report p-values without accompanying quantitative measures of spatial effect size. Unlike differential expression analysis in bulk RNA sequencing, where fold change provides a measure of magnitude, spatial analyses often lack an interpretable metric that quantifies the strength of spatial dependence. Consequently, statistical significance can be strongly influenced by sample-specific spot counts, limiting comparability across samples and studies, and constraining biological interpretation. Previous methods proposed the concept of spatial effect size as the proportion of spatial variance to total variance under a Gaussian process model^7,21,22^. However, this formulation is model-dependent, relying heavily on the choice of kernel and normalization methods. Its simplified variance structure and limited exploration of zero inflation further reduce interpretability and generalizability.

To address these challenges, we introduce STORM (**S**tatistical **TO**ol for spatially **R**esolved **M**ulti-omics), a statistical framework for analyzing spatial expression patterns in spatial multi-omics data. The primary contribution of STORM is the integration of spatial hypothesis testing with a principled structure for guiding experimental design and group-level inference in the analysis of spatial patterns. To the best of our knowledge, it is the first unified statistical framework that enables prospective power analysis and sample size determination at both the spot and biological replicate levels, allowing investigators to justify experimental design decisions before data collection and to conduct formal comparisons of spatial dependence between groups. In multiple studies using spatial omics at cellular or subcellular resolution, up to the million-level scale, STORM demonstrates robust detection of spatial patterns and supports a statistically grounded experimental design. Together, STORM provides a coherent statistical foundation that connects experimental planning, spatial dependence testing, and group-level comparison. By linking these components within a single framework, STORM addresses key limitations in current spatial multi-omics methodology while remaining compatible with existing analytical workflows.

## Results

### STORM Effectively and Efficiently Detects Spatial Patterns

#### Overview of STORM

STORM uses a nonparametric test to detect SVFs by assessing whether feature expression is independent of spatial location (Fig. 1A). It leverages local neighborhood structures to quantify how expression at each spatial location deviates from its surrounding context, and aggregates this information across the tissue to examine the presence of spatial organization. A corresponding effect size is computed as the proportion of variance explained by the spatial dependence, quantifying the contribution of spatial organization to overall expression variability. A parametric approximation is also provided when expression distributions approximate normality. Details are provided in Methods, and technical details are provided in the Supplementary Notes. A key innovation of STORM is its ability to connect statistical inference with experimental design. By explicitly characterizing how spatial signal strength relates to detectability, STORM enables group-level inference and prospective power analysis and sample size determination for both spatial resolution (number of spots) and biological replication. This enables planning spatial multi-omics studies in a statistically grounded manner prior to data collection.

**Fig. 1:**
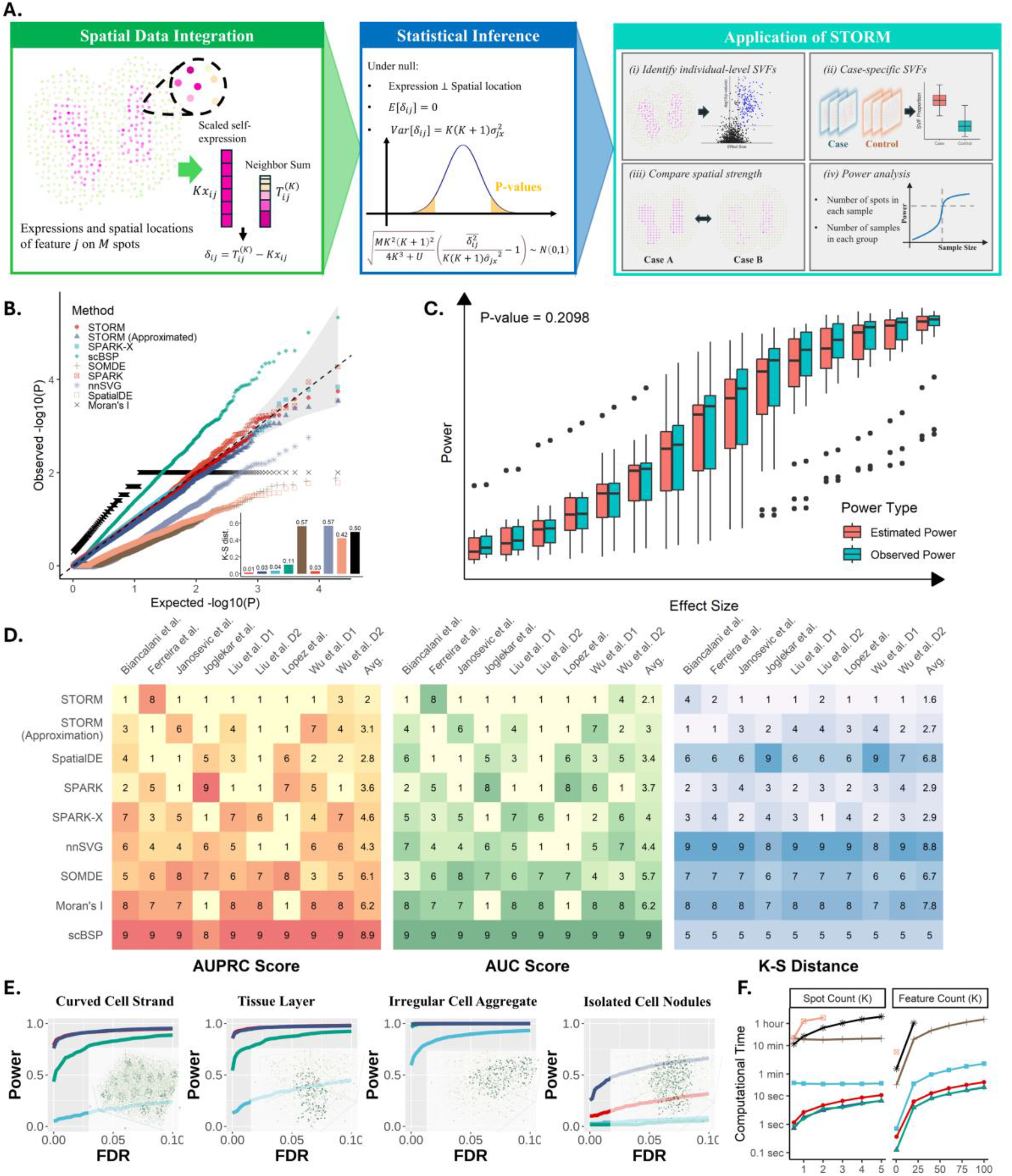
Overview of STORM and simulations. **A:** Schematic of STORM. **B:** Type I error of STORM and other methods under the null hypothesis. **C**: Type II error rate of STORM with approximation under the alternative hypothesis. **D.** Rankings based on AUPRC (left), AUC (middle), and K-S distance (right) on nine benchmark datasets. **E:** Comparisons of statistical power (y-axis) against false discovery rate (x-axis) on 3D simulations with Isolated Cell Nodules patterns, Curved Cell Strand, Tissue Layer, and Irregular Cell Aggregate (left to right) with moderate pattern size, signal strength, and noise. **F**: Computational time (y-axis) for analyzing spatial omics data comprising 20,000 features across 3,000 spots, varying number of spots and features while keeping the other constant.

#### Calibration of hypothesis testing

To assess the statistical calibration of STORM for detecting SVFs, we first assessed its Type I error using a null simulation consisting of 10,000 features without spatial structure measured across 900 spots and compared its performance with seven other existing methods^7,8,13,21,23,24^, which were commonly used and demonstrated strong performances in previous benchmarking studies^25,26^. Quantile–quantile (Q–Q) plots of observed versus expected p-values showed that both STORM and its approximation produced well-calibrated p-values under null hypothesis. In contrast, SpatialDE, SOMDE, and nnSVG were overly conservative, whereas scBSP and Moran’s I exhibited inflated Type I error. Calibration was further quantified using the Kolmogorov–Smirnov (K–S) distance between empirical and uniform p-value distributions.

STORM and its approximation achieved the smallest K–S distances, followed by SPARK and SPARK-X, while other methods showed substantially poorer calibration. These results remained stable across a broad range of neighborhood sizes and in a high-resolution setting comprising 1,000,000 spots with 0.5% nonzero values (Supplementary Fig. 1A-B).

Unlike existing approaches, STORM enables rigorous theoretical estimation of statistical power. We validated its statistical rigor by comparing theoretical and empirical power across simulations with a broad range of statistical power controlled by altering effect sizes (Methods). Using a significance level of 0.05, empirical power was defined as the proportion of detected SVFs, while theoretical power was estimated using observed sample size and average effect size. A generalized linear mixed model showed no significant difference between observed and estimated power (p = 0.2098; Fig. 1C), demonstrating STORM’s ability for accurate power estimation. Notably, this evaluation focused on the approximation of STORM, which does not require prior spatial information and is therefore more practical for study design.

#### Benchmarking SVF detection accuracy

In addition to statistical calibration, we evaluated the detection accuracy of STORM on published datasets already used in a benchmark study^25^. These datasets were generated with scDesign3^27^ from nine source data covering nine distinct spatial masks, five histological tissue types, two spatial transcriptomics platforms, and a range of sequencing depths. Performance was assessed and ranked using the area under the receiver operating characteristic curve (AUC) and the area under the precision–recall curve (AUPRC) on each dataset (Fig. 1D, Supplementary Table 1), while the corresponding statistical calibration was evaluated and ranked using K-S distance on shuffled data. STORM achieved the highest ranking in seven of the nine benchmark datasets and ranked highest overall for both AUC and AUPRC, while its approximation ranked second for AUC and third for AUPRC. Although its relative ranking was lower for the Melo Ferreira et al. dataset^28^, STORM still achieved strong absolute accuracy (AUC = 0.98, AUPRC = 0.97). scBSP performed poorly due to the violation of its underlying assumption that most features have no spatial structure. The robustness of STORM was further evaluated across varying numbers of neighbors, with only minor differences observed (Supplementary Figure 1C).

Besides two-dimensional simulations, we also performed three-dimensional simulations that can better reflect emerging high-resolution spatial profiling applications. STORM and its approximation consistently achieved higher AUPRC and statistical power than scBSP and SPARK-X across a wide range of false discovery rates (Fig. 1E, Supplementary Figs. 1D and 2-4). Notably, STORM performed slightly worse than its approximation when detecting small, isolated cell nodules. This is because it uses a larger neighborhood, which averages signals beyond the nodules and weakens the local signal. Nevertheless, both versions still consistently outperformed competing methods.

#### Validation of group-level inference

We next evaluated two types of group-level inferences using a subset of three-dimensional simulations with 1,000 features. Case-control comparisons were used to test for spatial structure in the case group relative to the control group, while case-case comparisons assessed differences in spatial dependence strength between two groups of spatially structured features. Both STORM and its approximation consistently detected significant differences (Supplementary Tables 2–3). To assess performance with smaller sample sizes, we randomly sampled 3–20 features per group and repeated the analysis 1,000 times. In most settings, the empirical power exceeded 0.9 when the sample size was greater than 11, even when individual-level tests had limited power, demonstrating the substantial gain in statistical power achieved through group-level inference (Supplementary Tables 4–5). The empirical power was subsequently compared with theoretical estimates and showed good alignment (Supplementary Fig. 5).

Type I error was also well controlled. For case–case comparisons under the null (equal spatial effects, which correspond to the proportion of variance that can be explained by the spatial structure), STORM produced well-calibrated p-values, while its approximation was slightly conservative. For case–control null comparisons, both versions were well calibrated.

#### Computational scalability of STORM

We next evaluated computational efficiency by resampling low- and high-resolution datasets, respectively (10x Genomics Visium mouse brain with 31,053 genes and 2,696 spots; high-definition spatial transcriptomics mouse brain with 19,950 features and 181,367 spots). On a workstation equipped with a 2.00 GHz AMD EPYC 7713 processor, STORM and its approximation completed low-resolution analyses within 10 seconds, matching the performance of scBSP and substantially outperforming other methods (Fig. 1F). In high-resolution settings, the STORM approximation and scBSP remained the fastest approaches, with SPARK-X achieving comparable runtime only at very large scales (Supplementary Fig. 6). Although memory requirements varied across methods and resolutions, all four above methods remained computationally practical, completing analyses within minutes using reasonable memory resources.

### STORM Enables Sample Size Determination in Kidney Spatial Pattern Analysis

A central challenge in spatial multi-omics studies is determining whether the proposed experiments have sufficient statistical power to detect molecular features that exhibit spatially structured expression and to identify how these patterns vary across conditions. To demonstrate how STORM can guide study design, we performed a case study using kidney spatial transcriptomics datasets generated by the 10x Genomics platforms Visium^29^ and Xenium^30^, which differ substantially in spatial resolution, gene counts, and measurement density (Fig. 2A).

**Fig. 2:**
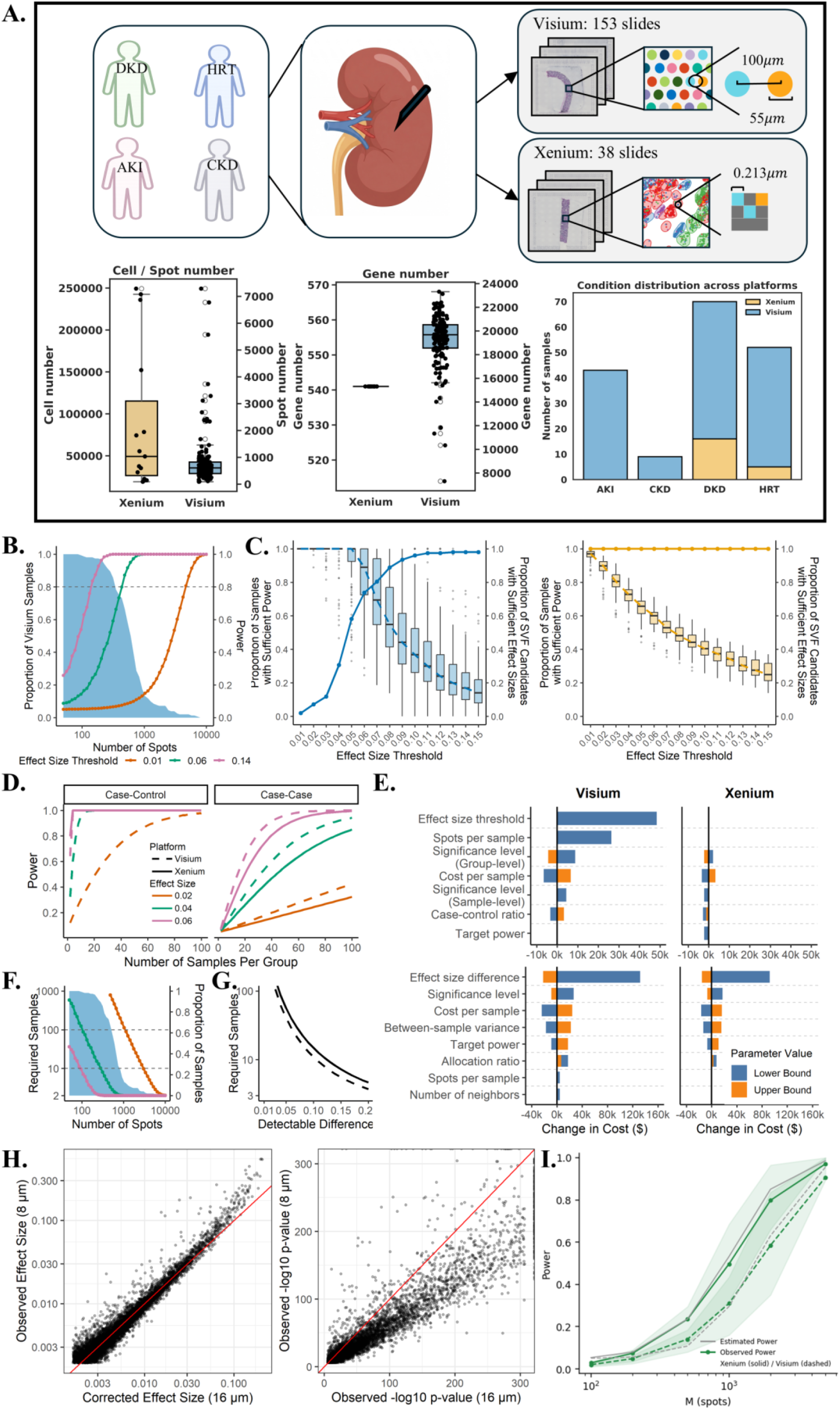
Power analysis and study design for SVF detection **A:** Overview of Visium and Xenium samples. **B.** Individual-sample power curves showing statistical power as a function of the number of spatial measurements for different effect size thresholds. Horizontal dashed lines indicate 80% power. **C:** Tradeoff between SVF detectability and sample-level feasibility across effect size thresholds. Boxplots show the distribution across samples of the proportion of SVFs (adjusted p-value < 0.05) exceeding each effect size threshold. Solid Lines indicate the proportion of samples with sufficient spatial resolution to achieve 80% power. **D:** Power curves for group-level comparisons. **E:** Tornado plot showing the sensitivity of the total cost estimate to variations in key design parameters. Bars represent the change in the estimated cost when each parameter is increased or decreased by 50% while holding other parameters fixed. Design parameters varied across significance levels ranging from 0.01 to 0.10 and target power levels ranging from 0.7 to 0.9. **F:** Required number of samples under a varied number of spots per sample for case-control comparisons. **G:** Required number of samples under varied detectable differences for case-case comparisons. **H.** Cross-resolution comparison of SVF effect sizes and p-values at 8 μm and 16 μm. **I.** Comparison of empirical and theoretical power on Xenium and Visium HD samples. The shaded area represents the inter-quartile range of empirical power.

#### Individual-sample power analysis

We first examined the relationship between spatial resolution, effect size threshold, and statistical power to guide practical detection of SVFs within individual samples across platforms. The estimated power increased rapidly within a certain range of spatial measurements, depending on the pre-defined effect size thresholds (Fig. 2B). At a significance level of 0.05, 80% power was achieved with 424 spots for moderate effects and 137 spots for large effects. These spot counts are achievable in all Xenium samples and in the majority of Visium samples. In contrast, detecting small spatial effects required substantially higher resolution (4,620 spots), achievable in all Xenium samples but rarely in Visium (<2%). These results indicate that moderate effect sizes are a practical target for Visium-based kidney analyses, whereas Xenium enables detection of finer spatial structure.

Importantly, increasing the effect size threshold improves statistical power but reduces the proportion of SVF candidates (defined by p-values) that exceed the threshold and are therefore detectable. This creates a fundamental tradeoff between sensitivity to biological signals and feasibility of detection under realistic sampling constraints. To identify a balanced point, we jointly evaluated the median proportion of SVFs per sample exceeding each threshold and the proportion of samples with sufficient spatial resolution to achieve 80% power. The intersection of these two curves defines the optimal effect size threshold, which was approximately 0.06 for Visium and 0.01 for Xenium, consistent with their differences in spatial resolution and sensitivity for detecting small effects.

#### Group-level design

While individual-sample power determines whether spatial patterns can be detected within a sample, many biological studies ultimately require comparisons across conditions to distinguish the SVFs associated with disease and to ensure the robustness of the biological findings. We therefore next evaluated power for group-level analyses, including both case-control comparisons and case-case comparisons, between four conditions (HRT: healthy reference tissue, DKD: diabetic kidney disease, AKI: acute kidney injury, CKD: chronic kidney disease). Power curves were generated across sample sizes, and one-way sensitivity analyses assessed the impact of involved parameters (Fig. 2D, E).

For case-control analyses that test whether a spatial pattern is present in one group but absent in another, Xenium consistently achieved higher power than Visium, particularly for patterns with small effect sizes arising from limited spatial extent or high noise. This advantage reflects the greater sensitivity of Xenium for detecting spatial patterns within individual samples. Owing to its higher spatial resolution, Xenium also produced more stable cost estimates across parameter settings, whereas Visium was more sensitive to the selected effect-size threshold and the number of spots per sample. Increasing spot counts substantially reduced the required sample size for Visium; for moderate effects, the number of samples per group decreased from approximately 100 to 10 as spot counts increased from 100 to 300 (Fig. 2F). Under typical study conditions, 10 or fewer Visium samples per group were sufficient to detect moderate spatial patterns, whereas small effects were generally detectable only with Xenium, requiring as few as two samples per group when 10,000 cells were available per sample.

For case-case analyses that compare the strength of spatial dependency between groups, Visium showed slightly higher power than Xenium while profiling substantially more features. In both platforms, the target difference in spatial dependency strength was the primary determinant of sample size and cost. Assuming 80% power and a significance level of 0.05, detecting a difference of 0.10 in spatial effect size required approximately 11 Visium or 15 Xenium samples per group (Fig. 2G). Detecting smaller differences substantially increased the required sample size, reaching approximately 41 Visium or 57 Xenium samples per group for a difference of 0.05.

#### Dropout correction for cross-resolution comparisons

The derived effect sizes can be harmonized across spatial resolutions by modeling technical dropout, enabling consistent and comparable detection of spatial signals across platforms. Applying STORM to Visium HD datasets at 2 µm, 8 µm, and 16 µm, we identified SVFs and restricted analysis to the top 25% by effect size to ensure robustness. Dropout rates were estimated using the median effect size ratios between resolutions (detailed in Methods). We then applied these estimates to an independent dataset and corrected effect sizes derived from the 16 µm data. The corrected effect sizes showed strong concordance with those observed at 8 µm, whereas uncorrected effect sizes exhibited systematic deviation (Fig. 2H). Notably, SVFs identified by current methods relying solely on p-values, which showed poor agreement across resolutions due to their sensitivity to sample size, sampling density, and platform-specific effects. To further validate STORM’s statistical power estimates in real-world applications, we compared theoretical and empirical power using the IU01 sample profiled by two sequencing platforms (Visium HD and Xenium, Fig. 2I). Genes with strong evidence of spatial organization (adjusted p-value < 0.001) were treated as benchmarks. For each dataset, spots were randomly split into pilot and validation sets of equal size. Theoretical power was estimated from the pilot data at a significance level of 0.05, whereas empirical power was calculated as the true positive rate in the validation set. Different sample sizes were evaluated through random spot subsampling. Across a broad range of spot counts (100–5,000), theoretical power closely matched empirical power for both platforms, supporting the accuracy of STORM’s power estimation framework.

### STORM Identifies Inflammation Severity in Kidney Disease

We evaluated STORM’s capacity to identify disease-associated spatial organization in human kidney tissue and to quantify molecular patterns associated with DKD severity using data from the KPMP (Kidney Precision Medicine Project) atlas^29,31^. On the 153 Visium samples from four conditions: healthy reference tissue (HRT), diabetic kidney disease (DKD), acute kidney injury (AKI), and hypertensive chronic kidney disease (H-CKD), we first identified sample-level SVFs using a stringent threshold of adjusted p-value<0.01 and effect size<0.06. We then retained SVFs that were recurrent in more than 5% samples within each condition after merging H-CKD and DKD as chronic kidney disease (CKD). The majority of SVFs (3,901) were shared across all three conditions, likely association with kidney morphology. Meanwhile, smaller subsets of SVFs represented AKI-specific, CKD-specific, HRT-specific, and disease-shared SVF patterns (Fig. 3A).

**Fig 3.**
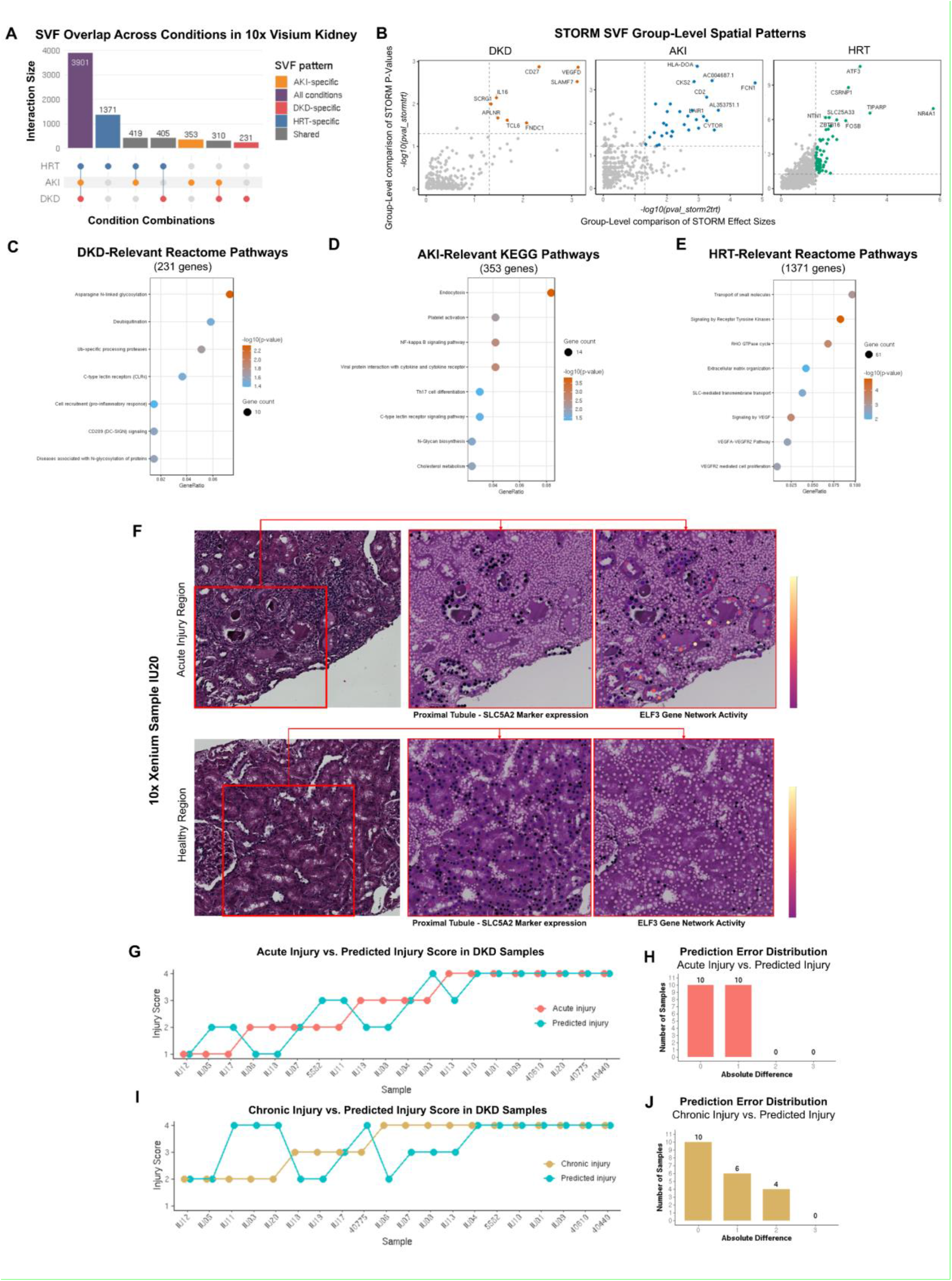
STORM identified disease- and injury-associated spatial patterns in kidney disease: **A:** UpSet plot showing overlaps of spatially variable features across HRT, AKI, and DKD conditions in 10x Visium kidney samples. **B:** Group-level STORM analysis comparing SVF recurrence and spatial effect-size differences across DKD, AKI, and HRT. Each point represents a feature, with the x-axis showing −log10 P value from between-condition effect size comparison and the y-axis showing −log10 P value from group-level significance comparison. Colored points indicate significant features by both group-level tests, and labeled genes highlight representative condition-associated spatial candidates. **C-E:** Pathway enrichment analysis of condition-associated SVFs prioritized from group-level STORM testing **F:** Representative 10x Xenium DKD sample regions showing H&E histology, proximal tubule marker expression, and ELF3 gene network activity in acute injury and healthy regions. **G, H:** Comparison of STORM-derived predicted acute injury scores with expert-assigned acute injury scores across DKD Xenium samples, with the corresponding absolute prediction error distribution. **I, J**, Comparison of STORM-derived predicted chronic injury scores with expert-assigned chronic injury scores across DKD Xenium samples, with the corresponding absolute prediction error distribution.

We next performed group-level tests on these condition-specific SVFs to assess whether spatial patterns were present within each condition and whether their spatial dependence differed across conditions, which reflects a disease-specific biological process rather than sample-specific variation. The overlap between these two group-level tests was identified for the most robust condition-associated spatial candidates (Fig. 3B), followed by a pathway enrichment analysis (Fig. 3C). CKD-associated SVFs were enriched for pathways related to proteostasis and immune remodeling, including *Deubiquitination, Asparagine N-linked glycosylation, and C-type lectin receptor signaling* (Fig. 3C and Supplementary Fig. 7A), consistent with protein-processing stress and innate immune activation in diabetic kidney injury^32,33^. AKI-associated SVFs showed a distinct acute injury profile, with enrichment of *Endocytosis, NF-κB signaling, and Th17 cell differentiation* (Fig. 3D and Supplementary Fig. 7B), supporting inflammatory signaling and immune activation responses^34–36^. In contrast, HRT-associated SVFs were enriched for pathways reflecting preserved tissue architecture and homeostatic kidney function, including *Transport of small molecules, Signaling by VEGF, and Extracellular matrix organization* (Fig. 3E and Supplementary Fig. 7C). The absence of these SVGs in disease-specific suggests that disease might be accompanied by loss of normal vascular-developmental and metabolic spatial organization, consistent with known roles of renal microvascular rarefaction, HOX-mediated kidney patterning, and lipid metabolism dysregulation in kidney disease^37–39^. To further visualize these condition-associated programs in tissue context, we mapped representative pathway expression scores onto Visium biopsy sections from DKD, AKI, and HRT samples (Supplementary Figs. 8-10). Together, these enrichment results indicate that STORM-derived SVFs associate with biologically distinct disease-associated spatial programs, distinguishing DKD-associated proteostasis and immune remodeling, AKI-associated inflammatory injury responses, and HRT-associated epithelial, vascular, and structural organization.

#### Xenium Kidney DKD Severity Analysis

To demonstrate the biological utility of spatial dependency strength, we examined whether STORM-derived effect sizes could quantify kidney injury severity in 20 10x Xenium DKD samples with expert nephropathologist annotations. We focused on the ELF3 gene network, an injury-associated transcriptional program active in the kidney cortex that includes epithelial injury markers such as KLF6, KLF10, ITGB3, and TPM1^40^. Although all six ELF3 network genes were significant (adjusted p < 0.01) in every sample, only KLF6 and KLF10 showed effect sizes below the 0.06 threshold in a subset of samples, and these reductions were associated with disease severity.

An illustrative example was DKD sample IU20, which contained a severely injured region of tubulointerstitium characterized by dilated proximal tubules, intraluminal casts, epithelial simplification, and peritubular immune cell infiltration, alongside a relatively preserved region with intact brush borders and minimal inflammation (Fig. 3F). Mapping ELF3 network activity onto tissue histology revealed strong, spatially localized activation within injured tubular regions, whereas healthier areas exhibited weaker and more diffuse activity which is consistent with previous results^40^. These findings demonstrate that STORM effect sizes capture localized molecular organization associated with histologically visible injury.

To evaluate the clinical relevance of these spatial features, we assessed whether ELF3 network effect sizes could predict sample-level injury severity. Acute and chronic injury scores were categorized into ordinal bins based on the proportion of injured tubulointerstitial tissue (Supplementary Table 6). Predicted injury scores were derived from the median STORM effect sizes of ELF3 network genes and grouped into four severity categories for each of acute and chronic injury. For acute injury, predictions were exact in 10 of 20 samples and differed by only one category in the remaining samples (Fig. 3G,H). Predicted scores were strongly correlated with acute injury severity (Spearman’s ρ = 0.82, p < 0.0001), indicating that STORM-derived spatial effect sizes capture the ordinal progression of acute tubular injury. In contrast, chronic injury predictions were less accurate, with exact agreement in 10 samples, one-category differences in 6 samples, and two-category differences in 4 samples (Fig. 3I, J). Correspondingly, the association with chronic injury severity was weak and non-significant (ρ = 0.22, p = 0.350). This divergence suggests that STORM-derived predictions of ELF3 spatial organizations can reflect active epithelial injury and acute tubular damage.

### STORM Identifies Unique Organelle Organization in Mouse Liver with Different Diets

In addition to traditional cellular-level omics, STORM extends spatial inference beyond tissue-level molecular profiling to subcellular organization in scale. We applied STORM to image-derived organelle data generated from high-resolution confocal microscopy of mouse liver under three dietary conditions: control (CNT), starvation (STV), and Western diet (WD) (Fig. 4A).

**Fig 4.**
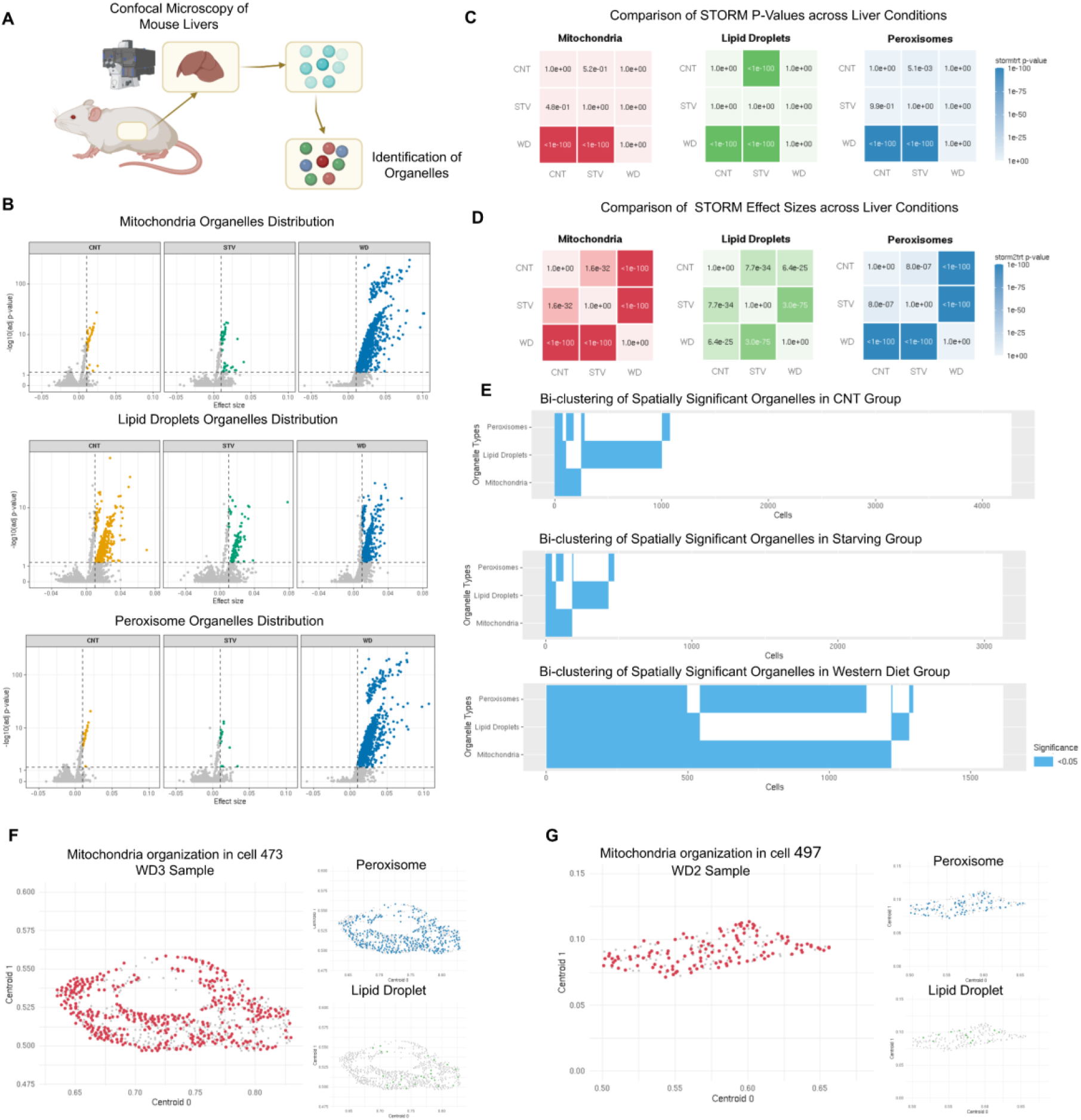
STORM reveals diet-associated organelle spatial organization in mouse liver: **A:** Overview of the experimental workflow. Mouse liver tissue is imaged using confocal microscopy, followed by segmentation and identification of mitochondria, lipid droplets, and peroxisomes at single-cell resolution. Organelle centroids are extracted and used as spatial coordinates for downstream STORM analysis. **B:** Scatter plots of STORM-derived effect size versus statistical significance for organelles across conditions. Each point represents a single cell. Dashed vertical lines denote STORM effect-size thresholds corresponding to small effect size (0.01), while the horizontal dashed line denotes the nominal significance threshold of p = 0.05. **C:** Group-level significance comparison heatmaps comparing the proportion of cells with significant spatial organelle organization across CNT, STV, and WD conditions for each organelle class. **D:** Pairwise between-condition effect size comparison heatmaps comparing STORM-derived organelle effect sizes across CNT, STV, and WD conditions for each organelle class. **E:** Bi-clustering heatmaps showing the presence of spatially significant organelles (p < 0.05) across cells for each condition. **F-G:** Representative single-cell spatial organization of organelles in WD samples. Spatial coordinates of mitochondria (red), lipid droplets (green), and peroxisomes (blue) are shown for two individual cells, illustrating structured and coordinated intracellular organization.

Mitochondria, lipid droplets, and peroxisomes were fluorescently labeled, segmented at single-cell resolution, and represented by organelle centroids coordinates for spatial analysis. Organelle classes and subtypes were encoded as binary features within each cell. After excluding outlier cells containing more than 10,000 organelles, the final dataset included 4,144 CNT cells containing 8,459,255 organelles, 3,022 STV cells containing 5,587,219 organelles, and 1,532 WD cells containing 4,414,345 organelles (Supplementary Fig. 11).

Across all three organelle classes, cells from the WD group exhibited substantially stronger intracellular spatial organization than those from the CNT and STV groups. Volcano plots showed that CNT and STV cells were concentrated at low effect sizes, whereas WD cells were enriched for spatially organized organelles, particularly mitochondria and peroxisomes (Fig. 4B). For mitochondria, 79% of WD cells showed effect sizes greater than 0.01 and 1% exceeded 0.06, with effect sizes reaching approximately 0.10. Peroxisomes displayed a similar pattern, with 72% of WD cells exceeding 0.01 and 0.7% approaching 0.10. Lipid droplets also exhibited increased spatial dependence in WD cells, although with smaller effect sizes. These results suggested that WD induces a broad increase in intracellular spatial structuring across multiple metabolic organelles, with the strongest effects observed for mitochondria and peroxisomes.

We next assessed group-level differences in both the prevalence and strength of organelle spatial organization. WD cells showed a significantly higher prevalence of spatial organization than CNT and STV cells for mitochondria, peroxisomes, and lipid droplets (all p < 0.0001; Fig. 4C). Effect-size comparisons similarly revealed stronger spatial dependence in WD cells across all three organelle classes (all p < 0.0001; Fig. 4D). STV cells also exhibited greater prevalence than CNT cells for peroxisomes (p = 0.0051) and lipid droplets (p < 0.0001), as well as stronger spatial dependence for all three organelles (all p < 0.0001). These results demonstrate that WD increases both the frequency of cells with significant organelle spatial patterns and the magnitude of spatial dependence.

To determine whether these statistical patterns reflect broad changes across the single-cell population, we constructed binary cell-by-organelle matrices indicating whether each organelle class was spatially significant within each cell, followed by bi-clustering to visualize condition-specific patterns. CNT and STV samples showed more restricted spatial organization, with significant organelle patterns appearing in a smaller subset of cells. In contrast, WD samples showed a broader and more coordinated pattern of spatially significant mitochondria, lipid droplets, and peroxisomes across the cell population (Fig. 4E).

Finally, representative WD cells illustrate the spatial configurations underlying the STORM results (Fig. 4F–G). In WD3 cell 473, mitochondria formed more structured, elongated patterns that were statistically significant by STORM while lipid droplets and peroxisomes did not show similar organization and did not reach statistical significance. Similarly, in WD2 cell 497, mitochondria again showed significant spatial organization, whereas lipid droplets and peroxisomes showed no significant spatial dependence. These single-cell examples are consistent with the group-level results, highlighting mitochondria as a prominent organelle class undergoing WD-associated spatial reorganization, while also demonstrating cell-to-cell heterogeneity in the spatial arrangement of other organelles. Collectively, this case study demonstrates that STORM detects and quantifies biologically meaningful subcellular spatial organization from imaging-derived organelle data, extending its utility beyond conventional spatial omics and enabling rigorous comparison of intracellular architecture across experimental conditions.

## Discussion

In this work, we present STORM, a principled statistical framework that unifies spatial pattern detection, quantitative effect size estimation group-level inference, and power analysis for spatial multi-omics studies. Unlike existing approaches that primarily focus on identifying SVFs within individual samples, STORM provides an integrated framework for discovery, interpretation, comparison, and experimental design. By linking spatial hypothesis testing with interpretable measures of spatial organization and prospective power calculations, STORM establishes a generalizable statistical foundation for both analyzing and planning spatial multi-omics experiments.

A key feature of STORM is its support for prospective experimental design. Most spatial analysis methods are intended for post hoc analysis and provide little guidance for study planning. STORM incorporates analytical power estimation that jointly considers spatial resolution, biological replication, and detectable effect sizes. Extensive simulations and benchmarking demonstrate that STORM provides well-calibrated statistical inference and strong detection performance across diverse scenarios, while scaling efficiently to high-resolution data. The kidney case study further illustrates the trade-off between spatial resolution and biological replication, showing that increased within-sample resolution can substantially reduce the number of samples required to achieve adequate statistical power. Such guidance is particularly valuable given the cost and logistical challenges of modern spatial multi-omics studies.

STORM also extends spatial inference to the group level beyond single-sample analyses. Many biological questions center on how tissue organization changes across disease states, treatments, or developmental conditions. However, most existing spatial analysis tools are designed primarily for within-sample inference and do not provide formal statistical frameworks for comparing spatial organization across groups. STORM addresses this gap by enabling both condition-specific detection of spatial patterns and direct comparisons of spatial effect sizes between biological groups. This capability shifts spatial multi-omics analyses from descriptive mapping toward rigorous statistical evaluation of spatial tissue remodeling.

The biological and translational utility of STORM was demonstrated by investigating kidney disease. Guided by the statistical power analysis, group-level analyses on Visium samples identified reproducible disease-associated spatial programs that differentiated chronic kidney disease, acute kidney injury, and healthy reference tissues. In Xenium data, sample-level spatial effect sizes of ELF3 activity provided quantitative measures of injury-associated spatial organization that closely tracked histopathological assessments of acute tubular injury. Together, these analyses highlight complementary strengths of the framework using cohort-level inference to identify disease-associated spatial programs, whereas sample-level effect sizes provide interpretable quantitative signatures that can be linked to disease severity and tissue pathology. The general formulation of STORM extends beyond transcriptomics. Because the framework requires only spatial coordinates and quantitative feature measurements, it can be applied to diverse spatial modalities, including proteomics, epigenomics, and image-derived phenotypes. This flexibility was demonstrated in the organelle analysis, where STORM was utilized to quantify spatial organization among millions of intracellular structures and identify condition-specific reorganization of organelles in mouse liver tissue.

Several limitations should be considered. First, STORM quantifies spatial dependence with a neighborhood approach, and its performance may be affected when spatial patterns are highly localized. Although simulations demonstrated robust performance across a range of neighborhood sizes, overly large neighborhoods may dilute signals arising from small, isolated spatial domains. Second, group-level inference currently assumes independence among biological samples. While inference is based on sample-level intermediate measures and is therefore expected to be robust to batch effects or within-subject heterogeneity, residual dependencies may underestimate variance and thereby increase the error rate. Future work will focus on extending STORM with hierarchical modeling frameworks that more explicitly account for complex sample-to-sample variability while preserving its utility for power analysis, sample size determination, and experimental design.

## Methods

### Derivation of Test Statistics

For a spatially resolved multi-omics sample with *M* spots and *N* features, the *D*-dimensional coordinates of spot *i* are denoted as a *D*-vector ***s***_*i*_, where *i* = 1, …, *M*, and the corresponding coordinates matrix of all spots are denoted as a *M* × *D* matrix 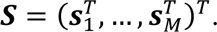 The expression of a feature *j* is denoted as ***x***_***j***_ = (*x*_1*j*_, …, *x*_*Mj*_), where *j* = 1, …, *N*, and the expression at the spot *i* is denoted as *x*_*ij*_. For each spot in the space, the *K* nearest neighboring spots are determined, denoted as a binary M-vector 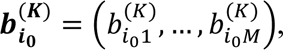 where 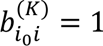 if spot *i* is within the *K* nearest neighboring spots of spot *i*_0_. For each feature *j* at each spot *i*, the sum of the expressions of all neighboring spots, 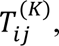 can be calculated as 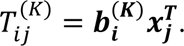 The *<u>null hypothesis</u>* is that a feature is not a spatially variable feature, i.e., the expression of a feature does not rely on the spatial location of spots (***x***_***j***_ ⊥ ***S***).

Denote the distribution of feature *j* as *F*_*j*_(*x*) with mean and variance as *μ*_*jx*_ and 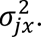 Under null hypothesis, the squared difference between a scaled expression and the sum of its neighbors, 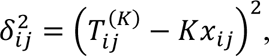 follows a normal distribution as follows:

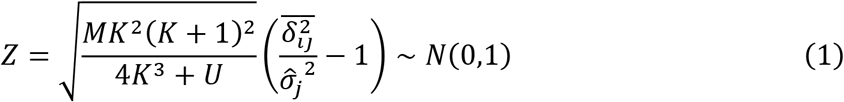

where *U* is a constant that depends on *M* and 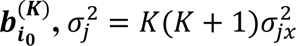 denotes the variance of *δ*_*ij*_ (Supplementary Note 1-4).

The null hypothesis is equivalent to

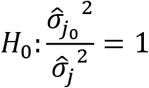

The sample variance 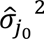 can be approximated with 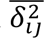 for a large M since *E*[*δ*_*ij*_] = 0. For a two-sided test, the probability of rejecting null hypothesis can be obtained from Equation (1) as

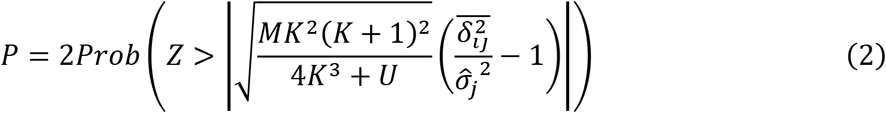

To measure the proportion of variance that can be explained by the dependency of expressions, the effect size for this test can be calculated as follows:

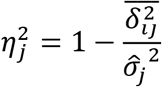

For a feature whose expressions rely on the spatial location of cells, the variance of *δ*_*ij*_ would be 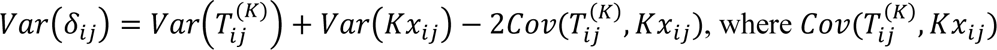 is the covariance between 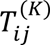 and *Kx*_*ij*_. Therefore, 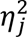 can be rewritten as

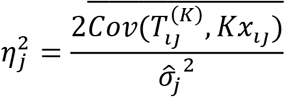

which represents the proportion of variance that can be explained by spatial dependency. The value of 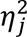 will be zero when there is no spatial autocorrelation, and it tends to be large when a high spatial autocorrelation presents. It is independent of the number of spots; however, its theoretical upper limit is affected by the dropout rate. Assuming the dropout event occurs independently across the spots and is independent of the expressions, the observed effect size after dropout can be derived as

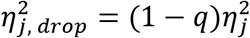

where *q* is the dropout rate (Supplementary Note 5).

Similarly, we also consider an additional noise term in addition to the inherent biological variability, ***ϵ***_***j***_, with mean zero and variance 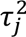 in the expressions of each feature to account for the technically introduced variations. The observed effect size after considering noise can be derived as

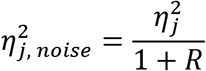

where 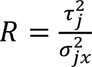 represents the signal-noise ratio (Supplementary Note 6).

These underestimated effect size reduces the power of the statistical test, making it less likely to reject the null hypothesis even when it is false. That is, the dropout events and technical noise will increase the risk of Type II errors (false negatives), potentially causing true spatial patterns to go undetected. As a result, the test is more likely to fail to identify the spatial variations with increased dropout rate or noise levels. To mitigate the underestimation of the effect magnitude, an adjusted threshold for the effect size should be applied, as outlined in equations above. When the expression of a feature can be approximated by a normal distribution (e.g., Poisson or negative binomial distribution), the test statistics can be approximated as follows (Supplementary Note 7-8):

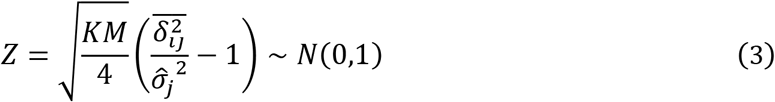

This approximation provides a computationally efficient framework and enables more straightforward statistical power analysis. Both the non-parametric test and its parametric approximation can be extended to the cell-type-related or cell-type-specific SVF detection (Supplementary Note 9).

### Sample Size Determination

Denote the expected proportion of variance that can be explained by the dependency of expressions under the alternative hypothesis as 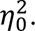 The statistical power (1 − *β*) can be determined as

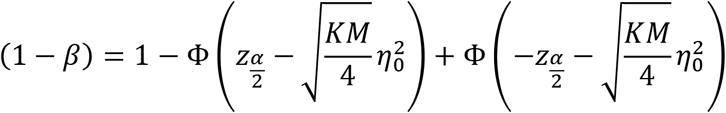

where Φ(⋅) is the cumulative distribution function (CDF) of the standard normal distribution. The minimum sample size can be derived from the equation above as

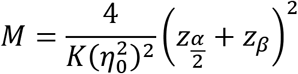

Considering the effect of additional noise, the minimum sample size should be adjusted to

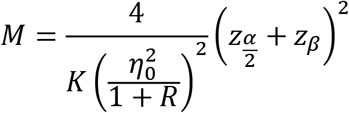

where 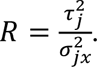To ensure the weakly dependency of *K*^2^-dependent sequence ***δ_j_***, *K* should satisfy *K*^2^ < *M*. Therefore, the minimum sample size could be determined as:

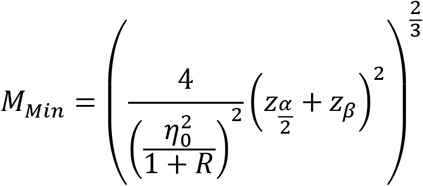

Similarly, the statistical power for the generalized test statistics can be derived as

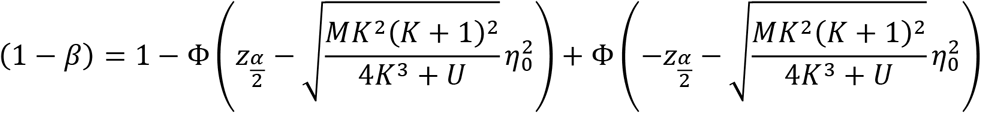

and the sample size can be obtained by solving the equation above. The term of *U* can be estimated from a bootstrap method in simulations.

### Comparisons Between Two Case Groups

To compare the differences in the magnitudes of spatial patterns between two case groups, a random effect sampling model can be applied using the effect sizes of the spatial patterns.

Denote the effect size of the spatial pattern on sample *l* as 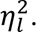 The effect size on sample *l* follows a normal distribution with variance of *ϕ*_*l*_ since

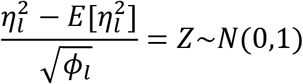

based on Equation (4) or (7), where

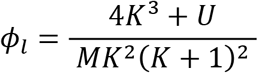

or

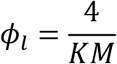

Assuming the between-sample variation follows a normal distribution with variance of *ψ*_*G*_ for group *G* and denoting the group means of effect sizes as *ξ*_*G*_ the distribution of effect sizes can be derived as

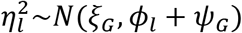

where *G* = 1 or 2 representing the case group index. The inverse-variance weights for each sample can be calculated as

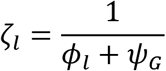

The inverse-variance weighted mean in each group can be then calculated as

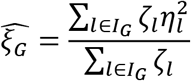

where *I*_*G*_ denotes the set of indicators for group *G*. The estimated variance of 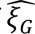 derived as 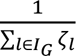 since the samples are independent. The null hypothesis of this test is

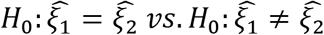

which is equivalent to

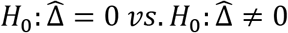

Since the samples are independent, the variance of Δ̂ can be calculated as 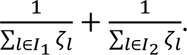

Therefore, the test statistics can be derived as

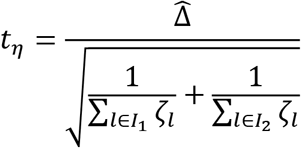

which approximately follows a t-distribution when sample sizes are reasonably large (for example, both are greater than 5^41^. Specifically, when designing an experiment where samples are collected using the same sequencing platform with a comparable number of spots, and the tissues have similar shapes, the differences in within-sample variation (*ϕ*_*l*_) are expected to be small with the same number of neighbors (*K*) during the analysis, and thus the test statistics can be approximated by

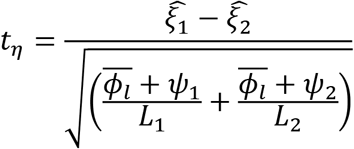

with the degree of freedom estimated using the Satterthwaite method

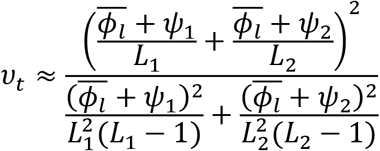

The minimum sample size to ensure a statistical power of 1 − *β*_*G*_ can be solved iteratively as the t-quantiles depend on the degree of freedom which also depends on the sample sizes. When the sample sizes are sufficiently large such that, for example, the estimated Satterthwaite degree of freedom is greater than 30^42^, the distribution of test statistics *t*_*η*_ can be normally approximated and the minimum sample sizes can be determined as

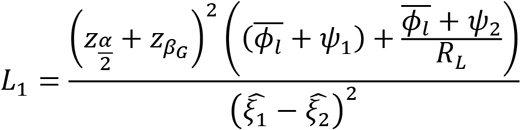

and

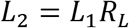

where 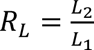 denotes the ratio of allocation.

### Comparisons Between Control and Case Groups

When comparing the significance of an SVF between control and case groups, a two-sample one-sided proportion test can be used as an alternative by comparing the proportion of samples that reject the individual test in the two groups. This approach is applicable if the control samples truly have no spatial pattern, while the case samples exhibit spatial patterns. Specifically, let *p*_*t*_ and *p*_*c*_ denote the probabilities that a case and control sample, respectively, is declared significant. The null hypothesis of this test is

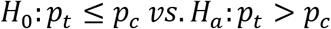

For the equal sample scenario, denote the number of biological samples in the control and case groups as *L*. When the significant level and statistical power for each single test have been determined, the case effect at the group level can be tested using a two-proportion z-test:

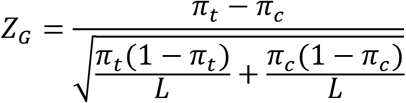

which follows a standard normal distribution under null hypothesis that the significance of a SVF between control and case groups are identical, where *π*_*c*_ = *α* is the probability that a control sample is significant (Type I error) and *π*_*t*_ refers to the probability that a case sample is significant (statistical power of each single test) which can be written as

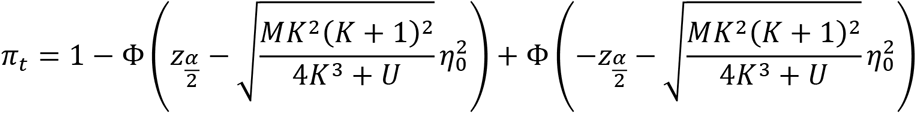

The number of biological samples can be determined as

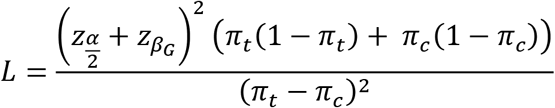

where *β*_*G*_ refers to the statistical power for the multi-sample tests.

In case of unequal biological samples in the control and case groups, the sample size can be determined as

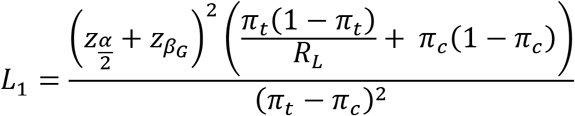

and

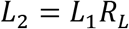

where *L*_1_ and *L*_2_ are the number of biological samples in the control and case groups and 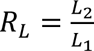 denotes the ratio of allocation.

### Assessing Type I Error Under the Null Hypothesis

Type I error was evaluated using two simulation settings that varied in spatial resolution and sparsity. The low-resolution scenario consisted of 900 spots with 90% non-zero expressions, whereas the high-resolution scenario included one million spots with 0.5% non-zero expressions. For each scenario, expressions for 10,000 features without any spatial structure were generated from a scaled beta distribution, [10 × Beta(1,5)], where ⌊x⌋ denotes the greatest integer less than or equal to x. The calculated p-values for each method were compared with their theoretical uniform distribution using a quantile-quantile plot. The Kolmogorov–Smirnov (K–S) distance between the empirical p-value distribution and the theoretical Uniform distribution was calculated for the quantitative comparisons, following prior work. A smaller K–S distance indicates closer agreement with the expected uniform distribution, reflecting better calibration and more accurate control of Type I error. To evaluate Type I error in realistic applications, we further performed analyses using permuted datasets derived from nine sources spanning diverse spatial masks, histological types, sequencing platforms, and sequencing depths. For each dataset, the spatial coordinates were randomly permuted to eliminate genuine spatial structure while preserving expression characteristics. Statistical calibration was assessed by computing the K–S distance between the empirical p-value distribution and the theoretical Uniform distribution from 0 to 1.

### Evaluating Type II Error Under the Alternative Hypothesis

To assess Type II error under the alternative hypothesis, we adapted a previously established framework to generate SVFs (Supplementary Note 10). The continuous structures Curved Cell Strand defined in previous work were included which typically had the smallest effect sizes compared to the aggregated cell clusters. We simulated 1000 coordinates through a 3D random-point-pattern Poisson process. The patterns were formed by generating center points through a random walk with a fixed step length of 2 units. The monotonicity of movement directions was controlled to be monotonic in two directions. The expression of SVFs was simulated by distinguishing between marked and non-marked cells. For marked cells within the pattern, we randomly sampled feature expression values from the upper quantile of the feature expression distribution in seqFISH data. In contrast, non-marked cells and those outside the pattern were assigned feature expressions randomly from the entire seqFISH dataset. The effect size of the spatial pattern was controlled by varying the pattern radius between 1.1 and 1.8, ensuring that the resulting statistical power covered a broad range from 0 to 1. For each fixed radius, 100,000 SVFs were generated per dataset, and this procedure was repeated 20 times. Observed statistical power was calculated as the proportion of p-values below 0.05 from the approximate test, while theoretical power was estimated using the observed number of spots and average effect sizes at a 0.05 significance level. The difference between observed and estimated statistical power was tested as a fixed effect using a generalized linear mixed model, with degrees of freedom calculated via the Satterthwaite approximation.

### Benchmarking Accuracy in Detecting Spatially Variable Features

To benchmark the detection accuracy of STORM and ensure fair comparisons with other methods, we utilized simulation datasets from a previously published benchmarking study^25^. Specifically, nine datasets were generated using scDesign3, with source data spanning nine distinct spatial masks, five tissue histology types, two spatial transcriptomics platforms, and varying sequencing depths. Each method was applied to the simulated datasets, and performance was quantified using the AUC and the AUPRC.

We further conducted simulations following the established framework in previous studies to evaluate the detection capacities in three-dimensional space (**Supplementary Note 6**). Four three-dimensional spatial patterns were considered, including three continuous patterns (Curved Cell Strand, Tissue Layer, and Irregular Cell Aggregate) and one discrete pattern (Isolated Cell Nodules). Statistical power was computed as the true positive rate across a range of false discovery rate thresholds to account for method-specific p-value distributions. Precision–recall curves were constructed for each condition and replicate, and AUPRC was calculated for quantitative comparisons.

### Validation of group-level inference

Group-level inference was evaluated using the three-dimensional simulated datasets described above. For each spatial pattern, 1,000 SVFs were generated under varying pattern sizes, signal strengths, and noise levels. Spatial dependence on each feature was quantified using the STORM effect size. Two types of group comparisons were performed. For case–control comparisons, features with spatial patterns were compared against features without spatial structure. For case–case comparisons, two groups of SVFs with differing spatial dependence strengths were compared. All analyses were conducted using both STORM and its approximation.

To assess statistical power at small sample sizes, feature subsets were randomly sampled to form groups of 3–20 samples. For each sample size, group-level tests were repeated 1,000 times. Empirical power was calculated as the proportion of tests with p-values below 0.05. Theoretical power was estimated based on the corresponding sample size and average effect size for the approximation of STORM. Type I error was evaluated under null scenarios for both comparison types. For case–case comparisons, two groups of 100 SVFs with identical spatial effect sizes were randomly sampled. For case–control comparisons, two groups of 100 non-spatial features were sampled. Each scenario was repeated 50 times, and the resulting p-values were compared with the expected uniform distribution to assess calibration.

### Evaluating Computational Scalability

We evaluated computational efficiency by comparing runtime and peak memory usage across spatial analysis methods under low- and high-resolution scenarios by randomly sampling from two published datasets. In the low-resolution setting, the stxbrain dataset from the Seurat package^43^ was used to benchmark STORM, STORM (approximated), SPARK-X, scBSP, SOMDE, SPARK, nnSVG, SpatialDE, and Moran’s I. Scalability was assessed under two sub-scenarios: (i) fixing 20,000 features while sampling the number of spots from 500 to 5,000 (500, 1,000, and increments of 1,000), and (ii) fixing 3,000 spots while varying the number of features from 500 to 100,000 (500, 20,000, and increments of 20,000). For the high-resolution setting, we applied a similar framework to an HDST dataset, benchmarking STORM, STORM (approximated), SPARK-X, scBSP, and GPU-accelerated implementations of STORM and its approximation. Two sub-scenarios were considered: (i) fixing 20,000 features while varying spots from 10,000 to 1,000,000 (10,000, 100,000, and increments of 100,000), and (ii) fixing 200,000 spots while varying features from 500 to 100,000 (500, 10,000, and increments of 10,000). Runtime and peak memory usage were recorded in all cases to assess scalability with respect to spatial resolution and feature dimensionality. Executions exceeding 2 hours were terminated. All methods were executed and recorded on a workstation with a 2.00GHz AMD EPYC 7713 64-Core Processor.

### Experimental Design Study

Kidney spatial transcriptomics datasets generated using the Visium Spatial Gene Expression and Xenium In Situ platforms from 10x Genomics were used for the experimental design case study. The Visium dataset included 153 samples from four groups: healthy controls, acute kidney injury, diabetic kidney disease, and hypertensive chronic kidney disease. Hypertensive chronic kidney disease and diabetic kidney disease samples were combined to form the CKD group analysis. The Xenium dataset included 38 samples from three groups: healthy controls, diabetic kidney disease, and systemic lupus erythematosus. Each sample contained spatial coordinates and gene expression measurements across spatial locations (spots or cells), with varying spatial resolution across platforms. For each sample, p-values and spatial association statistics were obtained using the approximation of STORM. Features with zero expression across all spatial locations were excluded from the analysis.

SVF candidates were defined as features with adjusted p-values below, corrected by the Benjamini-Hochberg procedure for multiple testing corrections. The estimated spatial effect size represents the proportion of expression variance explained by spatial dependence. Effect size thresholds of 0.01, 0.06, and 0.14 corresponding to small, moderate, and large spatial effects were considered based on commonly used variance-explained guidelines^44^. Statistical power for detecting SVFs within individual samples and between groups was calculated using the analytical power function implemented for the approximation of STORM.

For the individual sample test, power curves were generated across ranges of effect sizes and spot counts to evaluate spatial resolution requirements for achieving 80% statistical power. The minimum detectable effect size required to achieve 80% power was determined based on the observed number of spatial locations for each sample. The proportion of SVF candidates with estimated effect sizes exceeding this threshold was then calculated to assess the detectability of spatial signals within each sample. For the group-level inference, power curves were generated for number of samples ranging from 2 to 20 per group under a significance level of 0.05 and target power of 0.8. The number of spots per biological sample was estimated using the median number of spots. Between-sample variance was estimated from the observed effect sizes of SVF candidates. For each condition group of Visium and Xenium, features were ranked by the frequency of being detected as SVFs across samples, and the top 30 shared features were selected. The variance of their estimated effect sizes across samples within the same condition was calculated, and the platform-level between-sample variance was summarized as the mean of these feature-specific variances.

Nine parameters affecting study design were further examined to account for the uncertainties in data and design. Four design parameters including target power, group level significance level, individual test significance level, and allocation ratio reflect typical planning choices. Five biological and economic parameters including effect size or effect size difference, number of spots for the individual test, between sample variance of effect sizes, number of spots per sample, and cost per sample capture biological and practical variability. Baseline values were set at 80 percent target power, 0.05 significance levels, equal group allocation, threshold for moderate effect and 0.10 for the detectable difference, median number of spots for the samples from Visium and Xenium, averaged between-sample variance from top 30 shared SVF candidates, $2,200 and $1,100 cost per Visium and Xenium sample based on previous studies^30^. Design parameters were varied within realistic ranges with power from 70 to 90 percent, significance levels from 0.01 to 0.10, and allocation ratio from 0.5 to 1.5. Biological and economic parameters were varied by 50 percent above and below the baseline to account for uncertainty. One-way sensitivity analysis was performed to assess the effect of each parameter on total cost estimates, and results were summarized using tornado plots.

### Estimation and Correction of Dropout

To enable comparison of effect sizes across resolutions, we evaluated technical dropout using Visium HD kidney datasets generated at 2 µm, 8 µm, and 16 µm resolution. The top 25% of SVFs ranked by effect size within each resolution were selected, and the effect sizes were extracted for a common set of top SVFs present across all three resolutions. As indicated in previous literature^45^, the fraction of molecules being captured can be modeled with a Poisson process *x*_*ij*_ ∼ *Poisson*(*λA*_*res*_), where *λ* represent the molecular density per unit area and *A*_*res*_ is the capture area. The fraction of zeros is thereby calculated as *q* = 1 − *e*^*λA_res_*^, which can be estimated by the effect size ratios. Assuming squared areas at 2 µm, 8 µm, and 16 µm resolution, we derived *λ* by quantifying medians of effect size ratios between resolutions and minimizing

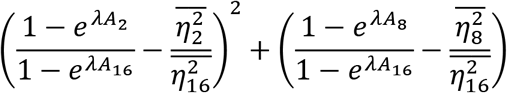

The dropout rate at each resolution was calculated accordingly.

To ensure generalizability, the estimated dropout parameters were applied to an independent Visium HD dataset. Effect sizes from 16 µm data were rescaled using the estimated dropout rates and compared with observed 8 µm effect sizes. For comparison, we also evaluated agreement using uncorrected effect sizes and p-values.

### Kidney Spatial Analysis

Kidney case study was performed using two complementary human kidney spatial transcriptomics datasets generated by 10x Genomics Visium and 10x Genomics Xenium platforms. The Visium cohort consisted of 153 kidney tissue sections spanning multiple disease and reference conditions, including 54 DKD, 43 AKI, 9 H-CKD, and 47 HRT samples, providing a large-scale cohort for evaluating spatial organization across kidney disease states^29^. The Xenium dataset was used as high-resolution validation and severity-stratification cohort focused on DKD tissue, comprising 20 DKD samples with matched histopathologic injury annotations^30^. For the 10x Visium analysis, each sample was processed independently using STORM to quantify spatial organization for each measured feature. For each tissue section, spot-level gene expression and spatial coordinates were used as input to STORM. These spot-level outputs were generated for each Visium sample and subsequently aggregated across samples according to disease condition.

Sample-level SVF candidates were defined using both statistical significance and effect-size criteria. Specifically, a feature was considered spatially variable in an individual sample if it had an adjusted p-value < 0.01 after the Benjamini–Hochberg procedure, and an effect size > 0.06. Features with missing adjusted p-values or missing effect sizes were not considered SVF candidates for that sample. To define reproducible condition-level SVFs, we used a condition-specific recurrence threshold based on the number of samples in each group. A feature was required to be spatially variable in more than 5% samples within each condition to be considered a group-level SVF for that condition. This recurrence criterion was used to control features that appeared spatially significant in only a small number of samples within a cohort. To identify condition-restricted spatial programs, group-level SVFs were further filtered by their recurrence across non-target conditions. A feature was defined as condition-associated if it exceeded the recurrence threshold in the target condition and remained at or below the corresponding recurrence threshold in each non-target condition. This procedure produced condition-associated SVF sets for CKD, AKI, and HRT, and enabled the identification of SVFs shared across multiple condition combinations. For chronic kidney disease pathway interpretation, DKD- and H-CKD-associated SVFs were considered jointly where appropriate to capture broader chronic disease-associated spatial programs, while group-level statistical comparisons were performed using the condition-specific SVF definitions.

Group-level comparisons were then performed to determine whether spatial patterns were specific to a condition and whether the strength of spatial dependence differed across conditions. Pathway enrichment analysis was performed on condition-associated SVF sets using KEGG and Reactome databases. Gene symbols were converted to Entrez identifiers using ‘*org.Hs.eg.db*’ prior to enrichment testing, resulting in some gene dropoff. KEGG enrichment was performed using ‘*enrichKEGG’*, and Reactome enrichment was performed using ‘*enrichPathway’* with readable gene symbols returned for interpretation. Enrichment analyses were run with permissive pathway-size parameters to retain kidney-relevant pathways represented by small candidate sets, and pathway-level results were prioritized based on biological relevance to kidney disease, nominal enrichment significance, gene ratio, and interpretability within the DKD, AKI, or HRT context. For visualization of pathway activity in tissue space, representative pathway gene sets were scored in Visium samples using ‘*scanpy.tl.score_genes’*, negative pathway scores were set to zero for visualization, and pathway scores were mapped back onto Seurat spatial objects as metadata for overlay on Visium histology images.

For the Xenium DKD analysis, the cohort consisted of 20 DKD tissue sections with matched nephropathology annotations for acute and chronic injury. STORM was run independently for each sample, and feature-level p-values and effect sizes were retained for downstream severity modeling. The analysis focused on transcriptional programs associated with kidney injury and proximal tubular dysfunction identified by ELF3 activity, including marker genes and gene-network activity scores selected based on their relevance to DKD-associated epithelial injury.

To connect spatial molecular organization with histopathology, STORM effect sizes were summarized across injury-associated gene sets for each Xenium sample. The six genes used to quantify ELF3 activity were ELF3, KLF6, KLF10, VCAM1, TPM1, and PROM1^40^. For each sample, effect sizes from selected injury-relevant features were aggregated into a predicted injury score, which was then compared with expert-assigned acute and chronic injury scores. Acute and chronic injury were evaluated separately to determine whether spatial organization of molecular injury programs captured distinct histopathologic processes. Each sample was scored on an ordinal scale from 1 to 4 to summarize the proportion of the cortical tubulointerstitium affected by acute or chronic injury (1: ≤10% affected, 2: 11-25% affected, 3: 26-50% affected, 4: ≥51% affected). Acute injury was defined by the presence of dilated proximal tubules, intraluminal casts, epithelial simplification, and peritubular immune cell infiltration. Chronic injury was defined by the presence of interstitial fibrosis and tubular atrophy. Prediction accuracy was assessed by *Spearman*’s correlation and comparing the absolute difference between the STORM-derived predicted score and the corresponding nephropathology score for each sample. Error distributions were summarized across the 20 DKD samples to evaluate the degree of agreement between molecular spatial organization and histologic severity annotations.

Representative Xenium regions were visualized to illustrate how STORM-derived injury programs mapped onto tissue morphology. Using *SpatialData*, histology images were paired with spatial overlays of proximal tubule marker expression and ELF3 gene network activity in regions annotated as acutely injured or relatively healthy. Spatial overlays were used to qualitatively assess whether injury-associated transcriptional activity localized to morphologically abnormal regions, while preserving tissue context.

### Organelle Analysis

High-resolution spatial organelle data were obtained from mouse liver tissue using a confocal microscopy–based scOrganellomics workflow, which enables simultaneous visualization and quantification of mitochondria, peroxisomes, and lipid droplets at single-cell resolution within intact liver lobules. Liver sections from mice subjected to three dietary conditions: control (CNT), starvation (STV), and Western diet (WD). These sections were fluorescently labeled to mark organelles and cell boundaries, followed by volumetric imaging across multiple lobules per sample. A Cellpose-based segmentation pipeline was applied to identify hepatocyte boundaries and segment individual organelles, yielding spatial coordinates (centroids) and quantitative measurements for each organelle. Across all samples, this dataset comprised more than 4,000 hepatocytes and millions of organelles, capturing subcellular organization across physiologically relevant metabolic conditions. Additional details regarding data generation are referred to the Organellomics paper^46^.

Organelle centroids were treated as spatial coordinates, and organelle-specific features were aggregated at the cell level to construct feature matrices for each sample. Cells containing extreme organelle counts (>10,000 organelles) were excluded to mitigate segmentation artifacts and ensure comparability across conditions. After filtering, cells and organelles from the CNT, STV, and WD samples were retained for downstream analysis. For each cell and organelle class, STORM was applied to test for spatial dependence, generating both p-values and spatial effect size estimates. To account for multiple hypothesis testing across organelle features within each cell, p-values were adjusted using the Benjamini–Hochberg false discovery rate procedure.

Adjusted p-values were then used to define spatially significant organelle organization, with significance determined at an FDR-adjusted threshold of 0.05. Binary cell-by-organelle matrices were generated from these adjusted significance calls to summarize whether each organelle class showed significant spatial organization within each cell. These matrices were used to visualize condition-specific patterns of spatial organization across the single-cell population.

Group-level comparisons across dietary conditions were subsequently performed using STORM-derived distributional metrics to quantify shifts in spatial organization across conditions. Condition-level differences were evaluated by testing both the proportion of cells exhibiting significant spatial organization and the magnitude of STORM-derived spatial effect sizes. Proportion differences were assessed using large-sample proportion tests, whereas effect-size differences were evaluated using Welch’s two-sample t-tests. Statistical comparisons of significance frequencies and spatial effect-size distributions quantified both the prevalence of significant spatial organization and differences in the magnitude of spatial dependence between conditions. In parallel, the calculated effect sizes were compared across conditions to evaluate differences in the magnitude of spatial organization independent of statistical significance alone.

## Data availability

All relvant data supporting the key findings of this study are available within the article and its Supplementary files. The stxbrain data can be downloaded with the “SeuratData” package in R. HDST data are available at Broad Institute’s single-cell repository with ID SCP420. The spatial organellomics dataset analyzed in this study is deposited in Figshare at https://figshare.com/s/54fc2f9698757f3ff5aa.^1^ The dataset includes organelle-level morphology measurements, centroid coordinates, cell identifiers, organelle class and subtype annotations, and information for mouse liver samples collected under control-fed, overnight-fasted, and Western diet–fed conditions.

## Code availability

All source code used in our experiments has been deposited at https://github.com/CastleLi/STORM/.

## Acknowledgements

This work is supported by National Institutes of Health grants R01DK138504 (to J.W. and Q.M.), R35GM126985 (to D.X.), R01GM152585, P01CA278732, U54AG075931, and P01AI177687 (to Q.M.), U19AG074879, R01AG019771, P30AG072976, U01AG072177, U01AG068057 (to M.Y.), NS121718 (to J-Q.K), R25HG012325 (to K.M.), R21DK140693 (to A.M.), the AnalytiXIN initiative (to J.W.), and the Alzheimer’s Association grants AARF-22-722571 (to M.Y.), as well as the Pelotonia Institute of Immuno-Oncology (PIIO) (to Q.M.). The Kidney Precision Medicine Project (KPMP) is supported by the National Institute of Diabetes and Digestive and Kidney Diseases (NIDDK) through the following grants: U01DK133081, U01DK133091, U01DK133092, U01DK133093, U01DK133095, U01DK133097, U01DK114866, U01DK114908, U01DK133090, U01DK133113, U01DK133766, U01DK133768, U01DK114907, U01DK114920, U01DK114923, U01DK114933, U24DK114886, UH3DK114926, UH3DK114861, UH3DK114915, and UH3DK114937. Research reported in this publication was partly supported by the Office of The Director, National Institutes of Health under Award Number U54DK134301 (HuBMAP consortium, S.J.). We gratefully acknowledge the essential contributions of our patient participants and the support of the American public through their tax dollars. The content is solely the responsibility of the authors and does not necessarily represent the official views of the National Institutes of Health. The authors acknowledge the University of Michigan Medical School Central Biorepository (RRID:SCR_026845) for providing biospecimen storage, management, and distribution services in support of the research reported in this publication/grant application/presentation.

## Contributions

Conceptualization: J.L., Q.M., J.W., and D.X.; methodology: J.L. and X.Y.; software coding: J.L. and Y.W.; data collection and investigation: J.H., A.C.R., P.H.N., M.V., M.L.C., L.B., M.K., S.J., P.C.D., T.M.E., J.L., D.F., M.R., M.T.E., and J.W.; data analysis: J.L., M.R., Y.W., S.Z., Y.Y., X.J., Y.C., and J.W.; pathology analysis: M.R., Y.C., R.F., and M.T.E.; manuscript writing, review, and editing: J.L., M.R., M.T.E., R.M.F., Q.M., J.W., and D.X.

## Footnotes

1 The link is currently set as a private reviewer-accessible record and is available for editorial and peer-review evaluation. Because these data derive from a companion spatial organellomics study currently under revision, the Figshare record will remain private during review and will be made publicly available with a citable DOI upon publication of the companion study or no later than publication of this article, in accordance with journal policy.

