## Supplementary Figures for "A Principled Statistical Framework for Analyzing Spatial Patterns in Spatially Resolved Multi-Omics"

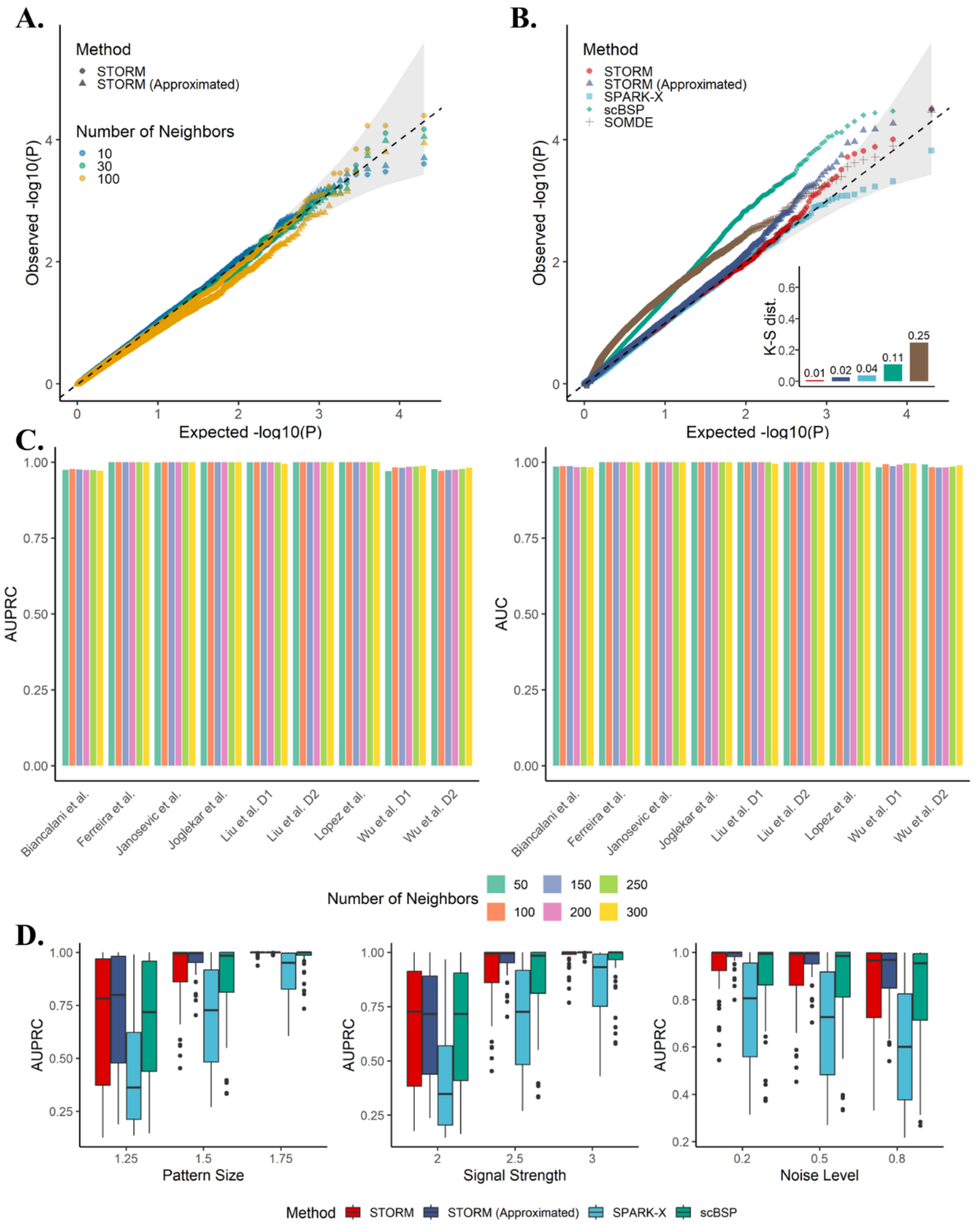


**Supplementary Fig. 1: A.** Type I error of STORM and its approximation with varied numbers of neighbors. **B.** Type I error of STORM and other methods under null hypothesis for high-resolution data. SPARK, SpatialDE, nnSVG, and Moran's I were excluded from this analysis due to the limited computational scalability. **C.** Comparisons of AUPRC and AUC scores with varied numbers of neighbors. **D.** Comparisons of AUPRC scores on 3D simulations.

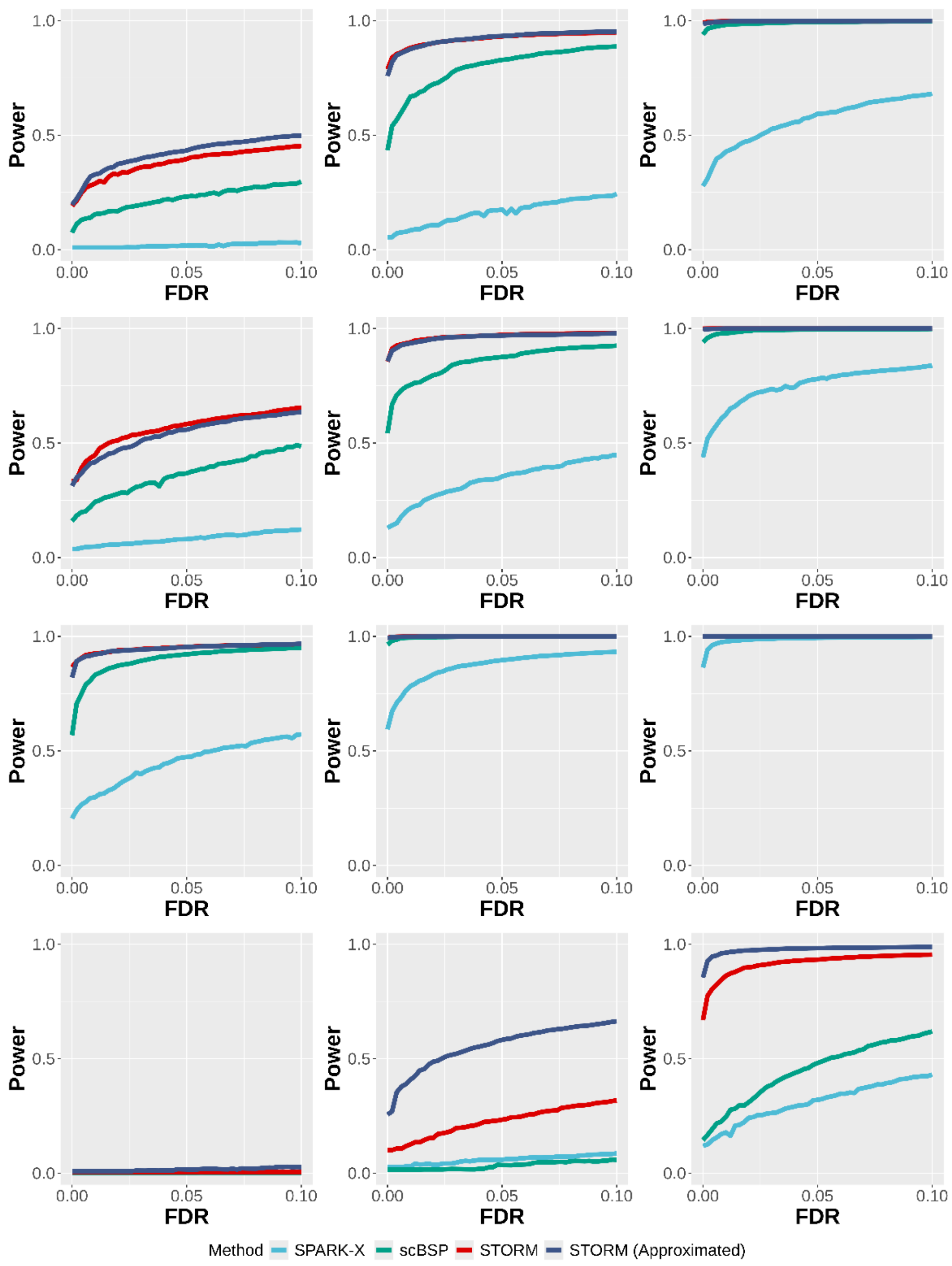


**Supplementary Fig. 2:** Statistical power on 3D simulations with varied pattern sizes. Pattern size was measured as the radius of the pattern as described in the Method section. Power curves were drawn using the averaged statistical power (y-axis) across ten replicates against the false discovery rates (x-axis) for the detected SVFs from each method. Results with small, moderate, and large pattern sizes are shown in the left, middle, and right columns for each of the four spatial patterns as shown in Figure 1E (top to bottom). All simulations were generated using a fixed moderate signal strength and noise level.

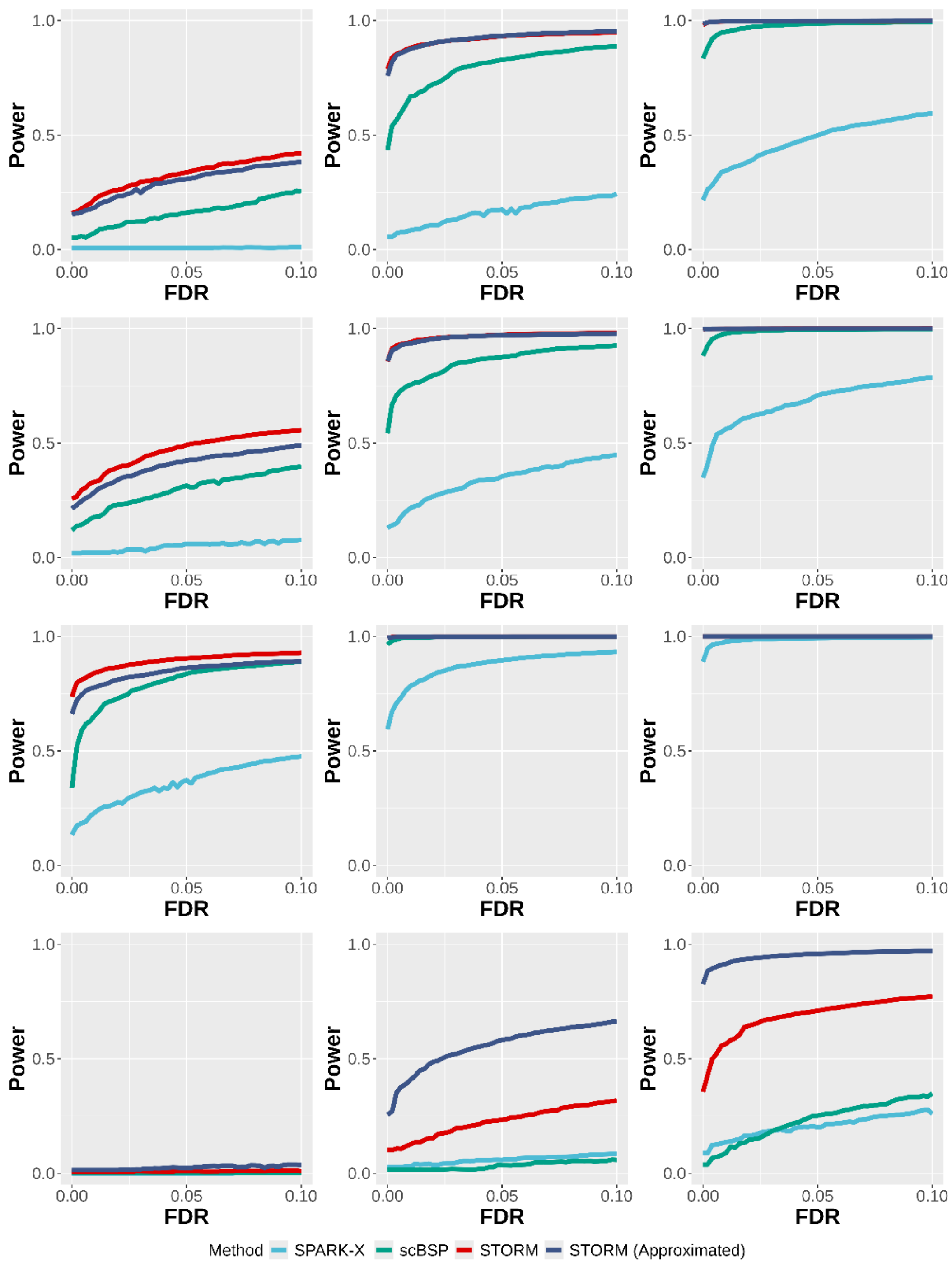


**Supplementary Fig. 3:** Statistical power on 3D simulations with varied signal strengths. Signal strength was measured as the fold changes of expressions between the spiked and remaining regions as described in the Method section. Power curves were drawn using the averaged statistical power (y-axis) across ten replicates against the false discovery rates (x-axis) for the detected SVFs from each method. Results with small, moderate, and large signal strengths are shown in the left, middle, and right columns for each of the four spatial patterns as shown in Figure 1E (top to bottom). All simulations were generated using a fixed moderate spatial pattern and noise level.

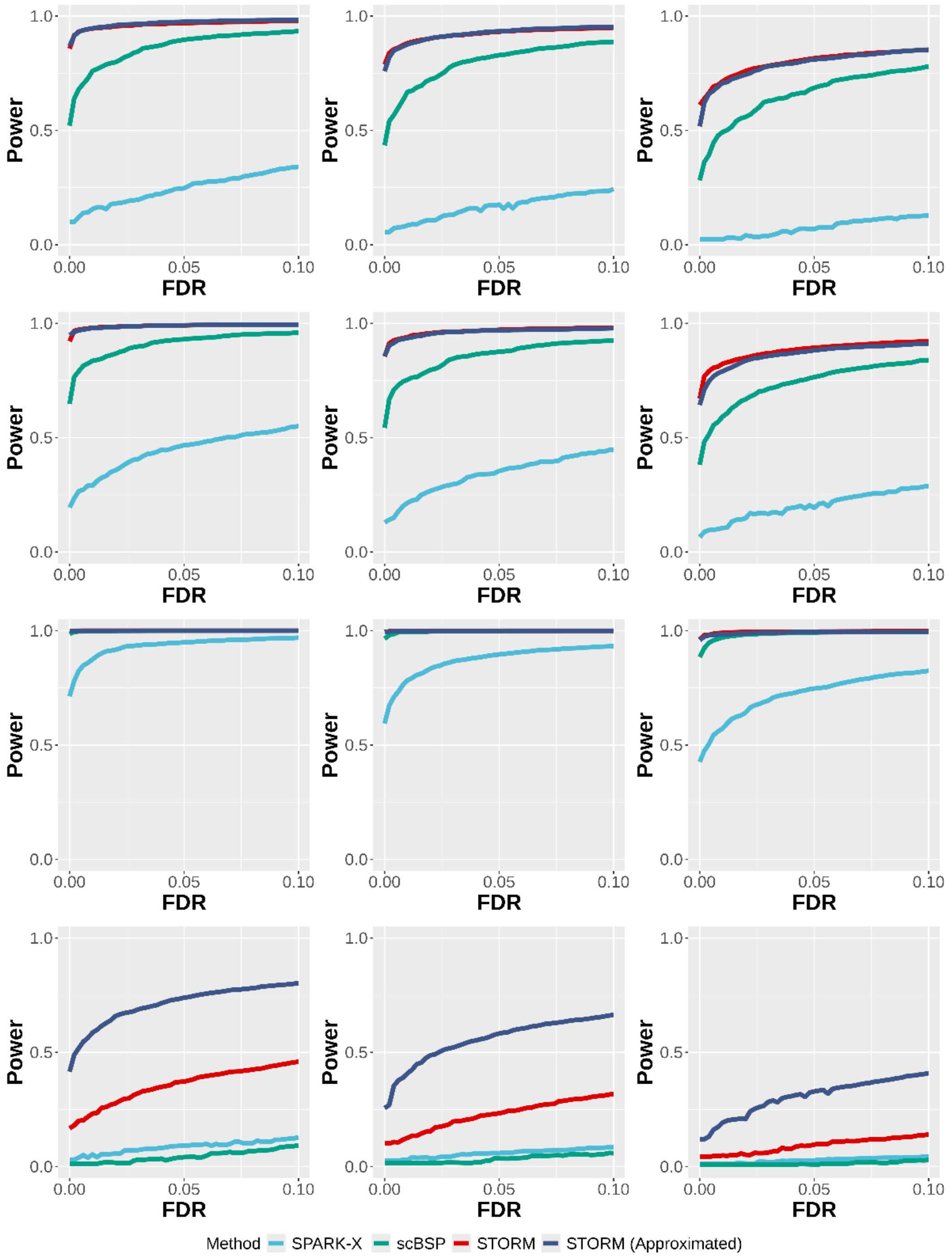


**Supplementary Fig. 4:** Statistical power on 3D simulations with varied noise levels. Noise level was measured as the ratio between the variance of random errors and sample variance as described in the Method section. Power curves were drawn using the averaged statistical power (y-axis) across ten replicates against the false discovery rates (x-axis) for the detected SVFs from each method. Results with small, moderate, and large noise levels are shown in the left, middle, and right columns for each of the four spatial patterns as shown in Figure 1E (top to bottom). All simulations were generated using a fixed moderate spatial pattern and signal strength.

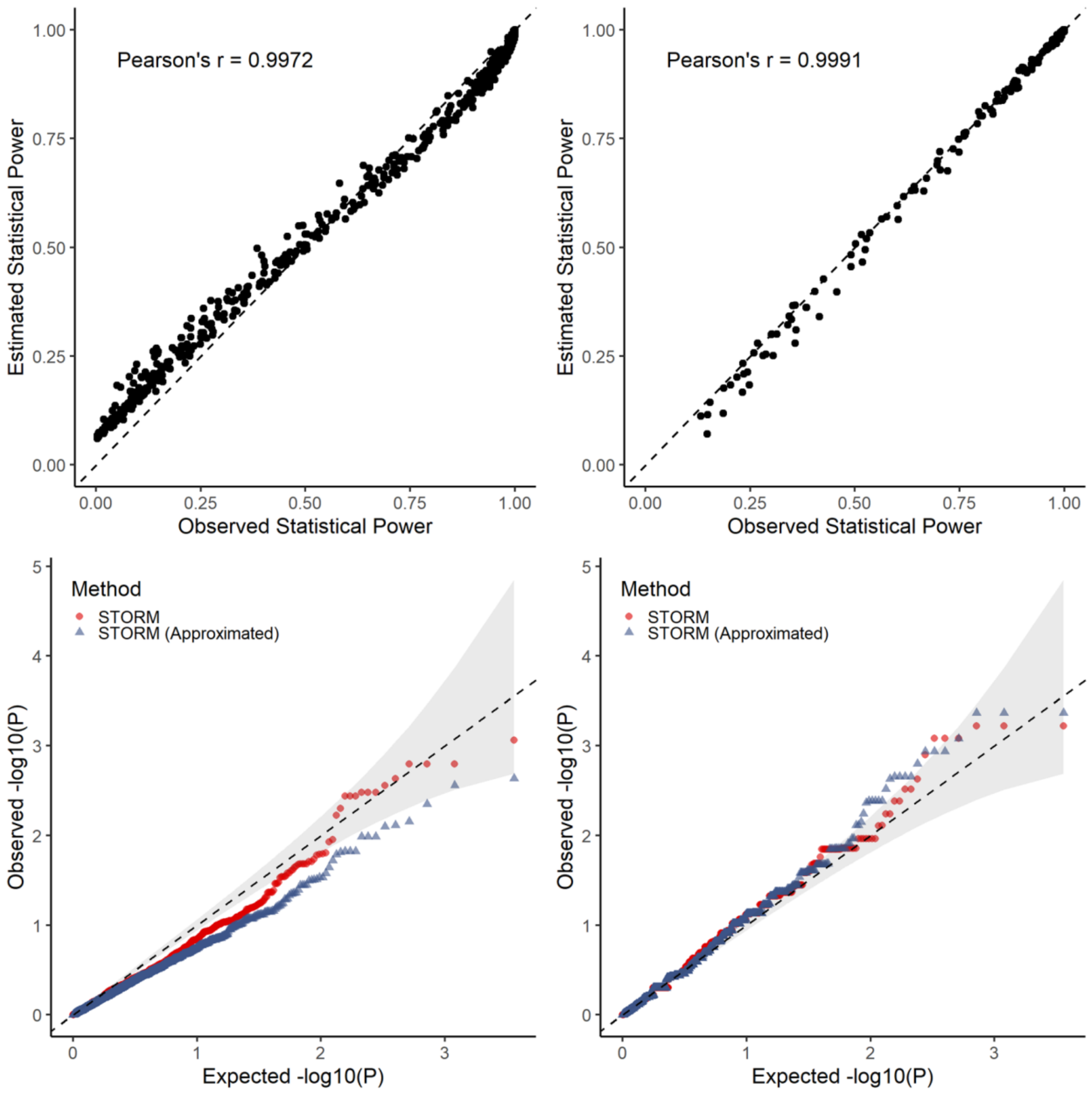


**Supplementary Fig. 5:** Statistical calibration for case–case (left) and case–control (right) comparisons. The top panels show the alignment of the estimated and observed statistical power using the STORM approximation. The bottom panels assess the calibration of the Type I error rate for both STORM and its approximation. The estimated statistical power for the original STORM method is not shown because its calculation requires prior knowledge of the spatial organization, which is generally unavailable in practical applications.

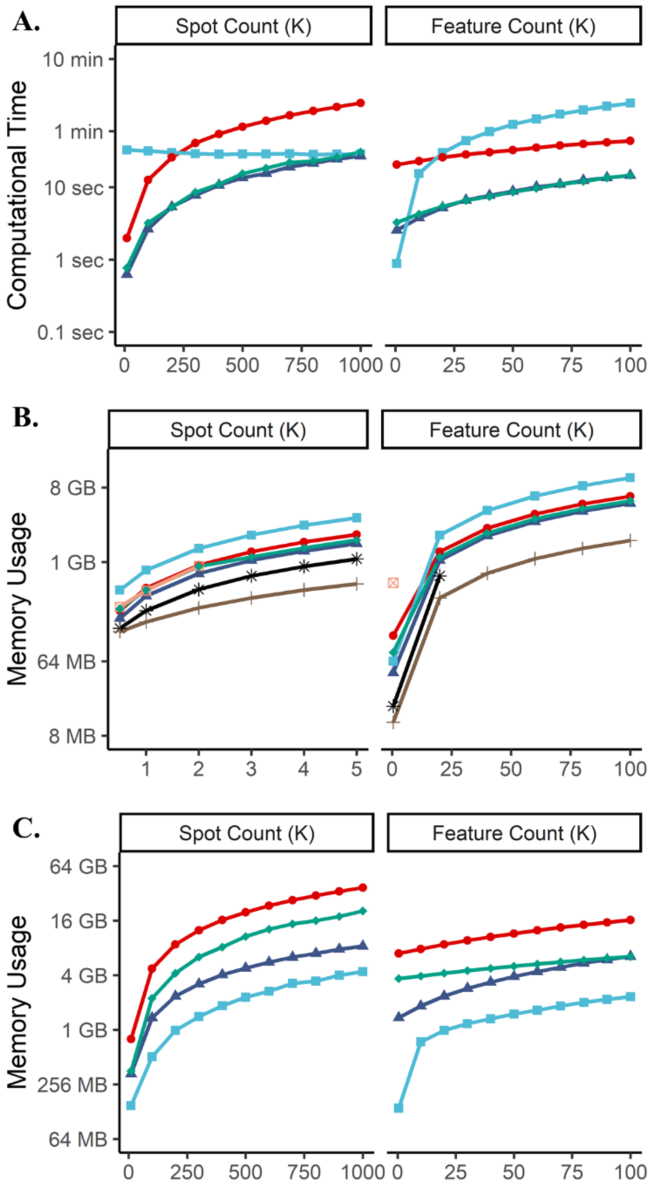


**Supplementary Fig. 6: A.** Computational time (y-axis) for analyzing spatial omics data comprising 20,000 features across 200,000 spots, varying number of spots and features while keeping the other constant. **B.** Memory usage (y-axis) for analyzing spatial omics data comprising 20,000 features across 3,000 spots. **C.** Memory usage (y-axis) for analyzing spatial omics data comprising 20,000 features across 200,000 spots.

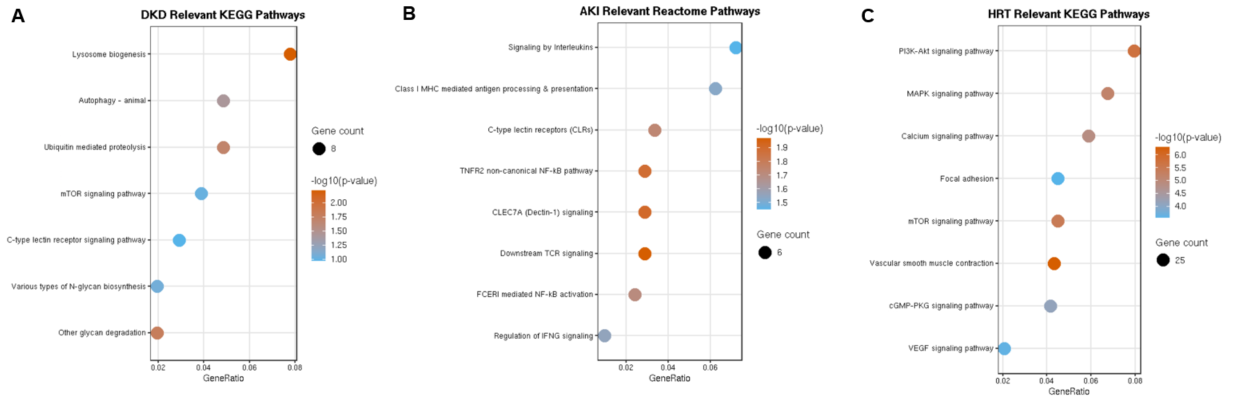


**Supplementary Fig. 7:** Condition-relevant pathway enrichment from STORM-prioritized Visium SVFs. **A.** DKD-relevant KEGG pathways**. B.** AKI-relevant Reactome pathways. **C.** HRT-relevant KEGG pathways. Dot size represents gene count and color indicates −log10(P value).

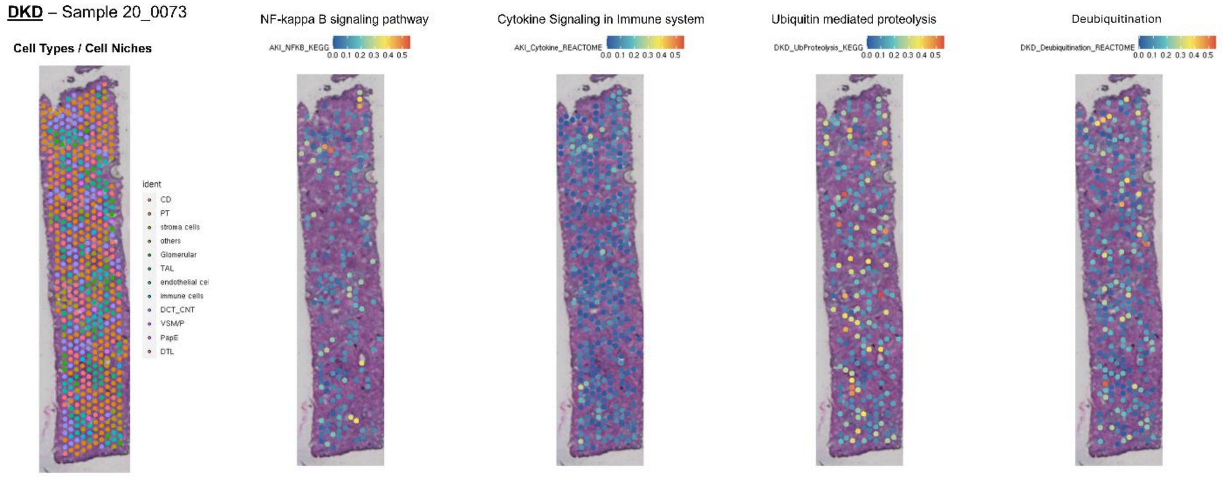


**Supplementary Fig. 8:** Representative DKD 10x Visium biopsy showing cell-type/cell-niche annotations and spatial overlays of pathway expression scores for NF-κB signaling, cytokine signaling, ubiquitin-mediated proteolysis, and deubiquitination. Pathway scores are mapped across tissue spots to visualize localized inflammatory and proteostasis-related activity within the biopsy.

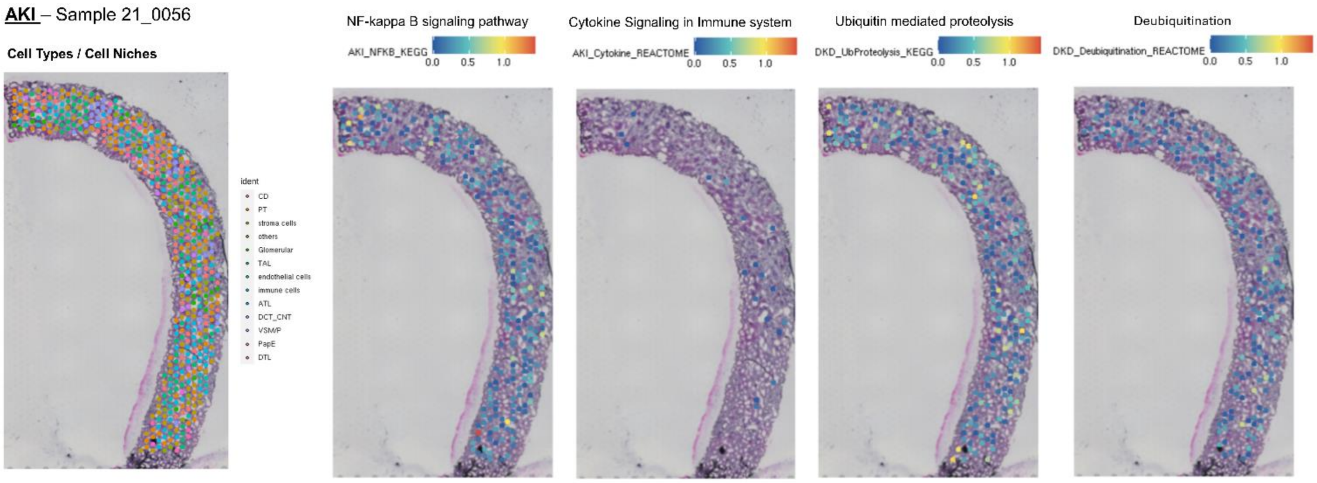


**Supplementary Fig. 9:** Representative AKI 10x Visium biopsy showing cell-type/cell-niche annotations and spatial overlays of pathway expression scores for NF-κB signaling, cytokine signaling, ubiquitin-mediated proteolysis, and deubiquitination. Pathway scores are mapped across tissue spots to visualize localized inflammatory and proteostasis-related activity within the biopsy.

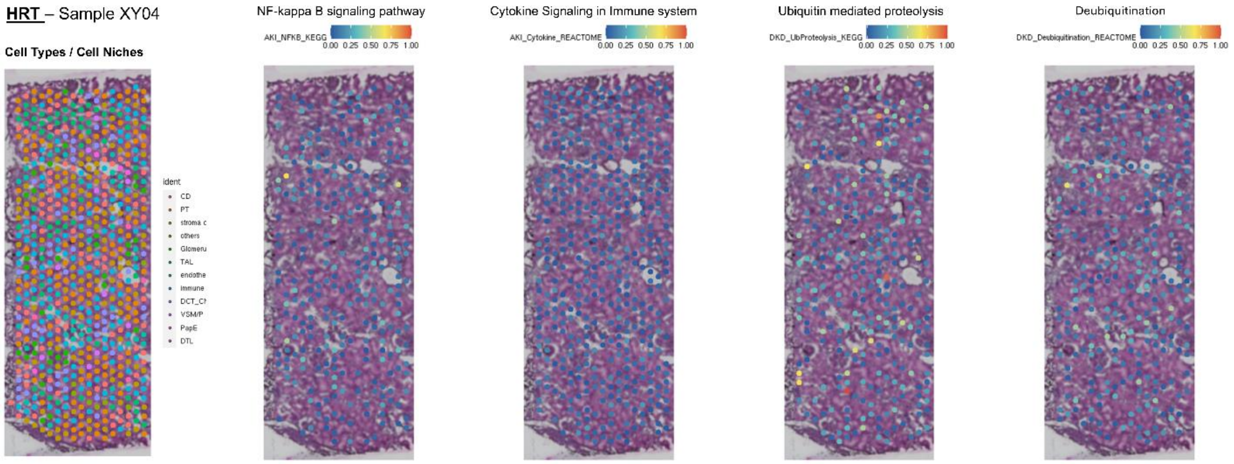


**Supplementary Fig. 10:** Representative HRT 10x Visium biopsy showing cell-type/cell-niche annotations and spatial overlays of pathway expression scores for NF-κB signaling, cytokine signaling, ubiquitin-mediated proteolysis, and deubiquitination. Pathway scores are mapped across tissue spots to visualize localized inflammatory and proteostasis-related activity within the biopsy.

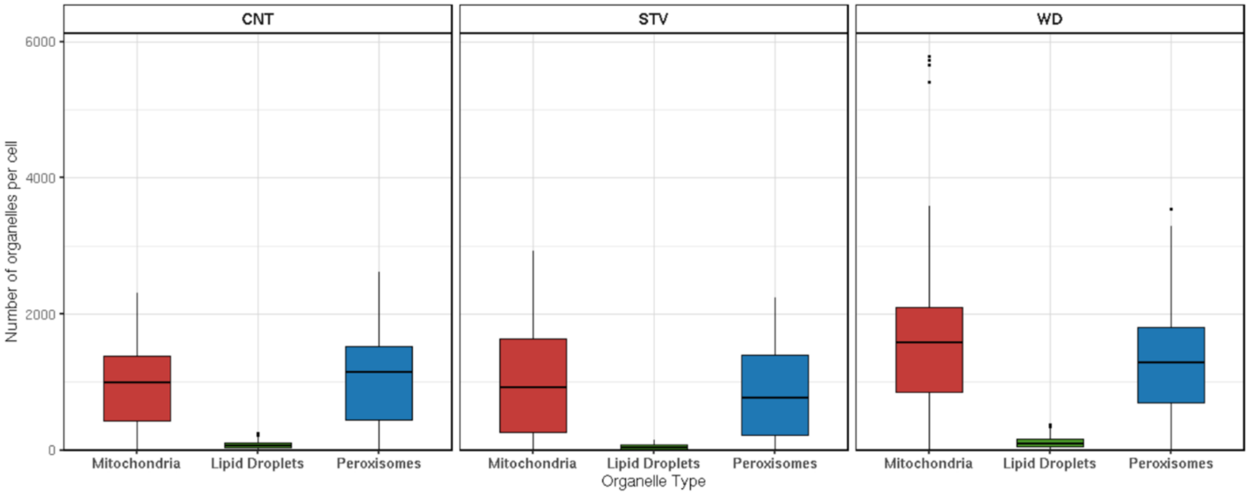


**Supplementary Fig. 11:** Boxplots showing the number of mitochondria, lipid droplets, and peroxisomes per cell across control (CNT), starvation (STV), and Western diet (WD) conditions. WD cells exhibit increased organelle abundance, particularly for mitochondria and peroxisomes, compared to CNT and STV, while lipid droplets remain relatively low across all conditions.
