## Supplementary Notes for "A Principled Statistical Framework for Analyzing Spatial Patterns in Spatially Resolved Multi-Omics"

**Supplementary Note 1**

**Full Derivation of Test Statistics**

For a spatially resolved multi-omics sample with $M$ spots and $N$ features, the $D$-dimensional coordinates of spot $i$ are denoted as a $D$-vector $\boldsymbol{s}_i$, where $i = 1, \ldots, M$, and the corresponding coordinates matrix of all spots are denoted as a $M \times D$ matrix $\boldsymbol{S} = (\boldsymbol{s}_1^T, \ldots, \boldsymbol{s}_M^T)^T$. The expression of a feature $j$ is denoted as $\boldsymbol{x_j} = (x_{1j}, \ldots, x_{Mj})$, where $j = 1, \ldots, N$, and the expression at the spot $i$ is denoted as $x_{ij}$. For each spot in the space, the $K$ nearest neighboring spots are determined, denoted as a binary M-vector $\boldsymbol{b}_{\boldsymbol{i_0}}^{(\boldsymbol{K})} = \left(b_{i_0 1}^{(K)}, \ldots, b_{i_0 M}^{(K)}\right)$, where $b_{i_0 i}^{(K)} = 1$ if spot $i$ is within the $K$ nearest neighboring spots of spot $i_0$. For each feature $j$ at each spot $i$, the sum of the expressions of all neighboring spots, $T_{ij}^{(K)}$, can be calculated as $T_{ij}^{(K)} = \boldsymbol{b}_{\boldsymbol{i}}^{(\boldsymbol{K})} \boldsymbol{x}_{\boldsymbol{j}}^{\boldsymbol{T}}$. The <u>*null hypothesis*</u> is that a feature is not a spatially variable feature, i.e., the expression of a feature does not rely on the spatial location of spots ($\boldsymbol{x_j} \perp \boldsymbol{S}$).

Denote the distribution of feature $j$ as $F_j(x)$ with mean and variance as $\mu_{jx}$ and $\sigma_{jx}^2$. Under null hypothesis, the expression at each spot is independent of each other, i.e., $x_{1j}, \ldots, x_{Mj}\ i.i.d. \sim F_j(x)$. In this case, the mean and variance of $T_{ij}^{(K)}$ can be computed as $K\mu_{jx}$ and $K\sigma_{jx}^2$. Moreover, since $T_{ij}^{(K)}$ and $Kx_{ij}$ are independent for a fixed constant of $K$, the expectation and variance of their difference, $\delta_{ij} = T_{ij}^{(K)} - Kx_{ij}$, can be derived as

$$E[\delta_{ij}] = E\left[T_{ij}^{(K)}\right] - E[Kx_{ij}] = K\mu_{jx} - K\mu_{jx} = 0$$

and

$$Var(\delta_{ij}) = Var\left(T_{ij}^{(K)}\right) + Var(Kx_{ij}) - 2Cov(T_{ij}^{(K)}, Kx_{ij}) = K\sigma_{jx}^2 + K^2\sigma_{jx}^2 = K(K+1)\sigma_{jx}^2$$

where $Cov\left(T_{ij}^{(K)}, Kx_{ij}\right) = 0$ under null hypothesis.

Considering a stationary $m$-dependent sequence of $\boldsymbol{\delta_j^2} = \left(\delta_{1j}^2, \dots \delta_{Mj}^2\right)^T$ where $m < M$, the expectation and variance of each term are

$$E\left[\delta_{ij}^2\right] = Var\left(\delta_{ij}\right) + E\left[\delta_{ij}\right]^2 = K(K+1)\sigma_{jx}^2$$

and

$$Var\left(\delta_{ij}^2\right) = E\left[\delta_{ij}^4\right] - \left(E\left[\delta_{ij}^2\right]\right)^2 = (\kappa_j - 1)\left(E\left[\delta_{ij}^2\right]\right)^2 = (\kappa_j - 1)K^2(K+1)^2\sigma_{jx}^4$$

where $\kappa_j$ is the kurtosis of $\boldsymbol{\delta_{ij}}$. Let

$$\sigma_j^2 = K(K+1)\sigma_{jx}^2 \quad (1)$$

we have

$$\sqrt{M}\left(\overline{\delta_{ij}^2} - \sigma_j^2\right) \sim N\left(0, (\kappa_j - 1)\sigma_j^4 + \frac{1}{M}\sum_{p \neq q} Cov\left(\delta_{pj}^2, \delta_{qj}^2\right)\right) \quad (2)$$

based on the Central Limit Theorem (CLT) for $m$-dependent random variables. The sum of covariance between $\delta_{pj}^2$ and $\delta_{qj}^2$ can be derived as (**Supplementary Note 2**).

$$2MK^3\left(\kappa_{jx} - 1\right)\sigma_{jx}^4 + \left(\kappa_{jx} - 3\right)\sigma_{jx}^4 w_{n1} + 2\sigma_{jx}^4 w_{n2} - 8Kw_{n3}\sigma_{jx}^4 + 4K^2 w_{n4}\sigma_{jx}^4$$

where $w_{n1}$, $w_{n2}$, $w_{n3}$ and $w_{n4}$ represent the total number of shared neighbors between all distinct spot pairs, sum of squared number of shared neighbors between all distinct spot pairs, total number of shared neighbors for directly connected pairs, and total number of mutual nearest neighbors (pairs of spots that are each other's nearest neighbors), which can be computed as follows

$$w_{n1} = \sum_{p \neq q} n = \mathbf{1}^{\boldsymbol{T}} \boldsymbol{W_{n1}} \mathbf{1}$$

$$w_{n2} = \sum_{p \neq q} n^2 = \mathbf{1}^{\boldsymbol{T}} \boldsymbol{W_{n2}} \mathbf{1}$$

$$w_{n3} = \sum_{q \in KNN(p)} n = \mathbf{1}^{\boldsymbol{T}} \boldsymbol{W}_{\boldsymbol{n3}} \mathbf{1}$$

$$w_{n4} = \sum_{p \in KNN(q), q \in KNN(p)} 1 = \mathbf{1}^{\boldsymbol{T}} \boldsymbol{W}_{\boldsymbol{n4}} \mathbf{1}$$

where $\boldsymbol{W}_{\boldsymbol{n1}} = \boldsymbol{B}\boldsymbol{B}^{\boldsymbol{T}} - K\boldsymbol{I}_{\boldsymbol{M}}$, $\boldsymbol{W}_{\boldsymbol{n2}} = (\boldsymbol{B}\boldsymbol{B}^{\boldsymbol{T}})^2 - K^2\boldsymbol{I}_{\boldsymbol{M}}$, $\boldsymbol{W}_{\boldsymbol{n3}} = \boldsymbol{B} \circ (\boldsymbol{B}\boldsymbol{B}^{\boldsymbol{T}})$, $\boldsymbol{W}_{\boldsymbol{n4}} = \boldsymbol{B} \circ \boldsymbol{B}^{\boldsymbol{T}}$, $\boldsymbol{B} = \left(\boldsymbol{b}_{\mathbf{1}}^{(K)^{\boldsymbol{T}}}, \dots, \boldsymbol{b}_{\boldsymbol{M}}^{(K)^{\boldsymbol{T}}}\right)^T$ and $\boldsymbol{I}_{\boldsymbol{M}}$ is an $M \times M$ identity matrix. The value of kurtosis $\kappa_j$ could be estimated from the kurtosis of $x_{ij}$ (**Supplementary Note 3**) as

$$\kappa_j = \frac{(K^3 + 1)\kappa_{jx} + 6K^2 + 3(K - 1)}{K(K + 1)^2} \tag{3}$$

where $\kappa_{jx}$ is the kurtosis of $\boldsymbol{x}_{\boldsymbol{j}}$ which can be determined from the sample. Taking Equation (3) into Equation (2) and using the approximation that $w_{n1} \approx MK^2$, the variance of $\sqrt{M}\left(\overline{\delta_{ij}^2} - \sigma_j^2\right)$ can be computed as

$$Var\left(\sqrt{M}\left(\overline{\delta_{ij}^2} - \sigma_j^2\right)\right) = \left(\left(\kappa_j - 1\right)K^2(K + 1)^2 + 4K^3 + \frac{2w_{n2} - 8Kw_{n3} + 4K^2 w_{n4}}{M}\right)\sigma_{jx}^4$$

Implementing Equation (1), the variance can be rewritten as follows

$$Var\left(\sqrt{M}\left(\overline{\delta_{ij}^2} - \sigma_j^2\right)\right) = \left(\left(\kappa_j - 1\right) + \frac{4K^3}{K^2(K + 1)^2} + \frac{2w_{n2} - 8Kw_{n3} + 4K^2 w_{n4}}{MK^2(K + 1)^2}\right)\sigma_j^4$$

Since $\sigma_j^2$ can be estimated from the sample as $\hat{\sigma}_j{}^2 = K(K + 1)\hat{\sigma}_{jx}{}^2$ which is greater than 0 and finite, the variance of $(\overline{\delta_{ij}^2} - \hat{\sigma}_j{}^2)/\hat{\sigma}_j{}^2$ can be derived from the first-order of Taylor expansion as

$$Var\left(\frac{\overline{\delta_{ij}^2} - \hat{\sigma}_j{}^2}{\hat{\sigma}_j{}^2}\right) \approx \frac{Var\left(\overline{\delta_{ij}^2}\right)}{\left(E\left[\hat{\sigma}_j{}^2\right]\right)^2} - \frac{2E\left[\overline{\delta_{ij}^2}\right]Cov\left(\overline{\delta_{ij}^2}, \hat{\sigma}_j{}^2\right)}{\left(E\left[\hat{\sigma}_j{}^2\right]\right)^3} + \frac{\left(E\left[\overline{\delta_{ij}^2}\right]\right)^2 Var\left(\hat{\sigma}_j{}^2\right)}{\left(E\left[\hat{\sigma}_j{}^2\right]\right)^4}$$

which is calculated $\frac{4K^3}{MK^2(K+1)^2} + \frac{2w_{n2} - 8Kw_{n3} + 4K^2 w_{n4}}{M^2K^2(K+1)^2}$ (**Supplementary Note 4**). Therefore, the test statistics can be converted to the standard normal form

$$Z = \sqrt{\frac{MK^2(K+1)^2}{4K^3 + \frac{2w_{n2} - 8Kw_{n3} + 4K^2 w_{n4}}{M}}} \left( \frac{\overline{\delta_{ij}^2}}{\hat{\sigma}_j{}^2} - 1 \right) \sim N(0,1) \tag{4}$$

**Supplementary Note 2**

For convenience, let $x_i = x_{ij} - \mu_{jx}$, $T_i = T_{ij}^{(K)} - K\mu_{jx}$, and $\delta_i = T_i - Kx_i = T_{ij}^{(K)} - x_{ij} = \delta_{ij}$. The expectation and variance of $x_i$ can be derived as $E[x_i] = 0$ and $Var(x_i) = E[x_i^2] = \sigma_{jx}^2$. Denote the kurtosis of $x_i$ as $\kappa$, $\kappa = \kappa_{jx}$ as the kurtosis is invariant with the centering. Similarly, the expectation and variance for $T_i$ can be computed as 0 and $K\sigma_{jx}^2$. Based on the definition of covariance, the covariance between $\delta_p^2$ and $\delta_q^2$ can be derived as

$$Cov\left(\delta_p^2, \delta_q^2\right) = E\left[\delta_p^2\delta_q^2\right] - E\left[\delta_p^2\right]E\left[\delta_q^2\right]$$

Expanding the term of $\delta_p^2\delta_q^2$ using $\delta_i = T_i - Kx_i$ and taking expectations:

$$E\left[\delta_p^2\delta_q^2\right] = E\left[\left(T_p - Kx_p\right)^2\left(T_q - Kx_q\right)^2\right]$$

$$= E\left[T_p^2T_q^2\right] - 2KE\left[T_p^2T_qx_q\right] + K^2E\left[T_p^2x_q^2\right] - 2KE\left[T_px_pT_q^2\right] + 4K^2E\left[T_px_pT_qx_q\right]$$

$$- 2K^3E\left[T_px_px_q^2\right] + K^2E\left[x_p^2T_q^2\right] - 2K^3E\left[x_p^2T_qx_q\right] + K^4E\left[x_p^2x_q^2\right]$$

For the first term, let

$$T_{pq} = \sum_{i \in KNN(p) \cap KNN(q)} x_i$$

which includes $n$ overlapping terms between $T_p$ and $T_q$. Denote $\widetilde{T_p} = T_p - T_{pq}$ and $\widetilde{T_q} = T_q - T_{pq}$, $T_{pq}$, $\widetilde{T_p}$ and $\widetilde{T_q}$ are independent with mean 0. Expanding $E\left[T_p^2T_q^2\right]$ and vanishing the odd-order terms:

$$E\left[T_p^2T_q^2\right] = E\left[\widetilde{T_p}^2\widetilde{T_q}^2\right] + E\left[\widetilde{T_p}^2T_{pq}^2\right] + E\left[\widetilde{T_q}^2T_{pq}^2\right] + E\left[T_{pq}^4\right]$$

$$= E\left[\widetilde{T_p}^2\right]E\left[\widetilde{T_q}^2\right] + E\left[\widetilde{T_p}^2\right]E\left[T_{pq}^2\right] + E\left[\widetilde{T_q}^2\right]E\left[T_{pq}^2\right] + E\left[T_{pq}^4\right]$$

$$= (K-n)\sigma_{jx}^2(K-n)\sigma_{jx}^2 + 2(K-n)\sigma_{jx}^2n\sigma_{jx}^2 + \left(3 + \frac{\kappa - 3}{n}\right)\left(n\sigma_{jx}^2\right)^2$$

since the kurtosis of $T_{pq}$ is $\left(3 + \frac{\kappa - 3}{n}\right)$, where $n$ is the number of overlapping terms and $\kappa$ is the kurtosis of $x_i$. This equation can be simplified to

$$E\left[T_p^2 T_q^2\right] = K^2 \sigma_{jx}^4 + 2n^2 \sigma_{jx}^4 + (\kappa - 3) n \sigma_{jx}^4$$

Taking sum of $E\left[T_p^2 T_q^2\right]$ for $p \neq q$,

$$\sum_{p \neq q} E\left[T_p^2 T_q^2\right] = M(M-1) K^2 \sigma_{jx}^4 + 2\sigma_{jx}^4 \sum_{p \neq q} n^2 + (\kappa - 3) \sigma_{jx}^4 \sum_{p \neq q} n$$

Let $\boldsymbol{W_{n1}} = \boldsymbol{B}\boldsymbol{B^T} - K\boldsymbol{I_M}$ and $\boldsymbol{W_{n2}} = (\boldsymbol{B}\boldsymbol{B^T})^2 - K^2 \boldsymbol{I_M}$, where $\boldsymbol{B} = \left(\boldsymbol{b_1^{(K)}}^{\boldsymbol{T}}, \dots, \boldsymbol{b_M^{(K)}}^{\boldsymbol{T}}\right)^T$ and $\boldsymbol{I_M}$ is an $M \times M$ identity matrix, the sum of overlapping $w_{n1}$ and $w_{n2}$ can be calculated as follows

$$w_{n1} = \sum_{p \neq q} n = \boldsymbol{1^T} \boldsymbol{W_{n1}} \boldsymbol{1}$$

and

$$w_{n2} = \sum_{p \neq q} n^2 = \boldsymbol{1^T} \boldsymbol{W_{n2}} \boldsymbol{1}$$

where $\boldsymbol{1}$ is a column vector of ones.

For the second term, the expectation is 0 when $q \notin KNN(p)$ since $x_q$ is independent of $T_q$ and $T_p$ and $E\left[x_q\right] = 0$, i.e., $E\left[T_p^2 T_q x_q\right] = E\left[x_q\right] E\left[T_p^2 T_q\right] = 0$. If $q \in KNN(p)$, let $\widetilde{T_{pq}} = \widetilde{T_p} - x_q$ and the term $T_p^2 T_q x_q$ can be rewritten as

$$T_p^2 T_q x_q = \left(T_{pq} + \widetilde{T_{pq}} + x_q\right)^2 \left(T_{pq} + \widetilde{T_q}\right) x_q$$

Taking expectations and vanishing the odd-order terms:

$$E\left[T_p^2 T_q x_q\right] = 2E\left[T_{pq}^2 x_q^2\right] = 2E\left[T_{pq}^2\right] E\left[x_q^2\right] = 2n\sigma_{jx}^4$$

Taking sum of $p \neq q$:

$$\sum_{p \neq q} E\left[T_p^2 T_q x_q\right] = 2\sigma_{jx}^4 \sum_{q \in KNN(p)} n$$

Let $\boldsymbol{W_{n3}} = \boldsymbol{B} \circ (\boldsymbol{B}\boldsymbol{B}^{\boldsymbol{T}})$, the sum of overlapping $w_{n3}$ can be calculated as follows

$$w_{n3} = \sum_{q \in KNN(p)} n = \mathbf{1}^{\boldsymbol{T}} \boldsymbol{W_{n3}} \mathbf{1}$$

The second and fourth terms are thereby $-4Kw_{n3}\sigma_{jx}^4$.

For the third and seventh terms, The expectation can be computed as

$$E\left[T_p^2 x_q^2\right] = \begin{cases} E[\widetilde{T_{pq}}^2 x_q^2 + 2\widetilde{T_{pq}} x_q^3 + x_q^4] = (K-1)\sigma_{jx}^4 + \kappa\sigma_{jx}^4, & \text{if } q \in KNN(p), \\ E\left[T_p^2\right]E\left[x_q^2\right] = K\sigma_{jx}^4, & otherwise \end{cases}$$

where $\widetilde{T_{pq}} = T_p - x_q$. Summing this term for $p \neq q$, we have

$$\sum_{p \neq q} E\left[T_p^2 x_q^2\right] = MK\left((K-1)\sigma_{jx}^4 + \kappa\sigma_{jx}^4\right) + M(M-K-1)K\sigma_{jx}^4$$

which can be simplified as

$$K^2 \sum_{p \neq q} E\left[T_p^2 x_q^2\right] = M(M-1)K^3\sigma_{jx}^4 + MK^3(\kappa - 1)\sigma_{jx}^4$$

For the fifth term, the expectation $E\left[T_p x_p T_q x_q\right]$ is not zero if and only if $p \in KNN(q)$ and $q \in KNN(p)$, where the expectation would be

$$E\left[T_p x_p T_q x_q\right] = E\left[(T_{pq} + \widetilde{T_{pq}} + x_q)x_p(T_{pq} + \widetilde{T_{qp}} + x_p)x_q\right]$$

Expanding the right side and vanishing the odd-order terms:

$$E\left[T_p x_p T_q x_q\right] = E\left[x_p^2 x_q^2\right] = \sigma_{jx}^4$$

Summing this term for $p \neq q$:

$$\sum_{p \neq q} E\left[T_p x_p T_q x_q\right] = \sigma_{jx}^4 \sum_{p \in KNN(q), q \in KNN(p)} 1$$

Let $\boldsymbol{W_{n4}} = \boldsymbol{B} \circ \boldsymbol{B}^{\boldsymbol{T}}$ the sum of overlapping $w_{n4}$ can be calculated as follows

$$w_{n4} = \sum_{p \in KNN(q), q \in KNN(p)} 1 = \mathbf{1}^{\boldsymbol{T}} \boldsymbol{W_{n4}} \mathbf{1}$$

The fifth term is thereby $4K^2 w_{n4}\sigma_{jx}^4$. Since $x_p$ and $x_q$, $x_p$ and $T_p$ are independent and $E[x_p] = 0$, the sixth term

$$E[T_p x_p x_q^2] = E[x_p]E[T_p x_q^2] = 0$$

Similarly, the eighth term vanishes as well. The sum of last term is

$$K^4 \sum_{p \neq q} E[x_p^2 x_q^2] = K^4 \sum_{p \neq q} E[x_p^2]E[x_q^2] = M(M-1)K^4\sigma_{jx}^4$$

The total sum of the expectations is thereby:

$$\sum_{p \neq q} E[\delta_p^2 \delta_q^2] = M(M-1)K^2\sigma_{jx}^4 + 2\sigma_{jx}^4 w_{n2} + (\kappa - 3)\sigma_{jx}^4 w_{n1} - 8K w_{n3}\sigma_{jx}^4 + 2M(M-1)K^3\sigma_{jx}^4$$

$$+ 2MK^3(\kappa - 1)\sigma_{jx}^4 + 4K^2 w_{n4}\sigma_{jx}^4 + M(M-1)K^4\sigma_{jx}^4$$

The sum of covariances can be derived as

$$\sum_{p \neq q} Cov(\delta_p^2, \delta_q^2) = \sum_{p \neq q} E[\delta_p^2 \delta_q^2] - \sum_{p \neq q} \left(K(K+1)\sigma_{jx}^2\right)^2$$

$$= 2MK^3(\kappa - 1)\sigma_{jx}^4 + (\kappa - 3)\sigma_{jx}^4 w_{n1} + 2\sigma_{jx}^4 w_{n2} - 8K w_{n3}\sigma_{jx}^4 + 4K^2 w_{n4}\sigma_{jx}^4$$

**Supplementary Note 3**

For convenience, let $x_i = x_{ij} - \mu_{jx}$, $T_i = T_{ij}^{(K)} - K\mu_{jx}$, and $\delta_i = T_i - Kx_i = T_{ij}^{(K)} - x_{ij} = \delta_{ij}$. The expectation and variance of $x_i$ can be derived as $E[x_i] = 0$ and $Var(x_i) = E[x_i^2] = \sigma_{jx}^2$. Denote the kurtosis of $x_i$ as $\kappa$, $\kappa = \kappa_{jx}$ as the kurtosis is invariant with the centering. Similarly, the expectation and variance for $T_i$ can be computed as 0 and $K\sigma_{jx}^2$. The fourth moment of a $\delta_{ij}$ is

$$E\left[\delta_{ij}^4\right] = E[(T_i - Kx_i)^4]$$

Since $T_i$ and $x_i$ are independent, the expression above could be expanded as

$$E\left[\delta_{ij}^4\right] = E\left[{T_i}^4\right] + K^4 E[{x_i}^4] - 4KE\left[{T_i}^3\right]E[x_i] + 6K^2 E\left[{T_i}^2\right]E[{x_i}^2] - 4K^3 E[T_i]E[{x_i}^3]$$

The odd-order terms vanish since $E[T_i] = E[x_i] = 0$, and the equation simplifies to:

$$E\left[\delta_{ij}^4\right] = E\left[{T_i}^4\right] + K^4 E[{x_i}^4] + 6K^2 E\left[{T_i}^2\right]E[{x_i}^2]$$

Since

$$E\left[{T_i}^4\right] = KE[{x_i}^4] + 3K(K-1)\sigma_{jx}^4 = K\left(\kappa_{jx}\sigma_{jx}^4\right) + 3K(K-1)\sigma_{jx}^4$$

and

$$E[(Kx_i)^4] = K^4\left(\kappa_{jx}\sigma_{jx}^4\right)$$

we have

$$E\left[\delta_{ij}^4\right] = (K + K^4)\left(\kappa_{jx}\sigma_{jx}^4\right) + 3K(K-1)\sigma_{jx}^4 + 6K^3\sigma_{jx}^4$$

Therefore, the kurtosis for $\delta_{ij}$ can be calculated as

$$\kappa_j = \frac{E\left[\delta_{ij}^4\right]}{E\left[\delta_{ij}^2\right]} = \frac{(K + K^4)\left(\kappa_{jx}\sigma_{jx}^4\right) + 3K(K-1)\sigma_{jx}^4 + 6K^3\sigma_{jx}^4}{\left(K(K+1)\sigma_{jx}^2\right)^2}$$

which can be simplified as

$$\kappa_j = \frac{(K^3 + 1)\kappa_{jx} + 6K^2 + 3(K-1)}{K(K+1)^2}$$

**Supplementary Note 4**

For convenience, let $x_i = x_{ij} - \mu_{jx}$, $T_i = T_{ij}^{(K)} - K\mu_{jx}$, and $\delta_i = T_i - Kx_i = T_{ij}^{(K)} - x_{ij} = \delta_{ij}$. The expectation and variance of $x_i$ can be derived as $E[x_i] = 0$ and $Var(x_i) = E[x_i^2] = \sigma_{jx}^2$. Denote the kurtosis of $x_i$ as $\kappa$, $\kappa = \kappa_{jx}$ as the kurtosis is invariant with the centering. Similarly, the expectation and variance for $T_i$ can be computed as 0 and $K\sigma_{jx}^2$. Based on the definition of covariance, the covariance between $\overline{\delta_{ij}^2}$ and ${\hat{\sigma}_{jx}}^2$ can be derived as

$$Cov\left(\overline{\delta_{ij}^2}, {\hat{\sigma}_{jx}}^2\right) = E\left[\overline{\delta_{ij}^2}{\hat{\sigma}_{jx}}^2\right] - E\left[\overline{\delta_{ij}^2}\right]E\left[{\hat{\sigma}_{jx}}^2\right]$$

Expanding $\delta_i^2$ as $(T_i - Kx_i)^2$, $\overline{\delta_{ij}^2}{\hat{\sigma}_{jx}}^2$can be rewritten as

$$\overline{\delta_{ij}^2}{\hat{\sigma}_{jx}}^2 = \left(\frac{1}{M}\sum_p \delta_p^2\right)\left(\frac{1}{M}\sum_q x_q^2\right) = \frac{1}{M^2}\sum_p\sum_q\left(\left(T_p - Kx_p\right)^2 x_q^2\right)$$

$$= \frac{1}{M^2}\sum_p\sum_q\left(T_p^2x_q^2 - 2KT_px_px_q^2 + K^2x_p^2x_q^2\right)$$

Taking expectations:

$$E\left[\overline{\delta_{ij}^2}{\hat{\sigma}_{jx}}^2\right] = \frac{1}{M^2}\sum_p\sum_q\left(E\left[T_p^2x_q^2\right] - 2KE\left[T_px_px_q^2\right] + K^2E\left[x_p^2x_q^2\right]\right)$$

For the first term $E\left[T_p^2x_q^2\right]$,

$$E\left[T_p^2x_q^2\right] = \begin{cases} E[T_{pq}^2x_q^2 + 2T_{pq}x_q^3 + x_q^4] = (K-1)\sigma_{jx}^4 + \kappa\sigma_{jx}^4, & \text{if } q \in KNN(p), \\ E\left[T_p^2\right]E\left[x_q^2\right] = K\sigma_{jx}^4, & otherwise \end{cases}$$

where $T_{pq} = T_p - x_q$. Summing this term, we have

$$\sum_p\sum_q E\left[T_p^2x_q^2\right] = MK\left((K-1)\sigma_{jx}^4 + \kappa\sigma_{jx}^4\right) + M(M-K)K\sigma_{jx}^4$$

which can be simplified as

$$\sum_p \sum_q E\left[T_p^2 x_q^2\right] = M^2 K \sigma_{jx}^4 + MK(\kappa - 1)\sigma_{jx}^4$$

For the second term $E\left[T_p x_p x_q^2\right]$, since $x_p$ and $T_p$ are independent,

$$E\left[T_p x_p x_q^2\right] = \begin{cases} E\left[x_p\right]E\left[T_p x_q^2\right] = 0, \text{if } q \in KNN(p) \\ E\left[T_p\right]E\left[x_q^3\right] = 0, \quad\quad otherwise \end{cases}$$

For the third term $E\left[x_p^2 x_q^2\right]$,

$$E\left[x_p^2 x_q^2\right] = \begin{cases} E\left[x_p^2\right]E\left[x_q^2\right] = \sigma_{jx}^4, if\ p \neq q \\ E\left[x_q^4\right] = \kappa\sigma_{jx}^4, \quad\quad otherwise \end{cases}$$

Therefore,

$$\sum_p \sum_q E\left[x_p^2 x_q^2\right] = M(M-1)\sigma_{jx}^4 + M\kappa\sigma_{jx}^4 = M^2\sigma_{jx}^4 + M(\kappa - 1)\sigma_{jx}^4$$

The total sum of the expectation is computed as

$$E\left[\overline{\delta_{ij}^2}\hat{\sigma}_{jx}{}^2\right] = \frac{1}{M^2}\left(M^2 K\sigma_{jx}^4 + MK(\kappa - 1)\sigma_{jx}^4 + K^2\left(M^2\sigma_{jx}^4 + M(\kappa - 1)\sigma_{jx}^4\right)\right)$$

which can be simplified to

$$\left[\overline{\delta_{ij}^2}\hat{\sigma}_{jx}{}^2\right] = K(K+1)\sigma_{jx}^4 + \frac{K(K+1)}{M}(\kappa - 1)\sigma_{jx}^4$$

The covariance is thereby:

$$Cov\left(\overline{\delta_{ij}^2}, \hat{\sigma}_{jx}{}^2\right) = E\left[\overline{\delta_{ij}^2}\hat{\sigma}_{jx}{}^2\right] - E\left[\overline{\delta_{ij}^2}\right]E\left[\hat{\sigma}_{jx}{}^2\right] = \frac{K(K+1)}{M}(\kappa - 1)\sigma_{jx}^4$$

Since $\hat{\sigma}_j{}^2 = K(K+1)\hat{\sigma}_{jx}{}^2$, the covariance between $\overline{\delta_{ij}^2}$ and $\hat{\sigma}_j{}^2$ can be derived as $\frac{K(K+1)}{M}(\kappa - 1)\sigma_{jx}^4 = (\kappa - 1)\sigma_j^4$. The variance of $\frac{\overline{\delta_{ij}^2} - \hat{\sigma}_j{}^2}{\hat{\sigma}_j{}^2}$ can be derived as

$$Var\left(\frac{\overline{\delta_{ij}^2}}{\hat{\sigma}_j{}^2}\right) \approx \frac{Var\left(\overline{\delta_{ij}^2}\right)}{\left(E\left[\hat{\sigma}_j{}^2\right]\right)^2} - \frac{2E\left[\overline{\delta_{ij}^2}\right]Cov\left(\overline{\delta_{ij}^2}, \hat{\sigma}_j{}^2\right)}{\left(E\left[\hat{\sigma}_j{}^2\right]\right)^3} + \frac{\left(E\left[\overline{\delta_{ij}^2}\right]\right)^2 Var\left(\hat{\sigma}_j{}^2\right)}{\left(E\left[\hat{\sigma}_j{}^2\right]\right)^4}$$

$$= \frac{1}{M}\left(\frac{\left(\left(\kappa_j - 1\right) + U\right)\sigma_j^4}{\sigma_j^4} - 2\,\frac{\sigma_j^2(\kappa - 1)\sigma_j^4}{\sigma_j^6} + \frac{\sigma_j^4(\kappa - 1)\sigma_j^4}{\sigma_j^8}\right) = \frac{U}{M}$$

where $U = \frac{4K^3}{K^2(K+1)^2} + \frac{2w_{n2} - 8Kw_{n3} + 4K^2w_{n4}}{MK^2(K+1)^2}$

**Supplementary Note 5**

For convenience, let $x_i = x_{ij} - \mu_{jx}$, $T_i = T_{ij}^{(K)} - K\mu_{jx}$, and $\delta_i = T_i - Kx_i = T_{ij}^{(K)} - x_{ij} = \delta_{ij}$. The expectation and variance of $x_i$ can be derived as $E[x_i] = 0$ and $Var(x_i) = E[x_i^2] = \sigma_{jx}^2$.

Since

$$Cov\left(T_{ij}^{(K)}, Kx_{ij}\right) = E\left[\left(T_{ij}^{(K)} - E\left[T_{ij}^{(K)}\right]\right)\left(Kx_{ij} - E\left[Kx_{ij}\right]\right)\right] = E[T_i K x_i] = Cov(T_i, Kx_i)$$

the effect size of $\eta_j^2$ can be rewritten as

$$\eta_j^2 = \frac{2\overline{Cov(T_{ij}^{(K)}, Kx_{ij})}}{\hat{\sigma}_j^{\,2}} = \frac{2\overline{Cov(T_i, Kx_i)}}{\hat{\sigma}_j^{\,2}}$$

Denote the dropout indicator $Q_i \sim Bernoulli(1-q)$ which is independent across $i$ and independent of $x$, where $q$ represents the dropout rate, and denote the observed expressions after dropout as $x' = Q_i x_i$. The covariance of a spot and its neighbor after dropout can be derived as

$$Cov\left(Q_{i_1}x_{i_1}, Q_{i_2}x_{i_2}\right) = E\left[Q_{i_1}Q_{i_2}x_{i_1}x_{i_2}\right] - E[Q_{i_1}x_{i_1}]E[Q_{i_2}x_{i_2}]$$

Since $Q_i$ is independent of $x$ and independent across spots,

$$E\left[Q_{i_1}Q_{i_2}x_{i_1}x_{i_2}\right] = E\left[Q_{i_1}\right]E\left[Q_{i_2}\right]E\left[x_{i_1}x_{i_2}\right] = (1-q)^2 E\left[x_{i_1}x_{i_2}\right]$$

and

$$E[Q_i x_i] = E[Q_i]E[x_i] = 0$$

Therefore for $i_1 \neq i_2$

$$Cov\left(x'_{i_1}, x'_{i_2}\right) = (1-q)^2 Cov\left(x_{i_1}, x_{i_2}\right)$$

and thus

$$\overline{Cov(T'_i, Kx'_i)} = (1-q)^2 \overline{Cov(T_i, Kx_i)}$$

where $T'$ denotes the observed value of $T$ after dropout. Since $E[x'_i] = E[Q_i x_i] = 0$, the variance of observed expressions after dropout can be derived as

$$Var(x_i') = E\left[{x_i'}^2\right] - (E[x_i'])^2 = E[Q_i^2 x_i^2]$$

Given that $Q_i \in \{0,1\}$, we have $Q_i^2 = Q_i$, and thus

$$Var(x_i') = E[Q_i^2 x_i^2] = E[Q_i x_i^2] = Q_i E[x_i^2] = (1-q)\sigma_{jx}^2$$

Therefore, the denominator after dropout is ${\hat{\sigma}_j'}^2 = (1-q){\hat{\sigma}_j}^2$. The effect size after dropout can be derived as

$$\eta_{j,drop}^2 = \frac{2\overline{Cov(T'_i, Kx'_i)}}{{\hat{\sigma}_j'}^2} = \frac{2(1-q)^2\overline{Cov(T_i, Kx_i)}}{(1-q)\sigma_{jx}^2} = (1-q)\eta_j^2$$

**Supplementary Note 6**

Consider an additional noise term, $\boldsymbol{\epsilon_j}$, with mean zero and variance $\tau_j^2$ in the expressions of each feature to account for the technically introduced variations. Denote the observed expression of feature $j$ at spot $i$ as $x'_{ij}$, we have

$$x'_{ij} = x_{ij} + \epsilon_{ij}$$

In this case, the mean and variance of $T_{ij}^{(K)}$ are $K\mu_{jx}$ and $K(\sigma_{jx}^2 + \tau_j^2)$ while the mean and variance of $Kx'_{ij}$ are $K\mu_{jx}$ and $K^2(\sigma_{jx}^2 + \tau_j^2)$. Since $T_{ij}^{(K)}$ and $x'_{ij}$ are still independent for a fixed constant of $K$, the expectation and variance of $\delta_{ij}$ can be derived as

$$E[\delta'_{ij}] = E\left[T_{ij}^{(K)}\right] - E[Kx'_{ij}] = K\mu_{jx} - K\mu_{jx} = 0$$

and

$$Var(\delta'_{ij}) = Var\left(T_{ij}^{(K)}\right) + Var(Kx'_{ij}) = K(\sigma_{jx}^2 + \tau_j^2) + K^2(\sigma_{jx}^2 + \tau_j^2) = K(K+1)(\sigma_{jx}^2 + \tau_j^2)$$

Let

$$\sigma_j'^2 = K(K+1)(\sigma_{jx}^2 + \tau_j^2)$$

and take it into Equation (4) in **Supplementary Note 1**, we have

$$Z = \sqrt{\frac{MK^2(K+1)^2}{4K^3 + \frac{2w_{n2} - 8Kw_{n3} + 4K^2w_{n4}}{M}}}\left(\frac{\overline{\delta_{ij}'^2}}{\hat{\sigma}'^2_j} - 1\right) \sim N(0,1)$$

Compared to the scenario without additional noise term, the variance of noise, $\tau_j^2$, is added to both the numerator and denominator of the test statistics which will result in an observed effect size as follows

$$\eta_j'^2 = 1 - \frac{\overline{\delta_{ij}'^2}}{\hat{\sigma}'^2_j} = \frac{\eta_j^2}{1+R}$$

where $R = \frac{\tau_j^2}{\sigma_{jx}^2}$.

**Supplementary Note 7**

Assume the expression of a feature $j$ follows a negative binomial distribution. Under the null hypothesis, the expression at each spot is independent of each other, i.e., $x_{1j}, \dots, x_{Mj}\ i.i.d. \sim NB(r_j, p_j)$. For a spot $i$, the distribution of the sum of its neighboring spots follows a negative binomial distribution as well, i.e., $T_{ij}^{(K)} \sim NB(Kr_j, p_j)$. Denote the difference between $T_{ij}^{(K)}$ and $Kx_{ij}$ as $\delta_{ij}$. Since $T_{ij}^{(K)}$ and $x_{ij}$ are independent for a fixed constant of $K$ at spot $i$, $\delta_{ij}$ is the difference between two (scaled) negative binomial distribution. The expectation and variance of $\delta_{ij}$ can be derived as

$$E[\delta_{ij}] = E\left[T_{ij}^{(K)}\right] - E[Kx_{ij}] = \frac{Kr_j(1-p_j)}{p_j} - K\frac{r_j(1-p_j)}{p_j} = 0$$

and

$$Var(\delta_{ij}) = Var\left(T_{ij}^{(K)}\right) + Var(Kx_{ij}) = \frac{Kr_j(1-p_j)}{p_j^2} + K^2\frac{r_j(1-p_j)}{p_j^2} = \frac{K(K+1)r_j(1-p_j)}{p_j^2}$$

with the probability mass function of

$$P(\delta = d) = \sum_{t=0}^{\infty}\left(\binom{d+tK+Kr_j-1}{d+tK}(1-p_j)^{d+tK}p_j^{Kr_j}\binom{t+r_j-1}{t}(1-p_j)^t p^r\right)$$

For a large K and large r, the distribution of $T_{ij}^{(K)}$ and $Kx_{ij}$tends to approximate a normal distribution due to the CLT. The distribution of $\delta_{ij}$ can be approximated as

$$\delta_{ij} \sim N\left(0, K(K+1)\sigma_{jx}^2\right)$$

where $\sigma_{jx}^2 = \frac{r_j(1-p_j)}{p_j^2}$ is the variance of $x_j$. Therefore, $\boldsymbol{\delta_j} = \left(\delta_{1j}, \dots \delta_{Mj}\right)^T$ follows a multivariate normal distribution as follows:

$$\boldsymbol{\delta_j} \sim \boldsymbol{N}(\boldsymbol{0}, \sigma_{jx}^2\boldsymbol{\Sigma})$$

where $\sigma_{jx}^2\boldsymbol{\Sigma}$ is the covariance matrix of $\boldsymbol{\delta_j}$. Considering the $K^2$-dependent sequence $\boldsymbol{\delta_j^2} = \left(\delta_{1j}^2, \dots \delta_{Mj}^2\right)^T$ with expectation and variance of each term as $E\left[\delta_{ij}^2\right] = \sigma_j^2$ and $Var\left(\delta_{ij}^2\right) = E\left[\delta_{ij}^4\right] - \left(E\left[\delta_{ij}^2\right]\right)^2 = 3\sigma_j^4 - \sigma_j^4 = 2\sigma_j^4$ (since the kurtosis of normally distributed random variables is 3) where

$$\sigma_j^2 = \frac{K(K+1)r_j\left(1-p_j\right)}{p_j^2} = K(K+1)\sigma_{jx}^2 \quad (5)$$

Based on the CLT for $K^2$-dependent sequence, we have

$$\sqrt{M}\left(\overline{\delta_{ij}^2} - \sigma_j^2\right) \sim N\left(0, 2\sigma_j^4 + 2\sum_{k=1}^{K^2} Cov\left(\delta_{1j}^2, \delta_{(1+k)j}^2\right)\right) \quad (6)$$

Since $\delta_{ij}$ follows a normal distribution, their covariance of squared terms has property (**Supplementary Note 8**):

$$Cov\left(\delta_{1j}^2, \delta_{(1+k)j}^2\right) = 2Cov^2\left(\delta_{1j}, \delta_{(1+k)j}\right) = 2\Sigma_{1(1+k)}^2\sigma_{jx}^4$$

Using the combinatorial arguments and approximate symmetry of the covariance matrix, the total sum of $\sum_{1 \le i_1 \le M}\left(\Sigma_{\mathrm{i}_1 i_2}\right)^2$ for each $i$ is $K(K+1)^2$ since the total number of neighboring pairs are $K$ and each pair contributes approximately $(K+1)^2$ to the sum due to the squaring operation. Therefore,

$$2\sum_{k=1}^{K} Cov\left(\delta_{1j}^2, \delta_{(1+k)j}^2\right) = 2\sum_{1 \le i \le M} 2(\Sigma_{1\mathrm{i}})^2 = 4K(K+1)^2\sigma_{jx}^4 = \frac{4}{K}\sigma_j^4$$

Implementing this and Equation (5), Equation (6) can be rewritten as

$$\sqrt{M}\left(\overline{\delta_{ij}^2} - \sigma_j^2\right) \sim N\left(0, \left(2 + \frac{4}{K}\right)\sigma_j^4\right)$$

Since $\sigma_j^2$ can be estimated from the sample as ${\hat{\sigma}_j}^2 = K(K+1){\hat{\sigma}_{jx}}^2$ which is greater than 0 and finite, the variance of $(\overline{\delta_{ij}^2} - {\hat{\sigma}_j}^2)/{\hat{\sigma}_j}^2$ can be derived from the first order of Taylor expansion as

$$Var\left(\frac{\overline{\delta_{ij}^2} - \hat{\sigma}_j{}^2}{\hat{\sigma}_j{}^2}\right) \approx \frac{Var\left(\overline{\delta_{ij}^2}\right)}{\left(E\left[\hat{\sigma}_j{}^2\right]\right)^2} - \frac{2E\left[\overline{\delta_{ij}^2}\right]Cov\left(\overline{\delta_{ij}^2}, \hat{\sigma}_j{}^2\right)}{\left(E\left[\hat{\sigma}_j{}^2\right]\right)^3} + \frac{\left(E\left[\overline{\delta_{ij}^2}\right]\right)^2 Var\left(\hat{\sigma}_j{}^2\right)}{\left(E\left[\hat{\sigma}_j{}^2\right]\right)^4}$$

which is calculated $\frac{4}{MK}$ (**Supplementary Note 4**). Therefore, the test statistics can be converted to the standard normal form

$$Z = \sqrt{\frac{KM}{4}}\left(\frac{\overline{\delta_{ij}^2}}{\hat{\sigma}_j{}^2} - 1\right) \sim N(0,1) \qquad (7)$$

The null hypothesis is equivalent to

$$H_0\colon \frac{\hat{\sigma}_{j1}{}^2}{\hat{\sigma}_j{}^2} = 1$$

The sample variance $\hat{\sigma}_{j1}{}^2$ can be approximated with $\overline{\delta_{ij}^2}$ for a large M since $E\left[\delta_{ij}\right] = 0$. For a two-sided test using the test statistics of $\frac{\overline{\delta_{ij}^2}}{\hat{\sigma}_j{}^2} - 1$ in Equation (7), the probability of rejecting null hypothesis is

$$P = 2Prob\left(Z > \left|\sqrt{\frac{KM}{4}}\left(\frac{\overline{\delta_{ij}^2}}{\hat{\sigma}_j{}^2} - 1\right)\right|\right)$$

This approximation relies on the multivariate normality of $\boldsymbol{\delta_j^2}$. When the expression of a feature *j* follows a normal approximated distribution, e.g., a Poisson distribution with large $\lambda$, Equation (6) also holds and thereby the null hypothesis can be tested with Equation (7).

**Supplementary Note 8**

Assume there are two dependent and identically distributed random normal variables, $X$ and $Y$, with mean 0 and variance of $\sigma^2$, and the covariance between $X$ and $Y$ is $Cov(X, Y) = \rho\sigma^2$. Denoting the kurtosis of $X$ and $Y$ as $\kappa = \frac{E[X^4]}{(E[X^2])^2} = \frac{E[X^4]}{\sigma^4}$, we have $E[X^4] = \kappa\sigma^4 = 3\sigma^4$. Based on the definition, the covariance between $X$ and $Y$ is

$$Cov(X^2, Y^2) = E[X^2Y^2] - E[X^2][Y^2] = E[X^2Y^2] - \sigma^4$$

For the first term, we express $Y$ in terms of the linear combination of $X$ and an orthogonal component:

$$Y = \rho X + \sqrt{1-\rho^2}Z$$

where $Z$ is a normally distributed random variable independent of $X$, with $E[Z] = 0$ and $Var(Z) = \sigma^2$. Therefore, the squared term can be rewritten as

$$X^2Y^2 = X^2(\rho^2X^2 + 2\rho\sqrt{1-\rho^2}XZ + (1-\rho)Z^2)$$

Taking expectations on both sides

$$E[X^2Y^2] = \rho^2E[X^4] + 2\rho\sqrt{1-\rho^2}E[X^3Z] + (1-\rho^2)E[X^2Z^2] = \rho^2 3\sigma^4 + (1-\rho^2)\sigma^4$$

since $E[X^3Z] = E[X^3]E[Z] = 0$. The covariance between $X^2$ and $Y^2$ can be expressed as

$$Cov(X^2, Y^2) = \rho^2 3\sigma^4 + (1-\rho^2)\sigma^4 - \sigma^4 = \rho^2(3-1)\sigma^4$$

which can be simplified as

$$Cov(X^2, Y^2) = 2Cov^2(X, Y)$$

**Supplementary Note 9**

**Detection of Cell-type-related SVF**:

Unlike cellular or subcellular resolution data, where differences in feature expression between various cell types can be directly tested (e.g., using differentially expressed gene analysis), low-resolution data contain spots composed of multiple cell types. For low-resolution data with $Q$ types of cells, let $\boldsymbol{c_q} = \left(c_{q1}, \dots, c_{qM}\right)^T$ denote the cell type proportions in spot $i$ where $c_{qM}$ represents the proportion of cell type $q$ and $\sum_{q=1}^{Q} c_{qM} = 1$, which can be estimated using cell type deconvolution tools such as RCDT, For each spot in the sample, we define a patch as the set of $K$ nearest neighboring spots using $\boldsymbol{C} = \left(\boldsymbol{c_1}, \dots, \boldsymbol{c_Q}\right)$ as the coordinate matrix. For each spot in the cell type proportion space, the $K$ nearest neighboring spots is determined, denoted as a binary M-vector $\boldsymbol{b}_{\boldsymbol{i_0}}^{(\boldsymbol{K})} = \left(b_{i_01}^{(K)}, \dots, b_{i_0M}^{(K)}\right)$, where $b_{i_0i}^{(K)} = 1$ if spot $i$ is within the $K$ nearest neighboring spots of spot $i_0$. For feature $j$ at spot $i_0$, the sum of the expressions of all spots within the patch, $\tilde{T}_{i_0j}^{(K)}$, is computed as $\tilde{T}_{i_0j}^{(K)} = \tilde{\boldsymbol{b}}_{\boldsymbol{i_0}}^{(\boldsymbol{K})}\boldsymbol{x_j}$. The *null hypothesis* states that a feature is not a cell-type-related spatially variable feature, i.e., its expression does not depend on cell type proportions. This hypothesis can be tested using Equations (4) and Equation (7) by replacing $T_{ij}^{(K)}$ by $\tilde{T}_{ij}^{(K)}$.

**Detection of Cell-type-specific SVF**:

For cellular or subcellular resolution data, cell-type-specific SVFs can be identified by applying SVF detection separately to each cell population. For low-resolution data, we adopt a regression model from the published literature under the null hypothesis that no cell-type-specific spatial pattern exists:

$$\boldsymbol{X} = \boldsymbol{C}\boldsymbol{V} + \boldsymbol{e}$$

where $\boldsymbol{C}$ is the cell type proportion matrix, which can be estimated using cell type deconvolution tools, $\boldsymbol{V}$ is the cell-type-specific expression matrix, and $\boldsymbol{e}$ de is the residual matrix with mean 0 and variance of $\sigma_{ej}^2$ for each feature.

To extend this approach to detecting cell-type-specific spatial patterns, we first compute the adjusted expression matrix for each cell type as follows

$$\boldsymbol{X_q} = \boldsymbol{c_q V_q}$$

where $\boldsymbol{X_q}$ is an $M \times N$ matrix represents the estimated expression per spot after adjusting for cell type proportions, $\boldsymbol{c_q}$ is the $M \times 1$ cell type proportion matrix, and $\boldsymbol{V_q}$ is a $1 \times N$ cell-type-specific expression matrix. When the cell type proportion matrix $\boldsymbol{C}$ has full column rank, the least squares estimate of $\boldsymbol{V} = \left(\boldsymbol{V}_1^T, \dots \boldsymbol{V}_q^T\right)^T$ is given by:

$$\widehat{\boldsymbol{V}} = (\boldsymbol{C}^T\boldsymbol{C})^{-1}\boldsymbol{C}^T\boldsymbol{X}$$

To reduce noise from spots with very small proportions of a given cell type, we exclude spots where the proportion falls below a predefined threshold $\beta_{j1}$ when estimating $\widehat{\boldsymbol{V}}$. The cell-type-specific SVF can be inferred from Equation (4) and Equation (7) by applying SVF detection on the expression matrix, $\boldsymbol{X_q}$, which has been adjusted for cell type proportions, using the coordinate matrix while excluding spots with proportions below the threshold.

**Supplementary Note 10**

Four three-dimensional spatial patterns were considered, including three continuous patterns (Curved Cell Strand, Tissue Layer, and Irregular Cell Aggregate) and one discrete pattern (Isolated Cell Nodules). These patterns were designed to mimic biologically plausible spatial organizations such as strand-like cell arrangements, tissue layers, dense aggregates, and isolated spherical clusters. The simulated spatial data consisted of 10 segments along the z-axis. Each segment contained 225 spatial locations arranged on a two-dimensional grid in the x–y plane. The design mimicked cryo-sectioned tissue, where each section was placed on an individual array without physical contact between array surfaces. Spatial coordinates were therefore defined in three dimensions, with z indexing the section.

For continuous patterns, spatial structures were generated using a branch of spheres whose centers were produced via a random walk process with a fixed step length of 2 spatial units. The three continuous patterns differed in directional constraints applied to the random walk. For the Curved Cell Strand pattern, the center trajectory was constrained to be monotonic in two spatial directions. For the Tissue Layer pattern, monotonicity was imposed in one direction. For the Irregular Cell Aggregate pattern, no monotonic constraint was imposed, resulting in non-monotonic cell clusters in all directions. The union of spheres defined the patterned region. For the discrete pattern (Isolated Cell Nodules), nonadjacent spheres were generated with centers spaced 8 spatial units apart. Four center points were selected with a fixed z-coordinate of 5.5. The x- and y-coordinates were selected from the sequence 3, 11, and 15 with spacing of 8 units as applicable. To introduce spatial variability, independent uniform noise ranging from −2 to 2 was added to each coordinate of the sphere centers.

In each simulation, feature expression values were assigned according to whether a spatial location was inside or outside the pattern. Cells located inside the patterned region were labeled as marked. For each true SVF, expression values for marked cells were randomly sampled from the upper quantile of the empirical feature expression distribution derived from seqFISH data. For unmarked cells and all cells outside the pattern, expression values were randomly sampled from the full seqFISH distribution. Null features were generated by permuting the expression values of SVFs, thereby preserving marginal distributions while removing spatial structure. Additional Gaussian noise was added to all features proportionally to the average standard deviation of expression across features. Three simulation parameters were varied systematically. Pattern size was defined by the diameter of the spheres and set to 2.5, 3.0, and 3.5 units representing small, moderate, and large patterns. Signal strength was defined as the fold change between the average expression inside and outside the pattern and was set to 2.0, 2.5, and 3.0 representing low, moderate, and high signal strength. Noise level was defined as a proportion of the averaged standard deviation of expression across features and was set to 20%, 50%, and 80% representing low, moderate, and high noise. For each combination of spatial pattern, pattern size, signal strength, and noise level, 1,000 true SVFs were simulated together with 9,000 null features generated by permutation and ten independent replicates were generated to mitigate sampling variability.
