## Supplementary Tables for "A Principled Statistical Framework for Analyzing Spatial Patterns in Spatially Resolved Multi-Omics"

AUPRC

| Dataset | SPARK | SPARK.X | STORM | STORM_A | nnSVG | Moran's I | SOMDE | SpatialDE |
|---|---|---|---|---|---|---|---|---|
| Biancalani_ | 0.978117 | 0.960044 | 0.982777 | 0.974813 | 0.962636 | 0.889241 | 0.970885 | 0.973938 |
| Ferreira_et | 0.998378 | 0.999531 | 0.971329 | 1 | 0.999502 | 0.982648 | 0.995816 | 1 |
| Janosevic_ | 1 | 0.998839 | 1 | 0.998398 | 0.999245 | 0.996019 | 0.97835 | 1 |
| Joglekar_e | 0.35815 | 1 | 1 | 1 | 0.993631 | 1 | 0.992571 | 0.99779 |
| Liu_et_al_l | 1 | 0.988853 | 1 | 0.999541 | 0.996719 | 0.960063 | 0.990005 | 0.999959 |
| Liu_et_al_l | 1 | 0.998895 | 1 | 1 | 1 | 0.987922 | 0.998719 | 1 |
| Lopez_et_a | 0.996165 | 1 | 1 | 1 | 1 | 1 | 0.995742 | 0.999567 |
| Wu_et_al_ | 0.985129 | 0.987525 | 1 | 0.970395 | 0.972691 | 0.914658 | 0.988657 | 0.995786 |
| Wu_et_al_ | 0.995068 | 0.957151 | 0.977347 | 0.977258 | 0.963999 | 0.882906 | 0.965399 | 0.984723 |

AUC

| Dataset | SPARK | SPARK.X | STORM | STORM_A | nnSVG | Moran's I | SOMDE | SpatialDE |
|---|---|---|---|---|---|---|---|---|
| Biancalani_ | 0.990773 | 0.98094 | 0.993813 | 0.985766 | 0.97617 | 0.925576 | 0.990161 | 0.98057 |
| Ferreira_et | 0.99977 | 0.999947 | 0.98321 | 1 | 0.999941 | 0.990657 | 0.999297 | 1 |
| Janosevic_ | 1 | 0.999857 | 1 | 0.999775 | 0.999907 | 0.998988 | 0.995811 | 1 |
| Joglekar_e | 0.895577 | 1 | 1 | 1 | 0.999644 | 1 | 0.998681 | 0.999658 |
| Liu_et_al_l | 1 | 0.997845 | 1 | 0.999945 | 0.999484 | 0.985951 | 0.998133 | 0.999995 |
| Liu_et_al_l | 1 | 0.99987 | 1 | 1 | 1 | 0.992907 | 0.999826 | 1 |
| Lopez_et_a | 0.999194 | 1 | 1 | 1 | 1 | 1 | 0.999474 | 0.99995 |
| Wu_et_al_ | 0.983623 | 0.997397 | 1 | 0.983448 | 0.98615 | 0.942347 | 0.995061 | 0.995811 |
| Wu_et_al_ | 0.999374 | 0.982727 | 0.986926 | 0.992607 | 0.982617 | 0.932931 | 0.990872 | 0.986113 |

KS distance

| Dataset | spark | SPARKX | storm | storm_App | nnsvg | MoranI | SOMDE | spatialde | scBSP |
|---|---|---|---|---|---|---|---|---|---|
| Biancalani_ | 0.035677 | 0.039814 | 0.04395 | 0.024302 | 0.602379 | 0.500258 | 0.452762 | 0.430047 | 0.104447 |
| Ferreira_et | 0.037873 | 0.062637 | 0.027677 | 0.018937 | 0.584851 | 0.501093 | 0.421704 | 0.420976 | 0.094683 |
| Janosevic_ | 0.046381 | 0.037948 | 0.023893 | 0.040056 | 0.562193 | 0.500351 | 0.455376 | 0.434997 | 0.095573 |
| Joglekar_e | 0.03075 | 0.05189 | 0.011531 | 0.016015 | 0.552851 | 0.50032 | 0.352979 | 0.625881 | 0.099936 |
| Liu_et_al_l | 0.030769 | 0.03141 | 0.023077 | 0.040385 | 0.582692 | 0.500641 | 0.460897 | 0.419231 | 0.117308 |
| Liu_et_al_l | 0.047559 | 0.031915 | 0.040676 | 0.055695 | 0.548185 | 0.5 | 0.496871 | 0.4199 | 0.107635 |
| Lopez_et_a | 0.033762 | 0.047428 | 0.022508 | 0.03537 | 0.56672 | 0.500804 | 0.440515 | 0.428457 | 0.146302 |
| Wu_et_al_ | 0.04063 | 0.024046 | 0.022388 | 0.041459 | 0.567164 | 0.500414 | 0.472245 | 0.787904 | 0.151741 |

| | | | | | | | | | |
|---|---|---|---|---|---|---|---|---|---|
| Wu_et_al_ | 0.039295 | 0.028455 | 0.01897 | 0.021003 | 0.560976 | 0.500678 | 0.361111 | 0.5 | 0.123306 |

| Scenario | Spatial Pat | Parameter | STORM | STORM (Approximated) |
|---|---|---|---|---|
| scenario1 | RW1 | 1.25 | 0 | 0 |
| scenario1 | RW1 | 1.5 | 0 | 0 |
| scenario1 | RW1 | 1.75 | 0 | 0 |
| scenario1 | RW2 | 1.25 | 0 | 0 |
| scenario1 | RW2 | 1.5 | 0 | 0 |
| scenario1 | RW2 | 1.75 | 0 | 0 |
| scenario1 | RW3 | 1.25 | 0 | 0 |
| scenario1 | RW3 | 1.5 | 0 | 0 |
| scenario1 | RW3 | 1.75 | 0 | 0 |
| scenario1 | RW4 | 1.25 | 2.9429086 | 0 |
| scenario1 | RW4 | 1.5 | 0 | 0 |
| scenario1 | RW4 | 1.75 | 0 | 0 |
| scenario2 | RW1 | 2.5 | 0 | 0 |
| scenario2 | RW1 | 2 | 0 | 0 |
| scenario2 | RW1 | 3 | 0 | 0 |
| scenario2 | RW2 | 2.5 | 0 | 0 |
| scenario2 | RW2 | 2 | 0 | 0 |
| scenario2 | RW2 | 3 | 0 | 0 |
| scenario2 | RW3 | 2.5 | 0 | 0 |
| scenario2 | RW3 | 2 | 0 | 0 |
| scenario2 | RW3 | 3 | 0 | 0 |
| scenario2 | RW4 | 2.5 | 0 | 0 |
| scenario2 | RW4 | 2 | 0 | 0 |
| scenario2 | RW4 | 3 | 0 | 0 |
| scenario3 | RW1 | 0.2 | 0 | 0 |
| scenario3 | RW1 | 0.5 | 0 | 0 |
| scenario3 | RW1 | 0.8 | 0 | 0 |
| scenario3 | RW2 | 0.2 | 0 | 0 |
| scenario3 | RW2 | 0.5 | 0 | 0 |
| scenario3 | RW2 | 0.8 | 0 | 0 |
| scenario3 | RW3 | 0.2 | 0 | 0 |
| scenario3 | RW3 | 0.5 | 0 | 0 |
| scenario3 | RW3 | 0.8 | 0 | 0 |
| scenario3 | RW4 | 0.2 | 0 | 0 |
| scenario3 | RW4 | 0.5 | 0 | 0 |
| scenario3 | RW4 | 0.8 | 0 | 0 |

| Scenario | Spatial Pat | Parameter | Parameter | STORM | STORM (Approximated) |
|---|---|---|---|---|---|
| scenario1 | RW1 | 1.25 | 1.5 | 1.3086255 | 9.18709112579905e-115 |
| scenario1 | RW1 | 1.25 | 1.75 | 0 | 0 |
| scenario1 | RW1 | 1.5 | 1.75 | 6.6188388 | 6.08673398647494e-214 |
| scenario1 | RW2 | 1.25 | 1.5 | 1.0967560 | 3.20611879606319e-171 |
| scenario1 | RW2 | 1.25 | 1.75 | 0 | 0 |
| scenario1 | RW2 | 1.5 | 1.75 | 2.2183472 | 2.01217903177341e-132 |
| scenario1 | RW3 | 1.25 | 1.5 | 4.6738920 | 3.30920220388628e-251 |
| scenario1 | RW3 | 1.25 | 1.75 | 0 | 0 |
| scenario1 | RW3 | 1.5 | 1.75 | 2.7910369 | 1.64670849439645e-262 |
| scenario1 | RW4 | 1.25 | 1.5 | 4.5488397 | 1.66247481628509e-153 |
| scenario1 | RW4 | 1.25 | 1.75 | 0 | 0 |
| scenario1 | RW4 | 1.5 | 1.75 | 5.4210937 | 1.09142004109379e-179 |
| scenario2 | RW1 | 2.5 | 2 | 1.0120834 | 4.67335988050744e-143 |
| scenario2 | RW1 | 2.5 | 3 | 4.3734506 | 4.61040563434879e-174 |
| scenario2 | RW1 | 2 | 3 | 0 | 0 |
| scenario2 | RW2 | 2.5 | 2 | 1.8293949 | 7.11031488370053e-240 |
| scenario2 | RW2 | 2.5 | 3 | 1.1371428 | 6.35625888344644e-264 |
| scenario2 | RW2 | 2 | 3 | 0 | 0 |
| scenario2 | RW3 | 2.5 | 2 | 0 | 0 |
| scenario2 | RW3 | 2.5 | 3 | 2.2960013 | 6.76431500948388e-316 |
| scenario2 | RW3 | 2 | 3 | 0 | 0 |
| scenario2 | RW4 | 2.5 | 2 | 3.7063133 | 2.48702108866523e-117 |
| scenario2 | RW4 | 2.5 | 3 | 1.1750644 | 8.44142967618716e-133 |
| scenario2 | RW4 | 2 | 3 | 5.3856903 | 0 |
| scenario3 | RW1 | 0.2 | 0.5 | 5.1122399 | 7.61950990010379e-16 |
| scenario3 | RW1 | 0.2 | 0.8 | 9.5220047 | 2.44792174102923e-71 |
| scenario3 | RW1 | 0.5 | 0.8 | 1.3803746 | 4.31697657320596e-25 |
| scenario3 | RW2 | 0.2 | 0.5 | 6.7098397 | 1.45074538613161e-26 |
| scenario3 | RW2 | 0.2 | 0.8 | 3.3423780 | 2.8083276076156e-121 |
| scenario3 | RW2 | 0.5 | 0.8 | 2.0802652 | 2.93875579361936e-44 |
| scenario3 | RW3 | 0.2 | 0.5 | 4.0683710 | 1.01706933195479e-36 |
| scenario3 | RW3 | 0.2 | 0.8 | 1.7257229 | 1.32398803567771e-165 |
| scenario3 | RW3 | 0.5 | 0.8 | 3.9811008 | 7.89314549694057e-63 |
| scenario3 | RW4 | 0.2 | 0.5 | 1.3949118 | 6.81743696058874e-12 |
| scenario3 | RW4 | 0.2 | 0.8 | 3.4201970 | 1.14151627515147e-53 |
| scenario3 | RW4 | 0.5 | 0.8 | 2.2419543 | 4.80415208565492e-19 |

| Scenario | Spatial Pat | Parameter | Sample.Si: | Estimated. | Observed.Power |
|---|---|---|---|---|---|
| scenario1 | RW1 | 1.25 | 3 | 0.34069 | 0.415049 |
| scenario1 | RW1 | 1.25 | 4 | 0.465407 | 0.181425 |
| scenario1 | RW1 | 1.25 | 5 | 0.564767 | 0.289144 |
| scenario1 | RW1 | 1.25 | 6 | 0.639593 | 0.401026 |
| scenario1 | RW1 | 1.25 | 7 | 0.698456 | 0.534553 |
| scenario1 | RW1 | 1.25 | 8 | 0.616671 | 0.555444 |
| scenario1 | RW1 | 1.25 | 9 | 0.689602 | 0.61407 |
| scenario1 | RW1 | 1.25 | 10 | 0.747635 | 0.673674 |
| scenario1 | RW1 | 1.25 | 11 | 0.719436 | 0.702703 |
| scenario1 | RW1 | 1.25 | 12 | 0.763467 | 0.764 |
| scenario1 | RW1 | 1.25 | 13 | 0.802096 | 0.803 |
| scenario1 | RW1 | 1.25 | 14 | 0.835257 | 0.835 |
| scenario1 | RW1 | 1.25 | 15 | 0.858024 | 0.872 |
| scenario1 | RW1 | 1.25 | 16 | 0.881128 | 0.859 |
| scenario1 | RW1 | 1.25 | 17 | 0.900525 | 0.876 |
| scenario1 | RW1 | 1.25 | 18 | 0.916752 | 0.898 |
| scenario1 | RW1 | 1.25 | 19 | 0.930312 | 0.908 |
| scenario1 | RW1 | 1.25 | 20 | 0.922657 | 0.928 |
| scenario1 | RW1 | 1.5 | 3 | 0.920642 | 0.921844 |
| scenario1 | RW1 | 1.5 | 4 | 0.9241 | 0.886 |
| scenario1 | RW1 | 1.5 | 5 | 0.968202 | 0.965 |
| scenario1 | RW1 | 1.5 | 6 | 0.989139 | 0.991 |
| scenario1 | RW1 | 1.5 | 7 | 0.996597 | 0.986 |
| scenario1 | RW1 | 1.5 | 8 | 0.995662 | 0.998 |
| scenario1 | RW1 | 1.5 | 9 | 0.998379 | 1 |
| scenario1 | RW1 | 1.5 | 10 | 0.999448 | 1 |
| scenario1 | RW1 | 1.5 | 11 | 0.999806 | 1 |
| scenario1 | RW1 | 1.5 | 12 | 0.999942 | 1 |
| scenario1 | RW1 | 1.5 | 13 | 0.999983 | 1 |
| scenario1 | RW1 | 1.5 | 14 | 0.999979 | 1 |
| scenario1 | RW1 | 1.5 | 15 | 0.99999 | 1 |
| scenario1 | RW1 | 1.5 | 16 | 0.999997 | 1 |
| scenario1 | RW1 | 1.5 | 17 | 0.999999 | 1 |
| scenario1 | RW1 | 1.5 | 18 | 1 | 1 |
| scenario1 | RW1 | 1.5 | 19 | 1 | 1 |
| scenario1 | RW1 | 1.5 | 20 | 1 | 1 |
| scenario1 | RW1 | 1.75 | 3 | 0.991512 | 0.994 |
| scenario1 | RW1 | 1.75 | 4 | 0.997308 | 0.978 |
| scenario1 | RW1 | 1.75 | 5 | 0.999615 | 0.998 |
| scenario1 | RW1 | 1.75 | 6 | 0.999954 | 0.999 |
| scenario1 | RW1 | 1.75 | 7 | 0.999995 | 1 |
| scenario1 | RW1 | 1.75 | 8 | 0.999998 | 1 |
| scenario1 | RW1 | 1.75 | 9 | 1 | 1 |
| scenario1 | RW1 | 1.75 | 10 | 1 | 1 |
| scenario1 | RW1 | 1.75 | 11 | 1 | 1 |
| scenario1 | RW1 | 1.75 | 12 | 1 | 1 |
| scenario1 | RW1 | 1.75 | 13 | 1 | 1 |
| scenario1 | RW1 | 1.75 | 14 | 1 | 1 |
| scenario1 | RW1 | 1.75 | 15 | 1 | 1 |

| | | | | | |
|---|---|---|---|---|---|
| scenario1 | RW1 | 1.75 | 16 | 1 | 1 |
| scenario1 | RW1 | 1.75 | 17 | 1 | 1 |
| scenario1 | RW1 | 1.75 | 18 | 1 | 1 |
| scenario1 | RW1 | 1.75 | 19 | 1 | 1 |
| scenario1 | RW1 | 1.75 | 20 | 1 | 1 |
| scenario1 | RW2 | 1.25 | 3 | 0.810848 | 0.79598 |
| scenario1 | RW2 | 1.25 | 4 | 0.851128 | 0.707 |
| scenario1 | RW2 | 1.25 | 5 | 0.912132 | 0.838 |
| scenario1 | RW2 | 1.25 | 6 | 0.953609 | 0.936 |
| scenario1 | RW2 | 1.25 | 7 | 0.977202 | 0.96 |
| scenario1 | RW2 | 1.25 | 8 | 0.977144 | 0.953 |
| scenario1 | RW2 | 1.25 | 9 | 0.987395 | 0.978 |
| scenario1 | RW2 | 1.25 | 10 | 0.993325 | 0.986 |
| scenario1 | RW2 | 1.25 | 11 | 0.995492 | 0.996 |
| scenario1 | RW2 | 1.25 | 12 | 0.997949 | 0.996 |
| scenario1 | RW2 | 1.25 | 13 | 0.999081 | 0.998 |
| scenario1 | RW2 | 1.25 | 14 | 0.999354 | 0.998 |
| scenario1 | RW2 | 1.25 | 15 | 0.999448 | 1 |
| scenario1 | RW2 | 1.25 | 16 | 0.999724 | 1 |
| scenario1 | RW2 | 1.25 | 17 | 0.999865 | 0.999 |
| scenario1 | RW2 | 1.25 | 18 | 0.999936 | 1 |
| scenario1 | RW2 | 1.25 | 19 | 0.99997 | 1 |
| scenario1 | RW2 | 1.25 | 20 | 0.999984 | 1 |
| scenario1 | RW2 | 1.5 | 3 | 0.990665 | 0.988 |
| scenario1 | RW2 | 1.5 | 4 | 0.995846 | 0.982 |
| scenario1 | RW2 | 1.5 | 5 | 0.999361 | 0.996 |
| scenario1 | RW2 | 1.5 | 6 | 0.99992 | 1 |
| scenario1 | RW2 | 1.5 | 7 | 0.999991 | 1 |
| scenario1 | RW2 | 1.5 | 8 | 0.999995 | 1 |
| scenario1 | RW2 | 1.5 | 9 | 0.999999 | 1 |
| scenario1 | RW2 | 1.5 | 10 | 1 | 1 |
| scenario1 | RW2 | 1.5 | 11 | 1 | 1 |
| scenario1 | RW2 | 1.5 | 12 | 1 | 1 |
| scenario1 | RW2 | 1.5 | 13 | 1 | 1 |
| scenario1 | RW2 | 1.5 | 14 | 1 | 1 |
| scenario1 | RW2 | 1.5 | 15 | 1 | 1 |
| scenario1 | RW2 | 1.5 | 16 | 1 | 1 |
| scenario1 | RW2 | 1.5 | 17 | 1 | 1 |
| scenario1 | RW2 | 1.5 | 18 | 1 | 1 |
| scenario1 | RW2 | 1.5 | 19 | 1 | 1 |
| scenario1 | RW2 | 1.5 | 20 | 1 | 1 |
| scenario1 | RW2 | 1.75 | 3 | 0.99275 | 0.991 |
| scenario1 | RW2 | 1.75 | 4 | 0.999519 | 0.986 |
| scenario1 | RW2 | 1.75 | 5 | 0.99997 | 1 |
| scenario1 | RW2 | 1.75 | 6 | 0.999998 | 1 |
| scenario1 | RW2 | 1.75 | 7 | 1 | 1 |
| scenario1 | RW2 | 1.75 | 8 | 1 | 1 |
| scenario1 | RW2 | 1.75 | 9 | 1 | 1 |
| scenario1 | RW2 | 1.75 | 10 | 1 | 1 |
| scenario1 | RW2 | 1.75 | 11 | 1 | 1 |

| | | | | | |
|---|---|---|---|---|---|
| scenario1 | RW2 | 1.75 | 12 | 1 | 1 |
| scenario1 | RW2 | 1.75 | 13 | 1 | 1 |
| scenario1 | RW2 | 1.75 | 14 | 1 | 1 |
| scenario1 | RW2 | 1.75 | 15 | 1 | 1 |
| scenario1 | RW2 | 1.75 | 16 | 1 | 1 |
| scenario1 | RW2 | 1.75 | 17 | 1 | 1 |
| scenario1 | RW2 | 1.75 | 18 | 1 | 1 |
| scenario1 | RW2 | 1.75 | 19 | 1 | 1 |
| scenario1 | RW2 | 1.75 | 20 | 1 | 1 |
| scenario1 | RW3 | 1.25 | 3 | 0.919802 | 0.922 |
| scenario1 | RW3 | 1.25 | 4 | 0.923473 | 0.874 |
| scenario1 | RW3 | 1.25 | 5 | 0.967789 | 0.97 |
| scenario1 | RW3 | 1.25 | 6 | 0.988937 | 0.993 |
| scenario1 | RW3 | 1.25 | 7 | 0.996515 | 0.989 |
| scenario1 | RW3 | 1.25 | 8 | 0.995572 | 0.999 |
| scenario1 | RW3 | 1.25 | 9 | 0.998336 | 1 |
| scenario1 | RW3 | 1.25 | 10 | 0.99943 | 1 |
| scenario1 | RW3 | 1.25 | 11 | 0.999798 | 1 |
| scenario1 | RW3 | 1.25 | 12 | 0.999939 | 1 |
| scenario1 | RW3 | 1.25 | 13 | 0.999982 | 1 |
| scenario1 | RW3 | 1.25 | 14 | 0.999978 | 1 |
| scenario1 | RW3 | 1.25 | 15 | 0.999989 | 1 |
| scenario1 | RW3 | 1.25 | 16 | 0.999997 | 1 |
| scenario1 | RW3 | 1.25 | 17 | 0.999999 | 1 |
| scenario1 | RW3 | 1.25 | 18 | 1 | 1 |
| scenario1 | RW3 | 1.25 | 19 | 1 | 1 |
| scenario1 | RW3 | 1.25 | 20 | 1 | 1 |
| scenario1 | RW3 | 1.5 | 3 | 0.991929 | 0.993994 |
| scenario1 | RW3 | 1.5 | 4 | 0.998043 | 0.989 |
| scenario1 | RW3 | 1.5 | 5 | 0.999737 | 0.998 |
| scenario1 | RW3 | 1.5 | 6 | 0.99997 | 1 |
| scenario1 | RW3 | 1.5 | 7 | 0.999997 | 1 |
| scenario1 | RW3 | 1.5 | 8 | 0.999999 | 1 |
| scenario1 | RW3 | 1.5 | 9 | 1 | 1 |
| scenario1 | RW3 | 1.5 | 10 | 1 | 1 |
| scenario1 | RW3 | 1.5 | 11 | 1 | 1 |
| scenario1 | RW3 | 1.5 | 12 | 1 | 1 |
| scenario1 | RW3 | 1.5 | 13 | 1 | 1 |
| scenario1 | RW3 | 1.5 | 14 | 1 | 1 |
| scenario1 | RW3 | 1.5 | 15 | 1 | 1 |
| scenario1 | RW3 | 1.5 | 16 | 1 | 1 |
| scenario1 | RW3 | 1.5 | 17 | 1 | 1 |
| scenario1 | RW3 | 1.5 | 18 | 1 | 1 |
| scenario1 | RW3 | 1.5 | 19 | 1 | 1 |
| scenario1 | RW3 | 1.5 | 20 | 1 | 1 |
| scenario1 | RW3 | 1.75 | 3 | 0.99275 | 0.992 |
| scenario1 | RW3 | 1.75 | 4 | 0.999519 | 0.975 |
| scenario1 | RW3 | 1.75 | 5 | 0.99997 | 0.998 |
| scenario1 | RW3 | 1.75 | 6 | 0.999998 | 1 |
| scenario1 | RW3 | 1.75 | 7 | 1 | 1 |

| | | | | | |
|---|---|---|---|---|---|
| scenario1 | RW3 | 1.75 | 8 | 1 | 1 |
| scenario1 | RW3 | 1.75 | 9 | 1 | 1 |
| scenario1 | RW3 | 1.75 | 10 | 1 | 1 |
| scenario1 | RW3 | 1.75 | 11 | 1 | 1 |
| scenario1 | RW3 | 1.75 | 12 | 1 | 1 |
| scenario1 | RW3 | 1.75 | 13 | 1 | 1 |
| scenario1 | RW3 | 1.75 | 14 | 1 | 1 |
| scenario1 | RW3 | 1.75 | 15 | 1 | 1 |
| scenario1 | RW3 | 1.75 | 16 | 1 | 1 |
| scenario1 | RW3 | 1.75 | 17 | 1 | 1 |
| scenario1 | RW3 | 1.75 | 18 | 1 | 1 |
| scenario1 | RW3 | 1.75 | 19 | 1 | 1 |
| scenario1 | RW3 | 1.75 | 20 | 1 | 1 |
| scenario1 | RW4 | 1.25 | 3 | 0.071064 | 0.14786 |
| scenario1 | RW4 | 1.25 | 4 | 0.118379 | 0.037398 |
| scenario1 | RW4 | 1.25 | 5 | 0.166774 | 0.048295 |
| scenario1 | RW4 | 1.25 | 6 | 0.212616 | 0.071338 |
| scenario1 | RW4 | 1.25 | 7 | 0.254523 | 0.094775 |
| scenario1 | RW4 | 1.25 | 8 | 0.111717 | 0.121462 |
| scenario1 | RW4 | 1.25 | 9 | 0.143629 | 0.143337 |
| scenario1 | RW4 | 1.25 | 10 | 0.176666 | 0.165584 |
| scenario1 | RW4 | 1.25 | 11 | 0.183151 | 0.203209 |
| scenario1 | RW4 | 1.25 | 12 | 0.208687 | 0.234801 |
| scenario1 | RW4 | 1.25 | 13 | 0.233253 | 0.232461 |
| scenario1 | RW4 | 1.25 | 14 | 0.256821 | 0.258567 |
| scenario1 | RW4 | 1.25 | 15 | 0.27923 | 0.26762 |
| scenario1 | RW4 | 1.25 | 16 | 0.300858 | 0.29705 |
| scenario1 | RW4 | 1.25 | 17 | 0.321685 | 0.328267 |
| scenario1 | RW4 | 1.25 | 18 | 0.341808 | 0.332323 |
| scenario1 | RW4 | 1.25 | 19 | 0.361314 | 0.371717 |
| scenario1 | RW4 | 1.25 | 20 | 0.365812 | 0.35146 |
| scenario1 | RW4 | 1.5 | 3 | 0.801668 | 0.799389 |
| scenario1 | RW4 | 1.5 | 4 | 0.845266 | 0.705823 |
| scenario1 | RW4 | 1.5 | 5 | 0.907261 | 0.853 |
| scenario1 | RW4 | 1.5 | 6 | 0.949953 | 0.932933 |
| scenario1 | RW4 | 1.5 | 7 | 0.974812 | 0.961 |
| scenario1 | RW4 | 1.5 | 8 | 0.974927 | 0.959 |
| scenario1 | RW4 | 1.5 | 9 | 0.985939 | 0.978 |
| scenario1 | RW4 | 1.5 | 10 | 0.992396 | 0.987 |
| scenario1 | RW4 | 1.5 | 11 | 0.994676 | 0.996 |
| scenario1 | RW4 | 1.5 | 12 | 0.997521 | 0.997 |
| scenario1 | RW4 | 1.5 | 13 | 0.998863 | 1 |
| scenario1 | RW4 | 1.5 | 14 | 0.999216 | 1 |
| scenario1 | RW4 | 1.5 | 15 | 0.999321 | 1 |
| scenario1 | RW4 | 1.5 | 16 | 0.999652 | 1 |
| scenario1 | RW4 | 1.5 | 17 | 0.999826 | 0.999 |
| scenario1 | RW4 | 1.5 | 18 | 0.999915 | 1 |
| scenario1 | RW4 | 1.5 | 19 | 0.999959 | 1 |
| scenario1 | RW4 | 1.5 | 20 | 0.999977 | 1 |
| scenario1 | RW4 | 1.75 | 3 | 0.989363 | 0.987 |

| | | | | | |
|---|---|---|---|---|---|
| scenario1 | RW4 | 1.75 | 4 | 0.993667 | 0.972 |
| scenario1 | RW4 | 1.75 | 5 | 0.998954 | 0.995 |
| scenario1 | RW4 | 1.75 | 6 | 0.999862 | 1 |
| scenario1 | RW4 | 1.75 | 7 | 0.999984 | 1 |
| scenario1 | RW4 | 1.75 | 8 | 0.999987 | 0.999 |
| scenario1 | RW4 | 1.75 | 9 | 0.999998 | 1 |
| scenario1 | RW4 | 1.75 | 10 | 1 | 1 |
| scenario1 | RW4 | 1.75 | 11 | 1 | 1 |
| scenario1 | RW4 | 1.75 | 12 | 1 | 1 |
| scenario1 | RW4 | 1.75 | 13 | 1 | 1 |
| scenario1 | RW4 | 1.75 | 14 | 1 | 1 |
| scenario1 | RW4 | 1.75 | 15 | 1 | 1 |
| scenario1 | RW4 | 1.75 | 16 | 1 | 1 |
| scenario1 | RW4 | 1.75 | 17 | 1 | 1 |
| scenario1 | RW4 | 1.75 | 18 | 1 | 1 |
| scenario1 | RW4 | 1.75 | 19 | 1 | 1 |
| scenario1 | RW4 | 1.75 | 20 | 1 | 1 |
| scenario2 | RW1 | 2.5 | 3 | 0.920642 | 0.921844 |
| scenario2 | RW1 | 2.5 | 4 | 0.9241 | 0.886 |
| scenario2 | RW1 | 2.5 | 5 | 0.968202 | 0.965 |
| scenario2 | RW1 | 2.5 | 6 | 0.989139 | 0.991 |
| scenario2 | RW1 | 2.5 | 7 | 0.996597 | 0.986 |
| scenario2 | RW1 | 2.5 | 8 | 0.995662 | 0.998 |
| scenario2 | RW1 | 2.5 | 9 | 0.998379 | 1 |
| scenario2 | RW1 | 2.5 | 10 | 0.999448 | 1 |
| scenario2 | RW1 | 2.5 | 11 | 0.999806 | 1 |
| scenario2 | RW1 | 2.5 | 12 | 0.999942 | 1 |
| scenario2 | RW1 | 2.5 | 13 | 0.999983 | 1 |
| scenario2 | RW1 | 2.5 | 14 | 0.999979 | 1 |
| scenario2 | RW1 | 2.5 | 15 | 0.99999 | 1 |
| scenario2 | RW1 | 2.5 | 16 | 0.999997 | 1 |
| scenario2 | RW1 | 2.5 | 17 | 0.999999 | 1 |
| scenario2 | RW1 | 2.5 | 18 | 1 | 1 |
| scenario2 | RW1 | 2.5 | 19 | 1 | 1 |
| scenario2 | RW1 | 2.5 | 20 | 1 | 1 |
| scenario2 | RW1 | 2 | 3 | 0.279681 | 0.357786 |
| scenario2 | RW1 | 2 | 4 | 0.397071 | 0.14726 |
| scenario2 | RW1 | 2 | 5 | 0.49442 | 0.241158 |
| scenario2 | RW1 | 2 | 6 | 0.570041 | 0.340249 |
| scenario2 | RW1 | 2 | 7 | 0.630183 | 0.447179 |
| scenario2 | RW1 | 2 | 8 | 0.51955 | 0.467546 |
| scenario2 | RW1 | 2 | 9 | 0.595233 | 0.541877 |
| scenario2 | RW1 | 2 | 10 | 0.658743 | 0.592965 |
| scenario2 | RW1 | 2 | 11 | 0.631306 | 0.643932 |
| scenario2 | RW1 | 2 | 12 | 0.677014 | 0.704 |
| scenario2 | RW1 | 2 | 13 | 0.718389 | 0.745746 |
| scenario2 | RW1 | 2 | 14 | 0.755503 | 0.755511 |
| scenario2 | RW1 | 2 | 15 | 0.784085 | 0.792 |
| scenario2 | RW1 | 2 | 16 | 0.812371 | 0.797 |
| scenario2 | RW1 | 2 | 17 | 0.837163 | 0.817 |

| | | | | | |
|---|---|---|---|---|---|
| scenario2 | RW1 | 2 | 18 | 0.858773 | 0.825 |
| scenario2 | RW1 | 2 | 19 | 0.877539 | 0.854 |
| scenario2 | RW1 | 2 | 20 | 0.865756 | 0.873 |
| scenario2 | RW1 | 3 | 3 | 0.991512 | 0.989 |
| scenario2 | RW1 | 3 | 4 | 0.997308 | 0.992 |
| scenario2 | RW1 | 3 | 5 | 0.999615 | 0.999 |
| scenario2 | RW1 | 3 | 6 | 0.999954 | 1 |
| scenario2 | RW1 | 3 | 7 | 0.999995 | 1 |
| scenario2 | RW1 | 3 | 8 | 0.999998 | 1 |
| scenario2 | RW1 | 3 | 9 | 1 | 1 |
| scenario2 | RW1 | 3 | 10 | 1 | 1 |
| scenario2 | RW1 | 3 | 11 | 1 | 1 |
| scenario2 | RW1 | 3 | 12 | 1 | 1 |
| scenario2 | RW1 | 3 | 13 | 1 | 1 |
| scenario2 | RW1 | 3 | 14 | 1 | 1 |
| scenario2 | RW1 | 3 | 15 | 1 | 1 |
| scenario2 | RW1 | 3 | 16 | 1 | 1 |
| scenario2 | RW1 | 3 | 17 | 1 | 1 |
| scenario2 | RW1 | 3 | 18 | 1 | 1 |
| scenario2 | RW1 | 3 | 19 | 1 | 1 |
| scenario2 | RW1 | 3 | 20 | 1 | 1 |
| scenario2 | RW2 | 2.5 | 3 | 0.990665 | 0.988 |
| scenario2 | RW2 | 2.5 | 4 | 0.995846 | 0.982 |
| scenario2 | RW2 | 2.5 | 5 | 0.999361 | 0.996 |
| scenario2 | RW2 | 2.5 | 6 | 0.99992 | 1 |
| scenario2 | RW2 | 2.5 | 7 | 0.999991 | 1 |
| scenario2 | RW2 | 2.5 | 8 | 0.999995 | 1 |
| scenario2 | RW2 | 2.5 | 9 | 0.999999 | 1 |
| scenario2 | RW2 | 2.5 | 10 | 1 | 1 |
| scenario2 | RW2 | 2.5 | 11 | 1 | 1 |
| scenario2 | RW2 | 2.5 | 12 | 1 | 1 |
| scenario2 | RW2 | 2.5 | 13 | 1 | 1 |
| scenario2 | RW2 | 2.5 | 14 | 1 | 1 |
| scenario2 | RW2 | 2.5 | 15 | 1 | 1 |
| scenario2 | RW2 | 2.5 | 16 | 1 | 1 |
| scenario2 | RW2 | 2.5 | 17 | 1 | 1 |
| scenario2 | RW2 | 2.5 | 18 | 1 | 1 |
| scenario2 | RW2 | 2.5 | 19 | 1 | 1 |
| scenario2 | RW2 | 2.5 | 20 | 1 | 1 |
| scenario2 | RW2 | 2 | 3 | 0.628677 | 0.664916 |
| scenario2 | RW2 | 2 | 4 | 0.726469 | 0.464032 |
| scenario2 | RW2 | 2 | 5 | 0.806297 | 0.676382 |
| scenario2 | RW2 | 2 | 6 | 0.865045 | 0.795 |
| scenario2 | RW2 | 2 | 7 | 0.908682 | 0.878 |
| scenario2 | RW2 | 2 | 8 | 0.905921 | 0.856 |
| scenario2 | RW2 | 2 | 9 | 0.936738 | 0.908 |
| scenario2 | RW2 | 2 | 10 | 0.956997 | 0.938 |
| scenario2 | RW2 | 2 | 11 | 0.954939 | 0.96 |
| scenario2 | RW2 | 2 | 12 | 0.971271 | 0.971 |
| scenario2 | RW2 | 2 | 13 | 0.981993 | 0.979 |

| | | | | | |
|---|---|---|---|---|---|
| scenario2 | RW2 | 2 | 14 | 0.988064 | 0.976 |
| scenario2 | RW2 | 2 | 15 | 0.989662 | 0.988 |
| scenario2 | RW2 | 2 | 16 | 0.992997 | 0.988 |
| scenario2 | RW2 | 2 | 17 | 0.995289 | 0.997 |
| scenario2 | RW2 | 2 | 18 | 0.996861 | 0.994 |
| scenario2 | RW2 | 2 | 19 | 0.997932 | 0.997 |
| scenario2 | RW2 | 2 | 20 | 0.998258 | 0.998 |
| scenario2 | RW2 | 3 | 3 | 0.99275 | 0.992 |
| scenario2 | RW2 | 3 | 4 | 0.999519 | 0.986 |
| scenario2 | RW2 | 3 | 5 | 0.99997 | 1 |
| scenario2 | RW2 | 3 | 6 | 0.999998 | 1 |
| scenario2 | RW2 | 3 | 7 | 1 | 1 |
| scenario2 | RW2 | 3 | 8 | 1 | 1 |
| scenario2 | RW2 | 3 | 9 | 1 | 1 |
| scenario2 | RW2 | 3 | 10 | 1 | 1 |
| scenario2 | RW2 | 3 | 11 | 1 | 1 |
| scenario2 | RW2 | 3 | 12 | 1 | 1 |
| scenario2 | RW2 | 3 | 13 | 1 | 1 |
| scenario2 | RW2 | 3 | 14 | 1 | 1 |
| scenario2 | RW2 | 3 | 15 | 1 | 1 |
| scenario2 | RW2 | 3 | 16 | 1 | 1 |
| scenario2 | RW2 | 3 | 17 | 1 | 1 |
| scenario2 | RW2 | 3 | 18 | 1 | 1 |
| scenario2 | RW2 | 3 | 19 | 1 | 1 |
| scenario2 | RW2 | 3 | 20 | 1 | 1 |
| scenario2 | RW3 | 2.5 | 3 | 0.991929 | 0.993994 |
| scenario2 | RW3 | 2.5 | 4 | 0.998043 | 0.989 |
| scenario2 | RW3 | 2.5 | 5 | 0.999737 | 0.998 |
| scenario2 | RW3 | 2.5 | 6 | 0.99997 | 1 |
| scenario2 | RW3 | 2.5 | 7 | 0.999997 | 1 |
| scenario2 | RW3 | 2.5 | 8 | 0.999999 | 1 |
| scenario2 | RW3 | 2.5 | 9 | 1 | 1 |
| scenario2 | RW3 | 2.5 | 10 | 1 | 1 |
| scenario2 | RW3 | 2.5 | 11 | 1 | 1 |
| scenario2 | RW3 | 2.5 | 12 | 1 | 1 |
| scenario2 | RW3 | 2.5 | 13 | 1 | 1 |
| scenario2 | RW3 | 2.5 | 14 | 1 | 1 |
| scenario2 | RW3 | 2.5 | 15 | 1 | 1 |
| scenario2 | RW3 | 2.5 | 16 | 1 | 1 |
| scenario2 | RW3 | 2.5 | 17 | 1 | 1 |
| scenario2 | RW3 | 2.5 | 18 | 1 | 1 |
| scenario2 | RW3 | 2.5 | 19 | 1 | 1 |
| scenario2 | RW3 | 2.5 | 20 | 1 | 1 |
| scenario2 | RW3 | 2 | 3 | 0.836436 | 0.845921 |
| scenario2 | RW3 | 2 | 4 | 0.86744 | 0.753507 |
| scenario2 | RW3 | 2 | 5 | 0.925541 | 0.904 |
| scenario2 | RW3 | 2 | 6 | 0.963304 | 0.953 |
| scenario2 | RW3 | 2 | 7 | 0.983251 | 0.971 |
| scenario2 | RW3 | 2 | 8 | 0.982728 | 0.97 |
| scenario2 | RW3 | 2 | 9 | 0.990984 | 0.987 |

| | | | | | |
|---|---|---|---|---|---|
| scenario2 | RW3 | 2 | 10 | 0.995535 | 0.996 |
| scenario2 | RW3 | 2 | 11 | 0.997303 | 0.997 |
| scenario2 | RW3 | 2 | 12 | 0.998857 | 0.999 |
| scenario2 | RW3 | 2 | 13 | 0.999523 | 1 |
| scenario2 | RW3 | 2 | 14 | 0.999639 | 1 |
| scenario2 | RW3 | 2 | 15 | 0.99971 | 1 |
| scenario2 | RW3 | 2 | 16 | 0.999865 | 1 |
| scenario2 | RW3 | 2 | 17 | 0.999939 | 1 |
| scenario2 | RW3 | 2 | 18 | 0.999973 | 1 |
| scenario2 | RW3 | 2 | 19 | 0.999988 | 1 |
| scenario2 | RW3 | 2 | 20 | 0.999994 | 1 |
| scenario2 | RW3 | 3 | 3 | 0.99275 | 0.99 |
| scenario2 | RW3 | 3 | 4 | 0.999519 | 0.977 |
| scenario2 | RW3 | 3 | 5 | 0.99997 | 0.999 |
| scenario2 | RW3 | 3 | 6 | 0.999998 | 1 |
| scenario2 | RW3 | 3 | 7 | 1 | 1 |
| scenario2 | RW3 | 3 | 8 | 1 | 1 |
| scenario2 | RW3 | 3 | 9 | 1 | 1 |
| scenario2 | RW3 | 3 | 10 | 1 | 1 |
| scenario2 | RW3 | 3 | 11 | 1 | 1 |
| scenario2 | RW3 | 3 | 12 | 1 | 1 |
| scenario2 | RW3 | 3 | 13 | 1 | 1 |
| scenario2 | RW3 | 3 | 14 | 1 | 1 |
| scenario2 | RW3 | 3 | 15 | 1 | 1 |
| scenario2 | RW3 | 3 | 16 | 1 | 1 |
| scenario2 | RW3 | 3 | 17 | 1 | 1 |
| scenario2 | RW3 | 3 | 18 | 1 | 1 |
| scenario2 | RW3 | 3 | 19 | 1 | 1 |
| scenario2 | RW3 | 3 | 20 | 1 | 1 |
| scenario2 | RW4 | 2.5 | 3 | 0.801668 | 0.799389 |
| scenario2 | RW4 | 2.5 | 4 | 0.845266 | 0.705823 |
| scenario2 | RW4 | 2.5 | 5 | 0.907261 | 0.853 |
| scenario2 | RW4 | 2.5 | 6 | 0.949953 | 0.932933 |
| scenario2 | RW4 | 2.5 | 7 | 0.974812 | 0.961 |
| scenario2 | RW4 | 2.5 | 8 | 0.974927 | 0.959 |
| scenario2 | RW4 | 2.5 | 9 | 0.985939 | 0.978 |
| scenario2 | RW4 | 2.5 | 10 | 0.992396 | 0.987 |
| scenario2 | RW4 | 2.5 | 11 | 0.994676 | 0.996 |
| scenario2 | RW4 | 2.5 | 12 | 0.997521 | 0.997 |
| scenario2 | RW4 | 2.5 | 13 | 0.998863 | 1 |
| scenario2 | RW4 | 2.5 | 14 | 0.999216 | 1 |
| scenario2 | RW4 | 2.5 | 15 | 0.999321 | 1 |
| scenario2 | RW4 | 2.5 | 16 | 0.999652 | 1 |
| scenario2 | RW4 | 2.5 | 17 | 0.999826 | 0.999 |
| scenario2 | RW4 | 2.5 | 18 | 0.999915 | 1 |
| scenario2 | RW4 | 2.5 | 19 | 0.999959 | 1 |
| scenario2 | RW4 | 2.5 | 20 | 0.999977 | 1 |
| scenario2 | RW4 | 2 | 3 | 0.11499 | 0.148398 |
| scenario2 | RW4 | 2 | 4 | 0.18392 | 0.056787 |
| scenario2 | RW4 | 2 | 5 | 0.250541 | 0.080605 |

| | | | | | |
|---|---|---|---|---|---|
| scenario2 | RW4 | 2 | 6 | 0.310125 | 0.127315 |
| scenario2 | RW4 | 2 | 7 | 0.362003 | 0.165138 |
| scenario2 | RW4 | 2 | 8 | 0.201591 | 0.20282 |
| scenario2 | RW4 | 2 | 9 | 0.251284 | 0.249734 |
| scenario2 | RW4 | 2 | 10 | 0.300196 | 0.283489 |
| scenario2 | RW4 | 2 | 11 | 0.299243 | 0.300613 |
| scenario2 | RW4 | 2 | 12 | 0.334087 | 0.348932 |
| scenario2 | RW4 | 2 | 13 | 0.367028 | 0.356851 |
| scenario2 | RW4 | 2 | 14 | 0.398286 | 0.40404 |
| scenario2 | RW4 | 2 | 15 | 0.427329 | 0.425276 |
| scenario2 | RW4 | 2 | 16 | 0.455486 | 0.475806 |
| scenario2 | RW4 | 2 | 17 | 0.482438 | 0.474372 |
| scenario2 | RW4 | 2 | 18 | 0.508282 | 0.481444 |
| scenario2 | RW4 | 2 | 19 | 0.533077 | 0.513026 |
| scenario2 | RW4 | 2 | 20 | 0.529414 | 0.515516 |
| scenario2 | RW4 | 3 | 3 | 0.986644 | 0.984 |
| scenario2 | RW4 | 3 | 4 | 0.989365 | 0.977 |
| scenario2 | RW4 | 3 | 5 | 0.998046 | 0.997 |
| scenario2 | RW4 | 3 | 6 | 0.999719 | 1 |
| scenario2 | RW4 | 3 | 7 | 0.999964 | 1 |
| scenario2 | RW4 | 3 | 8 | 0.999961 | 1 |
| scenario2 | RW4 | 3 | 9 | 0.999994 | 1 |
| scenario2 | RW4 | 3 | 10 | 0.999999 | 1 |
| scenario2 | RW4 | 3 | 11 | 1 | 1 |
| scenario2 | RW4 | 3 | 12 | 1 | 1 |
| scenario2 | RW4 | 3 | 13 | 1 | 1 |
| scenario2 | RW4 | 3 | 14 | 1 | 1 |
| scenario2 | RW4 | 3 | 15 | 1 | 1 |
| scenario2 | RW4 | 3 | 16 | 1 | 1 |
| scenario2 | RW4 | 3 | 17 | 1 | 1 |
| scenario2 | RW4 | 3 | 18 | 1 | 1 |
| scenario2 | RW4 | 3 | 19 | 1 | 1 |
| scenario2 | RW4 | 3 | 20 | 1 | 1 |
| scenario3 | RW1 | 0.2 | 3 | 0.964752 | 0.973974 |
| scenario3 | RW1 | 0.2 | 4 | 0.962186 | 0.951 |
| scenario3 | RW1 | 0.2 | 5 | 0.989148 | 0.988 |
| scenario3 | RW1 | 0.2 | 6 | 0.997564 | 0.999 |
| scenario3 | RW1 | 0.2 | 7 | 0.999499 | 1 |
| scenario3 | RW1 | 0.2 | 8 | 0.999239 | 1 |
| scenario3 | RW1 | 0.2 | 9 | 0.999813 | 1 |
| scenario3 | RW1 | 0.2 | 10 | 0.999959 | 1 |
| scenario3 | RW1 | 0.2 | 11 | 0.999991 | 1 |
| scenario3 | RW1 | 0.2 | 12 | 0.999998 | 1 |
| scenario3 | RW1 | 0.2 | 13 | 1 | 1 |
| scenario3 | RW1 | 0.2 | 14 | 0.999999 | 1 |
| scenario3 | RW1 | 0.2 | 15 | 1 | 1 |
| scenario3 | RW1 | 0.2 | 16 | 1 | 1 |
| scenario3 | RW1 | 0.2 | 17 | 1 | 1 |
| scenario3 | RW1 | 0.2 | 18 | 1 | 1 |
| scenario3 | RW1 | 0.2 | 19 | 1 | 1 |

| | | | | | |
|---|---|---|---|---|---|
| scenario3 | RW1 | 0.2 | 20 | 1 | 1 |
| scenario3 | RW1 | 0.5 | 3 | 0.920642 | 0.921844 |
| scenario3 | RW1 | 0.5 | 4 | 0.9241 | 0.886 |
| scenario3 | RW1 | 0.5 | 5 | 0.968202 | 0.965 |
| scenario3 | RW1 | 0.5 | 6 | 0.989139 | 0.991 |
| scenario3 | RW1 | 0.5 | 7 | 0.996597 | 0.986 |
| scenario3 | RW1 | 0.5 | 8 | 0.995662 | 0.998 |
| scenario3 | RW1 | 0.5 | 9 | 0.998379 | 1 |
| scenario3 | RW1 | 0.5 | 10 | 0.999448 | 1 |
| scenario3 | RW1 | 0.5 | 11 | 0.999806 | 1 |
| scenario3 | RW1 | 0.5 | 12 | 0.999942 | 1 |
| scenario3 | RW1 | 0.5 | 13 | 0.999983 | 1 |
| scenario3 | RW1 | 0.5 | 14 | 0.999979 | 1 |
| scenario3 | RW1 | 0.5 | 15 | 0.99999 | 1 |
| scenario3 | RW1 | 0.5 | 16 | 0.999997 | 1 |
| scenario3 | RW1 | 0.5 | 17 | 0.999999 | 1 |
| scenario3 | RW1 | 0.5 | 18 | 1 | 1 |
| scenario3 | RW1 | 0.5 | 19 | 1 | 1 |
| scenario3 | RW1 | 0.5 | 20 | 1 | 1 |
| scenario3 | RW1 | 0.8 | 3 | 0.757511 | 0.762781 |
| scenario3 | RW1 | 0.8 | 4 | 0.816712 | 0.627255 |
| scenario3 | RW1 | 0.8 | 5 | 0.883299 | 0.792793 |
| scenario3 | RW1 | 0.8 | 6 | 0.931132 | 0.906 |
| scenario3 | RW1 | 0.8 | 7 | 0.961716 | 0.94 |
| scenario3 | RW1 | 0.8 | 8 | 0.962528 | 0.936 |
| scenario3 | RW1 | 0.8 | 9 | 0.977567 | 0.962 |
| scenario3 | RW1 | 0.8 | 10 | 0.986814 | 0.98 |
| scenario3 | RW1 | 0.8 | 11 | 0.989298 | 0.991 |
| scenario3 | RW1 | 0.8 | 12 | 0.994493 | 0.995 |
| scenario3 | RW1 | 0.8 | 13 | 0.997212 | 0.999 |
| scenario3 | RW1 | 0.8 | 14 | 0.998185 | 0.997 |
| scenario3 | RW1 | 0.8 | 15 | 0.998371 | 0.999 |
| scenario3 | RW1 | 0.8 | 16 | 0.99908 | 1 |
| scenario3 | RW1 | 0.8 | 17 | 0.999491 | 1 |
| scenario3 | RW1 | 0.8 | 18 | 0.999724 | 1 |
| scenario3 | RW1 | 0.8 | 19 | 0.999853 | 1 |
| scenario3 | RW1 | 0.8 | 20 | 0.999909 | 1 |
| scenario3 | RW2 | 0.2 | 3 | 0.99275 | 0.993 |
| scenario3 | RW2 | 0.2 | 4 | 0.999519 | 0.985 |
| scenario3 | RW2 | 0.2 | 5 | 0.99997 | 1 |
| scenario3 | RW2 | 0.2 | 6 | 0.999998 | 1 |
| scenario3 | RW2 | 0.2 | 7 | 1 | 1 |
| scenario3 | RW2 | 0.2 | 8 | 1 | 1 |
| scenario3 | RW2 | 0.2 | 9 | 1 | 1 |
| scenario3 | RW2 | 0.2 | 10 | 1 | 1 |
| scenario3 | RW2 | 0.2 | 11 | 1 | 1 |
| scenario3 | RW2 | 0.2 | 12 | 1 | 1 |
| scenario3 | RW2 | 0.2 | 13 | 1 | 1 |
| scenario3 | RW2 | 0.2 | 14 | 1 | 1 |
| scenario3 | RW2 | 0.2 | 15 | 1 | 1 |

| | | | | | |
|---|---|---|---|---|---|
| scenario3 | RW2 | 0.2 | 16 | 1 | 1 |
| scenario3 | RW2 | 0.2 | 17 | 1 | 1 |
| scenario3 | RW2 | 0.2 | 18 | 1 | 1 |
| scenario3 | RW2 | 0.2 | 19 | 1 | 1 |
| scenario3 | RW2 | 0.2 | 20 | 1 | 1 |
| scenario3 | RW2 | 0.5 | 3 | 0.990665 | 0.988 |
| scenario3 | RW2 | 0.5 | 4 | 0.995846 | 0.982 |
| scenario3 | RW2 | 0.5 | 5 | 0.999361 | 0.996 |
| scenario3 | RW2 | 0.5 | 6 | 0.99992 | 1 |
| scenario3 | RW2 | 0.5 | 7 | 0.999991 | 1 |
| scenario3 | RW2 | 0.5 | 8 | 0.999995 | 1 |
| scenario3 | RW2 | 0.5 | 9 | 0.999999 | 1 |
| scenario3 | RW2 | 0.5 | 10 | 1 | 1 |
| scenario3 | RW2 | 0.5 | 11 | 1 | 1 |
| scenario3 | RW2 | 0.5 | 12 | 1 | 1 |
| scenario3 | RW2 | 0.5 | 13 | 1 | 1 |
| scenario3 | RW2 | 0.5 | 14 | 1 | 1 |
| scenario3 | RW2 | 0.5 | 15 | 1 | 1 |
| scenario3 | RW2 | 0.5 | 16 | 1 | 1 |
| scenario3 | RW2 | 0.5 | 17 | 1 | 1 |
| scenario3 | RW2 | 0.5 | 18 | 1 | 1 |
| scenario3 | RW2 | 0.5 | 19 | 1 | 1 |
| scenario3 | RW2 | 0.5 | 20 | 1 | 1 |
| scenario3 | RW2 | 0.8 | 3 | 0.967835 | 0.969 |
| scenario3 | RW2 | 0.8 | 4 | 0.965466 | 0.954 |
| scenario3 | RW2 | 0.8 | 5 | 0.990509 | 0.988 |
| scenario3 | RW2 | 0.8 | 6 | 0.997965 | 0.997 |
| scenario3 | RW2 | 0.8 | 7 | 0.9996 | 0.997 |
| scenario3 | RW2 | 0.8 | 8 | 0.99939 | 0.998 |
| scenario3 | RW2 | 0.8 | 9 | 0.999857 | 1 |
| scenario3 | RW2 | 0.8 | 10 | 0.99997 | 1 |
| scenario3 | RW2 | 0.8 | 11 | 0.999994 | 1 |
| scenario3 | RW2 | 0.8 | 12 | 0.999999 | 1 |
| scenario3 | RW2 | 0.8 | 13 | 1 | 1 |
| scenario3 | RW2 | 0.8 | 14 | 1 | 1 |
| scenario3 | RW2 | 0.8 | 15 | 1 | 1 |
| scenario3 | RW2 | 0.8 | 16 | 1 | 1 |
| scenario3 | RW2 | 0.8 | 17 | 1 | 1 |
| scenario3 | RW2 | 0.8 | 18 | 1 | 1 |
| scenario3 | RW2 | 0.8 | 19 | 1 | 1 |
| scenario3 | RW2 | 0.8 | 20 | 1 | 1 |
| scenario3 | RW3 | 0.2 | 3 | 0.99275 | 0.99 |
| scenario3 | RW3 | 0.2 | 4 | 0.999519 | 0.986 |
| scenario3 | RW3 | 0.2 | 5 | 0.99997 | 1 |
| scenario3 | RW3 | 0.2 | 6 | 0.999998 | 1 |
| scenario3 | RW3 | 0.2 | 7 | 1 | 1 |
| scenario3 | RW3 | 0.2 | 8 | 1 | 1 |
| scenario3 | RW3 | 0.2 | 9 | 1 | 1 |
| scenario3 | RW3 | 0.2 | 10 | 1 | 1 |
| scenario3 | RW3 | 0.2 | 11 | 1 | 1 |

| | | | | | |
|---|---|---|---|---|---|
| scenario3 | RW3 | 0.2 | 12 | 1 | 1 |
| scenario3 | RW3 | 0.2 | 13 | 1 | 1 |
| scenario3 | RW3 | 0.2 | 14 | 1 | 1 |
| scenario3 | RW3 | 0.2 | 15 | 1 | 1 |
| scenario3 | RW3 | 0.2 | 16 | 1 | 1 |
| scenario3 | RW3 | 0.2 | 17 | 1 | 1 |
| scenario3 | RW3 | 0.2 | 18 | 1 | 1 |
| scenario3 | RW3 | 0.2 | 19 | 1 | 1 |
| scenario3 | RW3 | 0.2 | 20 | 1 | 1 |
| scenario3 | RW3 | 0.5 | 3 | 0.991929 | 0.993994 |
| scenario3 | RW3 | 0.5 | 4 | 0.998043 | 0.989 |
| scenario3 | RW3 | 0.5 | 5 | 0.999737 | 0.998 |
| scenario3 | RW3 | 0.5 | 6 | 0.99997 | 1 |
| scenario3 | RW3 | 0.5 | 7 | 0.999997 | 1 |
| scenario3 | RW3 | 0.5 | 8 | 0.999999 | 1 |
| scenario3 | RW3 | 0.5 | 9 | 1 | 1 |
| scenario3 | RW3 | 0.5 | 10 | 1 | 1 |
| scenario3 | RW3 | 0.5 | 11 | 1 | 1 |
| scenario3 | RW3 | 0.5 | 12 | 1 | 1 |
| scenario3 | RW3 | 0.5 | 13 | 1 | 1 |
| scenario3 | RW3 | 0.5 | 14 | 1 | 1 |
| scenario3 | RW3 | 0.5 | 15 | 1 | 1 |
| scenario3 | RW3 | 0.5 | 16 | 1 | 1 |
| scenario3 | RW3 | 0.5 | 17 | 1 | 1 |
| scenario3 | RW3 | 0.5 | 18 | 1 | 1 |
| scenario3 | RW3 | 0.5 | 19 | 1 | 1 |
| scenario3 | RW3 | 0.5 | 20 | 1 | 1 |
| scenario3 | RW3 | 0.8 | 3 | 0.989802 | 0.993 |
| scenario3 | RW3 | 0.8 | 4 | 0.994391 | 0.987 |
| scenario3 | RW3 | 0.8 | 5 | 0.999093 | 0.998 |
| scenario3 | RW3 | 0.8 | 6 | 0.999883 | 1 |
| scenario3 | RW3 | 0.8 | 7 | 0.999987 | 1 |
| scenario3 | RW3 | 0.8 | 8 | 0.99999 | 1 |
| scenario3 | RW3 | 0.8 | 9 | 0.999999 | 1 |
| scenario3 | RW3 | 0.8 | 10 | 1 | 1 |
| scenario3 | RW3 | 0.8 | 11 | 1 | 1 |
| scenario3 | RW3 | 0.8 | 12 | 1 | 1 |
| scenario3 | RW3 | 0.8 | 13 | 1 | 1 |
| scenario3 | RW3 | 0.8 | 14 | 1 | 1 |
| scenario3 | RW3 | 0.8 | 15 | 1 | 1 |
| scenario3 | RW3 | 0.8 | 16 | 1 | 1 |
| scenario3 | RW3 | 0.8 | 17 | 1 | 1 |
| scenario3 | RW3 | 0.8 | 18 | 1 | 1 |
| scenario3 | RW3 | 0.8 | 19 | 1 | 1 |
| scenario3 | RW3 | 0.8 | 20 | 1 | 1 |
| scenario3 | RW4 | 0.2 | 3 | 0.905927 | 0.891348 |
| scenario3 | RW4 | 0.2 | 4 | 0.913408 | 0.835 |
| scenario3 | RW4 | 0.2 | 5 | 0.960912 | 0.943 |
| scenario3 | RW4 | 0.2 | 6 | 0.985413 | 0.983 |
| scenario3 | RW4 | 0.2 | 7 | 0.995005 | 0.983 |

| | | | | | |
|---|---|---|---|---|---|
| scenario3 | RW4 | 0.2 | 8 | 0.993963 | 0.995 |
| scenario3 | RW4 | 0.2 | 9 | 0.997542 | 0.996 |
| scenario3 | RW4 | 0.2 | 10 | 0.999083 | 1 |
| scenario3 | RW4 | 0.2 | 11 | 0.999635 | 1 |
| scenario3 | RW4 | 0.2 | 12 | 0.999882 | 1 |
| scenario3 | RW4 | 0.2 | 13 | 0.999962 | 1 |
| scenario3 | RW4 | 0.2 | 14 | 0.999959 | 1 |
| scenario3 | RW4 | 0.2 | 15 | 0.999977 | 1 |
| scenario3 | RW4 | 0.2 | 16 | 0.999992 | 1 |
| scenario3 | RW4 | 0.2 | 17 | 0.999997 | 1 |
| scenario3 | RW4 | 0.2 | 18 | 0.999999 | 1 |
| scenario3 | RW4 | 0.2 | 19 | 1 | 1 |
| scenario3 | RW4 | 0.2 | 20 | 1 | 1 |
| scenario3 | RW4 | 0.5 | 3 | 0.801668 | 0.799389 |
| scenario3 | RW4 | 0.5 | 4 | 0.845266 | 0.705823 |
| scenario3 | RW4 | 0.5 | 5 | 0.907261 | 0.853 |
| scenario3 | RW4 | 0.5 | 6 | 0.949953 | 0.932933 |
| scenario3 | RW4 | 0.5 | 7 | 0.974812 | 0.961 |
| scenario3 | RW4 | 0.5 | 8 | 0.974927 | 0.959 |
| scenario3 | RW4 | 0.5 | 9 | 0.985939 | 0.978 |
| scenario3 | RW4 | 0.5 | 10 | 0.992396 | 0.987 |
| scenario3 | RW4 | 0.5 | 11 | 0.994676 | 0.996 |
| scenario3 | RW4 | 0.5 | 12 | 0.997521 | 0.997 |
| scenario3 | RW4 | 0.5 | 13 | 0.998863 | 1 |
| scenario3 | RW4 | 0.5 | 14 | 0.999216 | 1 |
| scenario3 | RW4 | 0.5 | 15 | 0.999321 | 1 |
| scenario3 | RW4 | 0.5 | 16 | 0.999652 | 1 |
| scenario3 | RW4 | 0.5 | 17 | 0.999826 | 0.999 |
| scenario3 | RW4 | 0.5 | 18 | 0.999915 | 1 |
| scenario3 | RW4 | 0.5 | 19 | 0.999959 | 1 |
| scenario3 | RW4 | 0.5 | 20 | 0.999977 | 1 |
| scenario3 | RW4 | 0.8 | 3 | 0.564052 | 0.603672 |
| scenario3 | RW4 | 0.8 | 4 | 0.675791 | 0.425926 |
| scenario3 | RW4 | 0.8 | 5 | 0.762105 | 0.584337 |
| scenario3 | RW4 | 0.8 | 6 | 0.825241 | 0.693079 |
| scenario3 | RW4 | 0.8 | 7 | 0.873621 | 0.814629 |
| scenario3 | RW4 | 0.8 | 8 | 0.86306 | 0.794 |
| scenario3 | RW4 | 0.8 | 9 | 0.903844 | 0.85 |
| scenario3 | RW4 | 0.8 | 10 | 0.931512 | 0.881 |
| scenario3 | RW4 | 0.8 | 11 | 0.923903 | 0.924 |
| scenario3 | RW4 | 0.8 | 12 | 0.947407 | 0.948 |
| scenario3 | RW4 | 0.8 | 13 | 0.964227 | 0.965 |
| scenario3 | RW4 | 0.8 | 14 | 0.975122 | 0.976 |
| scenario3 | RW4 | 0.8 | 15 | 0.978953 | 0.99 |
| scenario3 | RW4 | 0.8 | 16 | 0.984877 | 0.983 |
| scenario3 | RW4 | 0.8 | 17 | 0.989158 | 0.982 |
| scenario3 | RW4 | 0.8 | 18 | 0.992262 | 0.992 |
| scenario3 | RW4 | 0.8 | 19 | 0.994514 | 0.993 |
| scenario3 | RW4 | 0.8 | 20 | 0.994818 | 0.996 |

| Scenario | Spatial Pat | Parameter | Parameter | N_Rep | Estimated | Observed Power |
|---|---|---|---|---|---|---|
| scenario1 | RW1 | 1.25 | 1.5 | 3 | 0.178464 | 0.059 |
| scenario1 | RW1 | 1.25 | 1.5 | 4 | 0.254754 | 0.147 |
| scenario1 | RW1 | 1.25 | 1.5 | 5 | 0.328652 | 0.253 |
| scenario1 | RW1 | 1.25 | 1.5 | 6 | 0.399003 | 0.36 |
| scenario1 | RW1 | 1.25 | 1.5 | 7 | 0.46507 | 0.429 |
| scenario1 | RW1 | 1.25 | 1.5 | 8 | 0.526388 | 0.516 |
| scenario1 | RW1 | 1.25 | 1.5 | 9 | 0.582718 | 0.609 |
| scenario1 | RW1 | 1.25 | 1.5 | 10 | 0.634002 | 0.662 |
| scenario1 | RW1 | 1.25 | 1.5 | 11 | 0.68032 | 0.736 |
| scenario1 | RW1 | 1.25 | 1.5 | 12 | 0.721858 | 0.771 |
| scenario1 | RW1 | 1.25 | 1.5 | 13 | 0.758871 | 0.83 |
| scenario1 | RW1 | 1.25 | 1.5 | 14 | 0.791663 | 0.859 |
| scenario1 | RW1 | 1.25 | 1.5 | 15 | 0.820565 | 0.9 |
| scenario1 | RW1 | 1.25 | 1.5 | 16 | 0.845916 | 0.892 |
| scenario1 | RW1 | 1.25 | 1.5 | 17 | 0.868055 | 0.94 |
| scenario1 | RW1 | 1.25 | 1.5 | 18 | 0.887311 | 0.942 |
| scenario1 | RW1 | 1.25 | 1.5 | 19 | 0.903997 | 0.955 |
| scenario1 | RW1 | 1.25 | 1.5 | 20 | 0.918405 | 0.967 |
| scenario1 | RW1 | 1.25 | 1.75 | 3 | 0.677758 | 0.675 |
| scenario1 | RW1 | 1.25 | 1.75 | 4 | 0.873512 | 0.911 |
| scenario1 | RW1 | 1.25 | 1.75 | 5 | 0.954811 | 0.983 |
| scenario1 | RW1 | 1.25 | 1.75 | 6 | 0.984852 | 0.998 |
| scenario1 | RW1 | 1.25 | 1.75 | 7 | 0.995162 | 1 |
| scenario1 | RW1 | 1.25 | 1.75 | 8 | 0.998514 | 1 |
| scenario1 | RW1 | 1.25 | 1.75 | 9 | 0.999558 | 1 |
| scenario1 | RW1 | 1.25 | 1.75 | 10 | 0.999872 | 1 |
| scenario1 | RW1 | 1.25 | 1.75 | 11 | 0.999964 | 1 |
| scenario1 | RW1 | 1.25 | 1.75 | 12 | 0.99999 | 1 |
| scenario1 | RW1 | 1.25 | 1.75 | 13 | 0.999997 | 1 |
| scenario1 | RW1 | 1.25 | 1.75 | 14 | 0.999999 | 1 |
| scenario1 | RW1 | 1.25 | 1.75 | 15 | 1 | 1 |
| scenario1 | RW1 | 1.25 | 1.75 | 16 | 1 | 1 |
| scenario1 | RW1 | 1.25 | 1.75 | 17 | 1 | 1 |
| scenario1 | RW1 | 1.25 | 1.75 | 18 | 1 | 1 |
| scenario1 | RW1 | 1.25 | 1.75 | 19 | 1 | 1 |
| scenario1 | RW1 | 1.25 | 1.75 | 20 | 1 | 1 |
| scenario1 | RW1 | 1.5 | 1.75 | 3 | 0.319242 | 0.218 |
| scenario1 | RW1 | 1.5 | 1.75 | 4 | 0.468712 | 0.434 |
| scenario1 | RW1 | 1.5 | 1.75 | 5 | 0.595157 | 0.618 |
| scenario1 | RW1 | 1.5 | 1.75 | 6 | 0.697416 | 0.724 |
| scenario1 | RW1 | 1.5 | 1.75 | 7 | 0.777527 | 0.805 |
| scenario1 | RW1 | 1.5 | 1.75 | 8 | 0.838738 | 0.885 |
| scenario1 | RW1 | 1.5 | 1.75 | 9 | 0.884558 | 0.929 |
| scenario1 | RW1 | 1.5 | 1.75 | 10 | 0.918269 | 0.956 |
| scenario1 | RW1 | 1.5 | 1.75 | 11 | 0.942707 | 0.976 |
| scenario1 | RW1 | 1.5 | 1.75 | 12 | 0.960195 | 0.987 |
| scenario1 | RW1 | 1.5 | 1.75 | 13 | 0.972569 | 0.993 |
| scenario1 | RW1 | 1.5 | 1.75 | 14 | 0.981236 | 0.995 |
| scenario1 | RW1 | 1.5 | 1.75 | 15 | 0.987251 | 0.998 |

| | | | | | | |
|---|---|---|---|---|---|---|
| scenario1 | RW1 | 1.5 | 1.75 | 16 | 0.991392 | 0.996 |
| scenario1 | RW1 | 1.5 | 1.75 | 17 | 0.994222 | 1 |
| scenario1 | RW1 | 1.5 | 1.75 | 18 | 0.996142 | 0.999 |
| scenario1 | RW1 | 1.5 | 1.75 | 19 | 0.997437 | 1 |
| scenario1 | RW1 | 1.5 | 1.75 | 20 | 0.998306 | 1 |
| scenario1 | RW2 | 1.25 | 1.5 | 3 | 0.255672 | 0.146 |
| scenario1 | RW2 | 1.25 | 1.5 | 4 | 0.374885 | 0.292 |
| scenario1 | RW2 | 1.25 | 1.5 | 5 | 0.483108 | 0.453 |
| scenario1 | RW2 | 1.25 | 1.5 | 6 | 0.578161 | 0.574 |
| scenario1 | RW2 | 1.25 | 1.5 | 7 | 0.659655 | 0.687 |
| scenario1 | RW2 | 1.25 | 1.5 | 8 | 0.728177 | 0.781 |
| scenario1 | RW2 | 1.25 | 1.5 | 9 | 0.784862 | 0.822 |
| scenario1 | RW2 | 1.25 | 1.5 | 10 | 0.83111 | 0.885 |
| scenario1 | RW2 | 1.25 | 1.5 | 11 | 0.868392 | 0.904 |
| scenario1 | RW2 | 1.25 | 1.5 | 12 | 0.898133 | 0.94 |
| scenario1 | RW2 | 1.25 | 1.5 | 13 | 0.921638 | 0.955 |
| scenario1 | RW2 | 1.25 | 1.5 | 14 | 0.940061 | 0.966 |
| scenario1 | RW2 | 1.25 | 1.5 | 15 | 0.954392 | 0.982 |
| scenario1 | RW2 | 1.25 | 1.5 | 16 | 0.965465 | 0.988 |
| scenario1 | RW2 | 1.25 | 1.5 | 17 | 0.973968 | 0.995 |
| scenario1 | RW2 | 1.25 | 1.5 | 18 | 0.980461 | 0.994 |
| scenario1 | RW2 | 1.25 | 1.5 | 19 | 0.985392 | 0.995 |
| scenario1 | RW2 | 1.25 | 1.5 | 20 | 0.989119 | 0.997 |
| scenario1 | RW2 | 1.25 | 1.75 | 3 | 0.646852 | 0.581 |
| scenario1 | RW2 | 1.25 | 1.75 | 4 | 0.848258 | 0.841 |
| scenario1 | RW2 | 1.25 | 1.75 | 5 | 0.940131 | 0.953 |
| scenario1 | RW2 | 1.25 | 1.75 | 6 | 0.977726 | 0.988 |
| scenario1 | RW2 | 1.25 | 1.75 | 7 | 0.992078 | 0.997 |
| scenario1 | RW2 | 1.25 | 1.75 | 8 | 0.997283 | 0.997 |
| scenario1 | RW2 | 1.25 | 1.75 | 9 | 0.999096 | 1 |
| scenario1 | RW2 | 1.25 | 1.75 | 10 | 0.999707 | 1 |
| scenario1 | RW2 | 1.25 | 1.75 | 11 | 0.999907 | 1 |
| scenario1 | RW2 | 1.25 | 1.75 | 12 | 0.999971 | 1 |
| scenario1 | RW2 | 1.25 | 1.75 | 13 | 0.999991 | 1 |
| scenario1 | RW2 | 1.25 | 1.75 | 14 | 0.999997 | 1 |
| scenario1 | RW2 | 1.25 | 1.75 | 15 | 0.999999 | 1 |
| scenario1 | RW2 | 1.25 | 1.75 | 16 | 1 | 1 |
| scenario1 | RW2 | 1.25 | 1.75 | 17 | 1 | 1 |
| scenario1 | RW2 | 1.25 | 1.75 | 18 | 1 | 1 |
| scenario1 | RW2 | 1.25 | 1.75 | 19 | 1 | 1 |
| scenario1 | RW2 | 1.25 | 1.75 | 20 | 1 | 1 |
| scenario1 | RW2 | 1.5 | 1.75 | 3 | 0.202032 | 0.108 |
| scenario1 | RW2 | 1.5 | 1.75 | 4 | 0.291757 | 0.204 |
| scenario1 | RW2 | 1.5 | 1.75 | 5 | 0.377198 | 0.315 |
| scenario1 | RW2 | 1.5 | 1.75 | 6 | 0.456792 | 0.403 |
| scenario1 | RW2 | 1.5 | 1.75 | 7 | 0.529701 | 0.499 |
| scenario1 | RW2 | 1.5 | 1.75 | 8 | 0.595546 | 0.591 |
| scenario1 | RW2 | 1.5 | 1.75 | 9 | 0.654293 | 0.66 |
| scenario1 | RW2 | 1.5 | 1.75 | 10 | 0.706157 | 0.729 |
| scenario1 | RW2 | 1.5 | 1.75 | 11 | 0.751523 | 0.747 |

| | | | | | | |
|---|---|---|---|---|---|---|
| scenario1 | RW2 | 1.5 | 1.75 | 12 | 0.79088 | 0.827 |
| scenario1 | RW2 | 1.5 | 1.75 | 13 | 0.824775 | 0.847 |
| scenario1 | RW2 | 1.5 | 1.75 | 14 | 0.853775 | 0.866 |
| scenario1 | RW2 | 1.5 | 1.75 | 15 | 0.87844 | 0.914 |
| scenario1 | RW2 | 1.5 | 1.75 | 16 | 0.899304 | 0.924 |
| scenario1 | RW2 | 1.5 | 1.75 | 17 | 0.916865 | 0.942 |
| scenario1 | RW2 | 1.5 | 1.75 | 18 | 0.931578 | 0.946 |
| scenario1 | RW2 | 1.5 | 1.75 | 19 | 0.943854 | 0.961 |
| scenario1 | RW2 | 1.5 | 1.75 | 20 | 0.954056 | 0.971 |
| scenario1 | RW3 | 1.25 | 1.5 | 3 | 0.377442 | 0.274 |
| scenario1 | RW3 | 1.25 | 1.5 | 4 | 0.549622 | 0.494 |
| scenario1 | RW3 | 1.25 | 1.5 | 5 | 0.684556 | 0.693 |
| scenario1 | RW3 | 1.25 | 1.5 | 6 | 0.784596 | 0.825 |
| scenario1 | RW3 | 1.25 | 1.5 | 7 | 0.855974 | 0.908 |
| scenario1 | RW3 | 1.25 | 1.5 | 8 | 0.905413 | 0.952 |
| scenario1 | RW3 | 1.25 | 1.5 | 9 | 0.938841 | 0.978 |
| scenario1 | RW3 | 1.25 | 1.5 | 10 | 0.960992 | 0.986 |
| scenario1 | RW3 | 1.25 | 1.5 | 11 | 0.97542 | 0.996 |
| scenario1 | RW3 | 1.25 | 1.5 | 12 | 0.984679 | 0.996 |
| scenario1 | RW3 | 1.25 | 1.5 | 13 | 0.990544 | 0.999 |
| scenario1 | RW3 | 1.25 | 1.5 | 14 | 0.994215 | 0.999 |
| scenario1 | RW3 | 1.25 | 1.5 | 15 | 0.99649 | 1 |
| scenario1 | RW3 | 1.25 | 1.5 | 16 | 0.997886 | 1 |
| scenario1 | RW3 | 1.25 | 1.5 | 17 | 0.998735 | 1 |
| scenario1 | RW3 | 1.25 | 1.5 | 18 | 0.999248 | 1 |
| scenario1 | RW3 | 1.25 | 1.5 | 19 | 0.999556 | 1 |
| scenario1 | RW3 | 1.25 | 1.5 | 20 | 0.999739 | 1 |
| scenario1 | RW3 | 1.25 | 1.75 | 3 | 0.882223 | 0.887 |
| scenario1 | RW3 | 1.25 | 1.75 | 4 | 0.982617 | 0.997 |
| scenario1 | RW3 | 1.25 | 1.75 | 5 | 0.997835 | 0.999 |
| scenario1 | RW3 | 1.25 | 1.75 | 6 | 0.999757 | 1 |
| scenario1 | RW3 | 1.25 | 1.75 | 7 | 0.999975 | 1 |
| scenario1 | RW3 | 1.25 | 1.75 | 8 | 0.999998 | 1 |
| scenario1 | RW3 | 1.25 | 1.75 | 9 | 1 | 1 |
| scenario1 | RW3 | 1.25 | 1.75 | 10 | 1 | 1 |
| scenario1 | RW3 | 1.25 | 1.75 | 11 | 1 | 1 |
| scenario1 | RW3 | 1.25 | 1.75 | 12 | 1 | 1 |
| scenario1 | RW3 | 1.25 | 1.75 | 13 | 1 | 1 |
| scenario1 | RW3 | 1.25 | 1.75 | 14 | 1 | 1 |
| scenario1 | RW3 | 1.25 | 1.75 | 15 | 1 | 1 |
| scenario1 | RW3 | 1.25 | 1.75 | 16 | 1 | 1 |
| scenario1 | RW3 | 1.25 | 1.75 | 17 | 1 | 1 |
| scenario1 | RW3 | 1.25 | 1.75 | 18 | 1 | 1 |
| scenario1 | RW3 | 1.25 | 1.75 | 19 | 1 | 1 |
| scenario1 | RW3 | 1.25 | 1.75 | 20 | 1 | 1 |
| scenario1 | RW3 | 1.5 | 1.75 | 3 | 0.395435 | 0.326 |
| scenario1 | RW3 | 1.5 | 1.75 | 4 | 0.573613 | 0.531 |
| scenario1 | RW3 | 1.5 | 1.75 | 5 | 0.709665 | 0.713 |
| scenario1 | RW3 | 1.5 | 1.75 | 6 | 0.807628 | 0.84 |
| scenario1 | RW3 | 1.5 | 1.75 | 7 | 0.875376 | 0.911 |

| | | | | | | |
|---|---|---|---|---|---|---|
| scenario1 | RW3 | 1.5 | 1.75 | 8 | 0.920793 | 0.948 |
| scenario1 | RW3 | 1.5 | 1.75 | 9 | 0.950484 | 0.965 |
| scenario1 | RW3 | 1.5 | 1.75 | 10 | 0.969489 | 0.984 |
| scenario1 | RW3 | 1.5 | 1.75 | 11 | 0.981439 | 0.988 |
| scenario1 | RW3 | 1.5 | 1.75 | 12 | 0.988837 | 0.993 |
| scenario1 | RW3 | 1.5 | 1.75 | 13 | 0.993356 | 1 |
| scenario1 | RW3 | 1.5 | 1.75 | 14 | 0.996082 | 0.999 |
| scenario1 | RW3 | 1.5 | 1.75 | 15 | 0.997709 | 1 |
| scenario1 | RW3 | 1.5 | 1.75 | 16 | 0.998671 | 1 |
| scenario1 | RW3 | 1.5 | 1.75 | 17 | 0.999235 | 1 |
| scenario1 | RW3 | 1.5 | 1.75 | 18 | 0.999562 | 1 |
| scenario1 | RW3 | 1.5 | 1.75 | 19 | 0.999751 | 1 |
| scenario1 | RW3 | 1.5 | 1.75 | 20 | 0.999859 | 1 |
| scenario1 | RW4 | 1.25 | 1.5 | 3 | 0.23097 | 0.097 |
| scenario1 | RW4 | 1.25 | 1.5 | 4 | 0.336757 | 0.227 |
| scenario1 | RW4 | 1.25 | 1.5 | 5 | 0.435076 | 0.373 |
| scenario1 | RW4 | 1.25 | 1.5 | 6 | 0.523952 | 0.524 |
| scenario1 | RW4 | 1.25 | 1.5 | 7 | 0.602643 | 0.613 |
| scenario1 | RW4 | 1.25 | 1.5 | 8 | 0.671138 | 0.707 |
| scenario1 | RW4 | 1.25 | 1.5 | 9 | 0.729908 | 0.78 |
| scenario1 | RW4 | 1.25 | 1.5 | 10 | 0.779715 | 0.839 |
| scenario1 | RW4 | 1.25 | 1.5 | 11 | 0.821473 | 0.878 |
| scenario1 | RW4 | 1.25 | 1.5 | 12 | 0.856155 | 0.916 |
| scenario1 | RW4 | 1.25 | 1.5 | 13 | 0.884717 | 0.952 |
| scenario1 | RW4 | 1.25 | 1.5 | 14 | 0.908064 | 0.956 |
| scenario1 | RW4 | 1.25 | 1.5 | 15 | 0.927018 | 0.977 |
| scenario1 | RW4 | 1.25 | 1.5 | 16 | 0.942311 | 0.974 |
| scenario1 | RW4 | 1.25 | 1.5 | 17 | 0.954581 | 0.984 |
| scenario1 | RW4 | 1.25 | 1.5 | 18 | 0.964375 | 0.994 |
| scenario1 | RW4 | 1.25 | 1.5 | 19 | 0.972154 | 0.997 |
| scenario1 | RW4 | 1.25 | 1.5 | 20 | 0.978307 | 0.999 |
| scenario1 | RW4 | 1.25 | 1.75 | 3 | 0.687757 | 0.638 |
| scenario1 | RW4 | 1.25 | 1.75 | 4 | 0.88014 | 0.909 |
| scenario1 | RW4 | 1.25 | 1.75 | 5 | 0.958143 | 0.98 |
| scenario1 | RW4 | 1.25 | 1.75 | 6 | 0.986291 | 1 |
| scenario1 | RW4 | 1.25 | 1.75 | 7 | 0.995723 | 1 |
| scenario1 | RW4 | 1.25 | 1.75 | 8 | 0.998717 | 1 |
| scenario1 | RW4 | 1.25 | 1.75 | 9 | 0.999627 | 1 |
| scenario1 | RW4 | 1.25 | 1.75 | 10 | 0.999895 | 1 |
| scenario1 | RW4 | 1.25 | 1.75 | 11 | 0.999971 | 1 |
| scenario1 | RW4 | 1.25 | 1.75 | 12 | 0.999992 | 1 |
| scenario1 | RW4 | 1.25 | 1.75 | 13 | 0.999998 | 1 |
| scenario1 | RW4 | 1.25 | 1.75 | 14 | 0.999999 | 1 |
| scenario1 | RW4 | 1.25 | 1.75 | 15 | 1 | 1 |
| scenario1 | RW4 | 1.25 | 1.75 | 16 | 1 | 1 |
| scenario1 | RW4 | 1.25 | 1.75 | 17 | 1 | 1 |
| scenario1 | RW4 | 1.25 | 1.75 | 18 | 1 | 1 |
| scenario1 | RW4 | 1.25 | 1.75 | 19 | 1 | 1 |
| scenario1 | RW4 | 1.25 | 1.75 | 20 | 1 | 1 |
| scenario1 | RW4 | 1.5 | 1.75 | 3 | 0.268223 | 0.144 |

| | | | | | | |
|---|---|---|---|---|---|---|
| scenario1 | RW4 | 1.5 | 1.75 | 4 | 0.393601 | 0.353 |
| scenario1 | RW4 | 1.5 | 1.75 | 5 | 0.505979 | 0.486 |
| scenario1 | RW4 | 1.5 | 1.75 | 6 | 0.603222 | 0.641 |
| scenario1 | RW4 | 1.5 | 1.75 | 7 | 0.685242 | 0.711 |
| scenario1 | RW4 | 1.5 | 1.75 | 8 | 0.753018 | 0.798 |
| scenario1 | RW4 | 1.5 | 1.75 | 9 | 0.808073 | 0.867 |
| scenario1 | RW4 | 1.5 | 1.75 | 10 | 0.85215 | 0.894 |
| scenario1 | RW4 | 1.5 | 1.75 | 11 | 0.886998 | 0.928 |
| scenario1 | RW4 | 1.5 | 1.75 | 12 | 0.914249 | 0.954 |
| scenario1 | RW4 | 1.5 | 1.75 | 13 | 0.935353 | 0.971 |
| scenario1 | RW4 | 1.5 | 1.75 | 14 | 0.951556 | 0.979 |
| scenario1 | RW4 | 1.5 | 1.75 | 15 | 0.963898 | 0.986 |
| scenario1 | RW4 | 1.5 | 1.75 | 16 | 0.973234 | 0.993 |
| scenario1 | RW4 | 1.5 | 1.75 | 17 | 0.98025 | 0.993 |
| scenario1 | RW4 | 1.5 | 1.75 | 18 | 0.985492 | 0.997 |
| scenario1 | RW4 | 1.5 | 1.75 | 19 | 0.989387 | 0.998 |
| scenario1 | RW4 | 1.5 | 1.75 | 20 | 0.992267 | 0.999 |
| scenario2 | RW1 | 2.5 | 2 | 3 | 0.216262 | 0.093 |
| scenario2 | RW1 | 2.5 | 2 | 4 | 0.314148 | 0.226 |
| scenario2 | RW1 | 2.5 | 2 | 5 | 0.406298 | 0.34 |
| scenario2 | RW1 | 2.5 | 2 | 6 | 0.490902 | 0.495 |
| scenario2 | RW1 | 2.5 | 2 | 7 | 0.567124 | 0.564 |
| scenario2 | RW1 | 2.5 | 2 | 8 | 0.634727 | 0.637 |
| scenario2 | RW1 | 2.5 | 2 | 9 | 0.693894 | 0.736 |
| scenario2 | RW1 | 2.5 | 2 | 10 | 0.745084 | 0.795 |
| scenario2 | RW1 | 2.5 | 2 | 11 | 0.788932 | 0.84 |
| scenario2 | RW1 | 2.5 | 2 | 12 | 0.82616 | 0.879 |
| scenario2 | RW1 | 2.5 | 2 | 13 | 0.857519 | 0.917 |
| scenario2 | RW1 | 2.5 | 2 | 14 | 0.883748 | 0.94 |
| scenario2 | RW1 | 2.5 | 2 | 15 | 0.905547 | 0.96 |
| scenario2 | RW1 | 2.5 | 2 | 16 | 0.923559 | 0.951 |
| scenario2 | RW1 | 2.5 | 2 | 17 | 0.938362 | 0.983 |
| scenario2 | RW1 | 2.5 | 2 | 18 | 0.95047 | 0.988 |
| scenario2 | RW1 | 2.5 | 2 | 19 | 0.960328 | 0.99 |
| scenario2 | RW1 | 2.5 | 2 | 20 | 0.968321 | 0.994 |
| scenario2 | RW1 | 2.5 | 3 | 3 | 0.260089 | 0.149 |
| scenario2 | RW1 | 2.5 | 3 | 4 | 0.381317 | 0.313 |
| scenario2 | RW1 | 2.5 | 3 | 5 | 0.490889 | 0.479 |
| scenario2 | RW1 | 2.5 | 3 | 6 | 0.586654 | 0.615 |
| scenario2 | RW1 | 2.5 | 3 | 7 | 0.668322 | 0.688 |
| scenario2 | RW1 | 2.5 | 3 | 8 | 0.736604 | 0.785 |
| scenario2 | RW1 | 2.5 | 3 | 9 | 0.792759 | 0.848 |
| scenario2 | RW1 | 2.5 | 3 | 10 | 0.838295 | 0.899 |
| scenario2 | RW1 | 2.5 | 3 | 11 | 0.874775 | 0.934 |
| scenario2 | RW1 | 2.5 | 3 | 12 | 0.903689 | 0.936 |
| scenario2 | RW1 | 2.5 | 3 | 13 | 0.926391 | 0.959 |
| scenario2 | RW1 | 2.5 | 3 | 14 | 0.944067 | 0.977 |
| scenario2 | RW1 | 2.5 | 3 | 15 | 0.957724 | 0.991 |
| scenario2 | RW1 | 2.5 | 3 | 16 | 0.968205 | 0.988 |
| scenario2 | RW1 | 2.5 | 3 | 17 | 0.976197 | 0.995 |

| | | | | | | |
|---|---|---|---|---|---|---|
| scenario2 | RW1 | 2.5 | 3 | 18 | 0.982257 | 0.997 |
| scenario2 | RW1 | 2.5 | 3 | 19 | 0.986828 | 0.993 |
| scenario2 | RW1 | 2.5 | 3 | 20 | 0.990258 | 1 |
| scenario2 | RW1 | 2 | 3 | 3 | 0.666007 | 0.651 |
| scenario2 | RW1 | 2 | 3 | 4 | 0.863858 | 0.926 |
| scenario2 | RW1 | 2 | 3 | 5 | 0.949285 | 0.981 |
| scenario2 | RW1 | 2 | 3 | 6 | 0.982234 | 0.993 |
| scenario2 | RW1 | 2 | 3 | 7 | 0.994061 | 0.999 |
| scenario2 | RW1 | 2 | 3 | 8 | 0.998088 | 1 |
| scenario2 | RW1 | 2 | 3 | 9 | 0.999404 | 1 |
| scenario2 | RW1 | 2 | 3 | 10 | 0.999819 | 1 |
| scenario2 | RW1 | 2 | 3 | 11 | 0.999946 | 1 |
| scenario2 | RW1 | 2 | 3 | 12 | 0.999984 | 1 |
| scenario2 | RW1 | 2 | 3 | 13 | 0.999996 | 1 |
| scenario2 | RW1 | 2 | 3 | 14 | 0.999999 | 1 |
| scenario2 | RW1 | 2 | 3 | 15 | 1 | 1 |
| scenario2 | RW1 | 2 | 3 | 16 | 1 | 1 |
| scenario2 | RW1 | 2 | 3 | 17 | 1 | 1 |
| scenario2 | RW1 | 2 | 3 | 18 | 1 | 1 |
| scenario2 | RW1 | 2 | 3 | 19 | 1 | 1 |
| scenario2 | RW1 | 2 | 3 | 20 | 1 | 1 |
| scenario2 | RW2 | 2.5 | 2 | 3 | 0.359537 | 0.257 |
| scenario2 | RW2 | 2.5 | 2 | 4 | 0.525265 | 0.457 |
| scenario2 | RW2 | 2.5 | 2 | 5 | 0.658394 | 0.665 |
| scenario2 | RW2 | 2.5 | 2 | 6 | 0.759893 | 0.786 |
| scenario2 | RW2 | 2.5 | 2 | 7 | 0.834512 | 0.879 |
| scenario2 | RW2 | 2.5 | 2 | 8 | 0.887842 | 0.93 |
| scenario2 | RW2 | 2.5 | 2 | 9 | 0.925089 | 0.951 |
| scenario2 | RW2 | 2.5 | 2 | 10 | 0.950607 | 0.978 |
| scenario2 | RW2 | 2.5 | 2 | 11 | 0.967803 | 0.989 |
| scenario2 | RW2 | 2.5 | 2 | 12 | 0.979227 | 0.994 |
| scenario2 | RW2 | 2.5 | 2 | 13 | 0.986722 | 1 |
| scenario2 | RW2 | 2.5 | 2 | 14 | 0.991584 | 1 |
| scenario2 | RW2 | 2.5 | 2 | 15 | 0.994707 | 1 |
| scenario2 | RW2 | 2.5 | 2 | 16 | 0.996694 | 0.998 |
| scenario2 | RW2 | 2.5 | 2 | 17 | 0.997949 | 1 |
| scenario2 | RW2 | 2.5 | 2 | 18 | 0.998735 | 1 |
| scenario2 | RW2 | 2.5 | 2 | 19 | 0.999224 | 1 |
| scenario2 | RW2 | 2.5 | 2 | 20 | 0.999527 | 1 |
| scenario2 | RW2 | 2.5 | 3 | 3 | 0.398224 | 0.317 |
| scenario2 | RW2 | 2.5 | 3 | 4 | 0.576745 | 0.552 |
| scenario2 | RW2 | 2.5 | 3 | 5 | 0.71261 | 0.715 |
| scenario2 | RW2 | 2.5 | 3 | 6 | 0.810132 | 0.812 |
| scenario2 | RW2 | 2.5 | 3 | 7 | 0.877365 | 0.896 |
| scenario2 | RW2 | 2.5 | 3 | 8 | 0.922296 | 0.954 |
| scenario2 | RW2 | 2.5 | 3 | 9 | 0.951575 | 0.964 |
| scenario2 | RW2 | 2.5 | 3 | 10 | 0.970256 | 0.987 |
| scenario2 | RW2 | 2.5 | 3 | 11 | 0.981964 | 0.984 |
| scenario2 | RW2 | 2.5 | 3 | 12 | 0.989188 | 0.992 |
| scenario2 | RW2 | 2.5 | 3 | 13 | 0.993586 | 0.998 |

| | | | | | | |
|---|---|---|---|---|---|---|
| scenario2 | RW2 | 2.5 | 3 | 14 | 0.99623 | 0.999 |
| scenario2 | RW2 | 2.5 | 3 | 15 | 0.997803 | 1 |
| scenario2 | RW2 | 2.5 | 3 | 16 | 0.99873 | 0.999 |
| scenario2 | RW2 | 2.5 | 3 | 17 | 0.999271 | 1 |
| scenario2 | RW2 | 2.5 | 3 | 18 | 0.999584 | 1 |
| scenario2 | RW2 | 2.5 | 3 | 19 | 0.999764 | 1 |
| scenario2 | RW2 | 2.5 | 3 | 20 | 0.999867 | 1 |
| scenario2 | RW2 | 2 | 3 | 3 | 0.883214 | 0.887 |
| scenario2 | RW2 | 2 | 3 | 4 | 0.9826 | 0.991 |
| scenario2 | RW2 | 2 | 3 | 5 | 0.997804 | 0.999 |
| scenario2 | RW2 | 2 | 3 | 6 | 0.99975 | 1 |
| scenario2 | RW2 | 2 | 3 | 7 | 0.999973 | 1 |
| scenario2 | RW2 | 2 | 3 | 8 | 0.999997 | 1 |
| scenario2 | RW2 | 2 | 3 | 9 | 1 | 1 |
| scenario2 | RW2 | 2 | 3 | 10 | 1 | 1 |
| scenario2 | RW2 | 2 | 3 | 11 | 1 | 1 |
| scenario2 | RW2 | 2 | 3 | 12 | 1 | 1 |
| scenario2 | RW2 | 2 | 3 | 13 | 1 | 1 |
| scenario2 | RW2 | 2 | 3 | 14 | 1 | 1 |
| scenario2 | RW2 | 2 | 3 | 15 | 1 | 1 |
| scenario2 | RW2 | 2 | 3 | 16 | 1 | 1 |
| scenario2 | RW2 | 2 | 3 | 17 | 1 | 1 |
| scenario2 | RW2 | 2 | 3 | 18 | 1 | 1 |
| scenario2 | RW2 | 2 | 3 | 19 | 1 | 1 |
| scenario2 | RW2 | 2 | 3 | 20 | 1 | 1 |
| scenario2 | RW3 | 2.5 | 2 | 3 | 0.497135 | 0.385 |
| scenario2 | RW3 | 2.5 | 2 | 4 | 0.699513 | 0.7 |
| scenario2 | RW3 | 2.5 | 2 | 5 | 0.82973 | 0.87 |
| scenario2 | RW3 | 2.5 | 2 | 6 | 0.90722 | 0.946 |
| scenario2 | RW3 | 2.5 | 2 | 7 | 0.950999 | 0.977 |
| scenario2 | RW3 | 2.5 | 2 | 8 | 0.974782 | 0.994 |
| scenario2 | RW3 | 2.5 | 2 | 9 | 0.987305 | 0.997 |
| scenario2 | RW3 | 2.5 | 2 | 10 | 0.993729 | 0.999 |
| scenario2 | RW3 | 2.5 | 2 | 11 | 0.996954 | 1 |
| scenario2 | RW3 | 2.5 | 2 | 12 | 0.998542 | 1 |
| scenario2 | RW3 | 2.5 | 2 | 13 | 0.999311 | 1 |
| scenario2 | RW3 | 2.5 | 2 | 14 | 0.999678 | 1 |
| scenario2 | RW3 | 2.5 | 2 | 15 | 0.999852 | 1 |
| scenario2 | RW3 | 2.5 | 2 | 16 | 0.999932 | 1 |
| scenario2 | RW3 | 2.5 | 2 | 17 | 0.999969 | 1 |
| scenario2 | RW3 | 2.5 | 2 | 18 | 0.999986 | 1 |
| scenario2 | RW3 | 2.5 | 2 | 19 | 0.999994 | 1 |
| scenario2 | RW3 | 2.5 | 2 | 20 | 0.999997 | 1 |
| scenario2 | RW3 | 2.5 | 3 | 3 | 0.481935 | 0.396 |
| scenario2 | RW3 | 2.5 | 3 | 4 | 0.681214 | 0.653 |
| scenario2 | RW3 | 2.5 | 3 | 5 | 0.813334 | 0.814 |
| scenario2 | RW3 | 2.5 | 3 | 6 | 0.894663 | 0.916 |
| scenario2 | RW3 | 2.5 | 3 | 7 | 0.942296 | 0.961 |
| scenario2 | RW3 | 2.5 | 3 | 8 | 0.969161 | 0.986 |
| scenario2 | RW3 | 2.5 | 3 | 9 | 0.983862 | 0.993 |

| | | | | | | |
|---|---|---|---|---|---|---|
| scenario2 | RW3 | 2.5 | 3 | 10 | 0.991708 | 0.997 |
| scenario2 | RW3 | 2.5 | 3 | 11 | 0.995807 | 1 |
| scenario2 | RW3 | 2.5 | 3 | 12 | 0.99791 | 1 |
| scenario2 | RW3 | 2.5 | 3 | 13 | 0.998971 | 1 |
| scenario2 | RW3 | 2.5 | 3 | 14 | 0.999499 | 1 |
| scenario2 | RW3 | 2.5 | 3 | 15 | 0.999759 | 1 |
| scenario2 | RW3 | 2.5 | 3 | 16 | 0.999885 | 1 |
| scenario2 | RW3 | 2.5 | 3 | 17 | 0.999946 | 1 |
| scenario2 | RW3 | 2.5 | 3 | 18 | 0.999974 | 1 |
| scenario2 | RW3 | 2.5 | 3 | 19 | 0.999988 | 1 |
| scenario2 | RW3 | 2.5 | 3 | 20 | 0.999995 | 1 |
| scenario2 | RW3 | 2 | 3 | 3 | 0.949539 | 0.943 |
| scenario2 | RW3 | 2 | 3 | 4 | 0.996932 | 0.999 |
| scenario2 | RW3 | 2 | 3 | 5 | 0.999853 | 1 |
| scenario2 | RW3 | 2 | 3 | 6 | 0.999994 | 1 |
| scenario2 | RW3 | 2 | 3 | 7 | 1 | 1 |
| scenario2 | RW3 | 2 | 3 | 8 | 1 | 1 |
| scenario2 | RW3 | 2 | 3 | 9 | 1 | 1 |
| scenario2 | RW3 | 2 | 3 | 10 | 1 | 1 |
| scenario2 | RW3 | 2 | 3 | 11 | 1 | 1 |
| scenario2 | RW3 | 2 | 3 | 12 | 1 | 1 |
| scenario2 | RW3 | 2 | 3 | 13 | 1 | 1 |
| scenario2 | RW3 | 2 | 3 | 14 | 1 | 1 |
| scenario2 | RW3 | 2 | 3 | 15 | 1 | 1 |
| scenario2 | RW3 | 2 | 3 | 16 | 1 | 1 |
| scenario2 | RW3 | 2 | 3 | 17 | 1 | 1 |
| scenario2 | RW3 | 2 | 3 | 18 | 1 | 1 |
| scenario2 | RW3 | 2 | 3 | 19 | 1 | 1 |
| scenario2 | RW3 | 2 | 3 | 20 | 1 | 1 |
| scenario2 | RW4 | 2.5 | 2 | 3 | 0.18206 | 0.05 |
| scenario2 | RW4 | 2.5 | 2 | 4 | 0.260273 | 0.137 |
| scenario2 | RW4 | 2.5 | 2 | 5 | 0.335841 | 0.259 |
| scenario2 | RW4 | 2.5 | 2 | 6 | 0.407565 | 0.361 |
| scenario2 | RW4 | 2.5 | 2 | 7 | 0.47469 | 0.447 |
| scenario2 | RW4 | 2.5 | 2 | 8 | 0.536757 | 0.548 |
| scenario2 | RW4 | 2.5 | 2 | 9 | 0.593548 | 0.618 |
| scenario2 | RW4 | 2.5 | 2 | 10 | 0.645033 | 0.667 |
| scenario2 | RW4 | 2.5 | 2 | 11 | 0.69133 | 0.712 |
| scenario2 | RW4 | 2.5 | 2 | 12 | 0.73266 | 0.783 |
| scenario2 | RW4 | 2.5 | 2 | 13 | 0.769316 | 0.828 |
| scenario2 | RW4 | 2.5 | 2 | 14 | 0.801637 | 0.86 |
| scenario2 | RW4 | 2.5 | 2 | 15 | 0.829984 | 0.896 |
| scenario2 | RW4 | 2.5 | 2 | 16 | 0.854724 | 0.921 |
| scenario2 | RW4 | 2.5 | 2 | 17 | 0.876221 | 0.94 |
| scenario2 | RW4 | 2.5 | 2 | 18 | 0.894822 | 0.949 |
| scenario2 | RW4 | 2.5 | 2 | 19 | 0.910856 | 0.96 |
| scenario2 | RW4 | 2.5 | 2 | 20 | 0.92463 | 0.968 |
| scenario2 | RW4 | 2.5 | 3 | 3 | 0.20234 | 0.08 |
| scenario2 | RW4 | 2.5 | 3 | 4 | 0.292384 | 0.203 |
| scenario2 | RW4 | 2.5 | 3 | 5 | 0.378114 | 0.332 |

| | | | | | | |
|---|---|---|---|---|---|---|
| scenario2 | RW4 | 2.5 | 3 | 6 | 0.457943 | 0.44 |
| scenario2 | RW4 | 2.5 | 3 | 7 | 0.531024 | 0.527 |
| scenario2 | RW4 | 2.5 | 3 | 8 | 0.596981 | 0.63 |
| scenario2 | RW4 | 2.5 | 3 | 9 | 0.655786 | 0.698 |
| scenario2 | RW4 | 2.5 | 3 | 10 | 0.707661 | 0.753 |
| scenario2 | RW4 | 2.5 | 3 | 11 | 0.752999 | 0.782 |
| scenario2 | RW4 | 2.5 | 3 | 12 | 0.792299 | 0.841 |
| scenario2 | RW4 | 2.5 | 3 | 13 | 0.826116 | 0.881 |
| scenario2 | RW4 | 2.5 | 3 | 14 | 0.855024 | 0.903 |
| scenario2 | RW4 | 2.5 | 3 | 15 | 0.879588 | 0.922 |
| scenario2 | RW4 | 2.5 | 3 | 16 | 0.900347 | 0.942 |
| scenario2 | RW4 | 2.5 | 3 | 17 | 0.917803 | 0.958 |
| scenario2 | RW4 | 2.5 | 3 | 18 | 0.932415 | 0.974 |
| scenario2 | RW4 | 2.5 | 3 | 19 | 0.944595 | 0.975 |
| scenario2 | RW4 | 2.5 | 3 | 20 | 0.954706 | 0.97 |
| scenario2 | RW4 | 2 | 3 | 3 | 0.549267 | 0.484 |
| scenario2 | RW4 | 2 | 3 | 4 | 0.756218 | 0.789 |
| scenario2 | RW4 | 2 | 3 | 5 | 0.87621 | 0.914 |
| scenario2 | RW4 | 2 | 3 | 6 | 0.939932 | 0.982 |
| scenario2 | RW4 | 2 | 3 | 7 | 0.971876 | 0.994 |
| scenario2 | RW4 | 2 | 3 | 8 | 0.987214 | 0.999 |
| scenario2 | RW4 | 2 | 3 | 9 | 0.994329 | 1 |
| scenario2 | RW4 | 2 | 3 | 10 | 0.997538 | 1 |
| scenario2 | RW4 | 2 | 3 | 11 | 0.998951 | 1 |
| scenario2 | RW4 | 2 | 3 | 12 | 0.99956 | 1 |
| scenario2 | RW4 | 2 | 3 | 13 | 0.999818 | 1 |
| scenario2 | RW4 | 2 | 3 | 14 | 0.999926 | 1 |
| scenario2 | RW4 | 2 | 3 | 15 | 0.99997 | 1 |
| scenario2 | RW4 | 2 | 3 | 16 | 0.999988 | 1 |
| scenario2 | RW4 | 2 | 3 | 17 | 0.999995 | 1 |
| scenario2 | RW4 | 2 | 3 | 18 | 0.999998 | 1 |
| scenario2 | RW4 | 2 | 3 | 19 | 0.999999 | 1 |
| scenario2 | RW4 | 2 | 3 | 20 | 1 | 1 |
| scenario3 | RW1 | 0.2 | 0.5 | 3 | 0.064134 | 0.007 |
| scenario3 | RW1 | 0.2 | 0.5 | 4 | 0.07217 | 0.018 |
| scenario3 | RW1 | 0.2 | 0.5 | 5 | 0.080158 | 0.015 |
| scenario3 | RW1 | 0.2 | 0.5 | 6 | 0.088135 | 0.032 |
| scenario3 | RW1 | 0.2 | 0.5 | 7 | 0.096122 | 0.037 |
| scenario3 | RW1 | 0.2 | 0.5 | 8 | 0.104127 | 0.053 |
| scenario3 | RW1 | 0.2 | 0.5 | 9 | 0.112154 | 0.053 |
| scenario3 | RW1 | 0.2 | 0.5 | 10 | 0.120206 | 0.059 |
| scenario3 | RW1 | 0.2 | 0.5 | 11 | 0.12828 | 0.078 |
| scenario3 | RW1 | 0.2 | 0.5 | 12 | 0.136376 | 0.088 |
| scenario3 | RW1 | 0.2 | 0.5 | 13 | 0.144491 | 0.083 |
| scenario3 | RW1 | 0.2 | 0.5 | 14 | 0.152624 | 0.096 |
| scenario3 | RW1 | 0.2 | 0.5 | 15 | 0.16077 | 0.111 |
| scenario3 | RW1 | 0.2 | 0.5 | 16 | 0.168928 | 0.143 |
| scenario3 | RW1 | 0.2 | 0.5 | 17 | 0.177095 | 0.123 |
| scenario3 | RW1 | 0.2 | 0.5 | 18 | 0.185268 | 0.13 |
| scenario3 | RW1 | 0.2 | 0.5 | 19 | 0.193444 | 0.15 |

| | | | | | | |
|---|---|---|---|---|---|---|
| scenario3 | RW1 | 0.2 | 0.5 | 20 | 0.201621 | 0.143 |
| scenario3 | RW1 | 0.2 | 0.8 | 3 | 0.124939 | 0.039 |
| scenario3 | RW1 | 0.2 | 0.8 | 4 | 0.169138 | 0.084 |
| scenario3 | RW1 | 0.2 | 0.8 | 5 | 0.213098 | 0.118 |
| scenario3 | RW1 | 0.2 | 0.8 | 6 | 0.256563 | 0.197 |
| scenario3 | RW1 | 0.2 | 0.8 | 7 | 0.299301 | 0.254 |
| scenario3 | RW1 | 0.2 | 0.8 | 8 | 0.34109 | 0.327 |
| scenario3 | RW1 | 0.2 | 0.8 | 9 | 0.381735 | 0.354 |
| scenario3 | RW1 | 0.2 | 0.8 | 10 | 0.421073 | 0.397 |
| scenario3 | RW1 | 0.2 | 0.8 | 11 | 0.458973 | 0.454 |
| scenario3 | RW1 | 0.2 | 0.8 | 12 | 0.495335 | 0.504 |
| scenario3 | RW1 | 0.2 | 0.8 | 13 | 0.530089 | 0.504 |
| scenario3 | RW1 | 0.2 | 0.8 | 14 | 0.563188 | 0.563 |
| scenario3 | RW1 | 0.2 | 0.8 | 15 | 0.594609 | 0.62 |
| scenario3 | RW1 | 0.2 | 0.8 | 16 | 0.624347 | 0.675 |
| scenario3 | RW1 | 0.2 | 0.8 | 17 | 0.652413 | 0.667 |
| scenario3 | RW1 | 0.2 | 0.8 | 18 | 0.678834 | 0.719 |
| scenario3 | RW1 | 0.2 | 0.8 | 19 | 0.703644 | 0.766 |
| scenario3 | RW1 | 0.2 | 0.8 | 20 | 0.726889 | 0.778 |
| scenario3 | RW1 | 0.5 | 0.8 | 3 | 0.073602 | 0.007 |
| scenario3 | RW1 | 0.5 | 0.8 | 4 | 0.087137 | 0.022 |
| scenario3 | RW1 | 0.5 | 0.8 | 5 | 0.100636 | 0.026 |
| scenario3 | RW1 | 0.5 | 0.8 | 6 | 0.114143 | 0.046 |
| scenario3 | RW1 | 0.5 | 0.8 | 7 | 0.127682 | 0.065 |
| scenario3 | RW1 | 0.5 | 0.8 | 8 | 0.141257 | 0.085 |
| scenario3 | RW1 | 0.5 | 0.8 | 9 | 0.154866 | 0.09 |
| scenario3 | RW1 | 0.5 | 0.8 | 10 | 0.1685 | 0.104 |
| scenario3 | RW1 | 0.5 | 0.8 | 11 | 0.182149 | 0.135 |
| scenario3 | RW1 | 0.5 | 0.8 | 12 | 0.195804 | 0.134 |
| scenario3 | RW1 | 0.5 | 0.8 | 13 | 0.209452 | 0.138 |
| scenario3 | RW1 | 0.5 | 0.8 | 14 | 0.223084 | 0.159 |
| scenario3 | RW1 | 0.5 | 0.8 | 15 | 0.236687 | 0.171 |
| scenario3 | RW1 | 0.5 | 0.8 | 16 | 0.250252 | 0.224 |
| scenario3 | RW1 | 0.5 | 0.8 | 17 | 0.263768 | 0.203 |
| scenario3 | RW1 | 0.5 | 0.8 | 18 | 0.277226 | 0.217 |
| scenario3 | RW1 | 0.5 | 0.8 | 19 | 0.290616 | 0.265 |
| scenario3 | RW1 | 0.5 | 0.8 | 20 | 0.303929 | 0.257 |
| scenario3 | RW2 | 0.2 | 0.5 | 3 | 0.075136 | 0.02 |
| scenario3 | RW2 | 0.2 | 0.5 | 4 | 0.08957 | 0.043 |
| scenario3 | RW2 | 0.2 | 0.5 | 5 | 0.103971 | 0.065 |
| scenario3 | RW2 | 0.2 | 0.5 | 6 | 0.118384 | 0.067 |
| scenario3 | RW2 | 0.2 | 0.5 | 7 | 0.13283 | 0.086 |
| scenario3 | RW2 | 0.2 | 0.5 | 8 | 0.147313 | 0.1 |
| scenario3 | RW2 | 0.2 | 0.5 | 9 | 0.161827 | 0.115 |
| scenario3 | RW2 | 0.2 | 0.5 | 10 | 0.176363 | 0.119 |
| scenario3 | RW2 | 0.2 | 0.5 | 11 | 0.190908 | 0.16 |
| scenario3 | RW2 | 0.2 | 0.5 | 12 | 0.205449 | 0.158 |
| scenario3 | RW2 | 0.2 | 0.5 | 13 | 0.219973 | 0.177 |
| scenario3 | RW2 | 0.2 | 0.5 | 14 | 0.234468 | 0.213 |
| scenario3 | RW2 | 0.2 | 0.5 | 15 | 0.24892 | 0.209 |

| | | | | | | |
|---|---|---|---|---|---|---|
| scenario3 | RW2 | 0.2 | 0.5 | 16 | 0.263318 | 0.227 |
| scenario3 | RW2 | 0.2 | 0.5 | 17 | 0.277649 | 0.249 |
| scenario3 | RW2 | 0.2 | 0.5 | 18 | 0.291902 | 0.256 |
| scenario3 | RW2 | 0.2 | 0.5 | 19 | 0.306068 | 0.28 |
| scenario3 | RW2 | 0.2 | 0.5 | 20 | 0.320136 | 0.27 |
| scenario3 | RW2 | 0.2 | 0.8 | 3 | 0.187254 | 0.09 |
| scenario3 | RW2 | 0.2 | 0.8 | 4 | 0.268468 | 0.183 |
| scenario3 | RW2 | 0.2 | 0.8 | 5 | 0.346653 | 0.288 |
| scenario3 | RW2 | 0.2 | 0.8 | 6 | 0.420516 | 0.414 |
| scenario3 | RW2 | 0.2 | 0.8 | 7 | 0.489273 | 0.463 |
| scenario3 | RW2 | 0.2 | 0.8 | 8 | 0.552473 | 0.54 |
| scenario3 | RW2 | 0.2 | 0.8 | 9 | 0.609934 | 0.594 |
| scenario3 | RW2 | 0.2 | 0.8 | 10 | 0.661678 | 0.647 |
| scenario3 | RW2 | 0.2 | 0.8 | 11 | 0.707883 | 0.743 |
| scenario3 | RW2 | 0.2 | 0.8 | 12 | 0.748834 | 0.757 |
| scenario3 | RW2 | 0.2 | 0.8 | 13 | 0.784883 | 0.83 |
| scenario3 | RW2 | 0.2 | 0.8 | 14 | 0.816428 | 0.844 |
| scenario3 | RW2 | 0.2 | 0.8 | 15 | 0.843878 | 0.885 |
| scenario3 | RW2 | 0.2 | 0.8 | 16 | 0.867646 | 0.911 |
| scenario3 | RW2 | 0.2 | 0.8 | 17 | 0.888131 | 0.92 |
| scenario3 | RW2 | 0.2 | 0.8 | 18 | 0.905712 | 0.931 |
| scenario3 | RW2 | 0.2 | 0.8 | 19 | 0.920742 | 0.941 |
| scenario3 | RW2 | 0.2 | 0.8 | 20 | 0.933544 | 0.954 |
| scenario3 | RW2 | 0.5 | 0.8 | 3 | 0.094092 | 0.03 |
| scenario3 | RW2 | 0.5 | 0.8 | 4 | 0.119757 | 0.062 |
| scenario3 | RW2 | 0.5 | 0.8 | 5 | 0.145417 | 0.104 |
| scenario3 | RW2 | 0.5 | 0.8 | 6 | 0.171074 | 0.124 |
| scenario3 | RW2 | 0.5 | 0.8 | 7 | 0.196701 | 0.134 |
| scenario3 | RW2 | 0.5 | 0.8 | 8 | 0.222252 | 0.173 |
| scenario3 | RW2 | 0.5 | 0.8 | 9 | 0.247671 | 0.2 |
| scenario3 | RW2 | 0.5 | 0.8 | 10 | 0.272899 | 0.204 |
| scenario3 | RW2 | 0.5 | 0.8 | 11 | 0.297883 | 0.279 |
| scenario3 | RW2 | 0.5 | 0.8 | 12 | 0.322569 | 0.267 |
| scenario3 | RW2 | 0.5 | 0.8 | 13 | 0.346911 | 0.306 |
| scenario3 | RW2 | 0.5 | 0.8 | 14 | 0.370866 | 0.354 |
| scenario3 | RW2 | 0.5 | 0.8 | 15 | 0.394397 | 0.357 |
| scenario3 | RW2 | 0.5 | 0.8 | 16 | 0.417472 | 0.391 |
| scenario3 | RW2 | 0.5 | 0.8 | 17 | 0.44006 | 0.435 |
| scenario3 | RW2 | 0.5 | 0.8 | 18 | 0.46214 | 0.439 |
| scenario3 | RW2 | 0.5 | 0.8 | 19 | 0.483689 | 0.475 |
| scenario3 | RW2 | 0.5 | 0.8 | 20 | 0.504691 | 0.503 |
| scenario3 | RW3 | 0.2 | 0.5 | 3 | 0.085904 | 0.036 |
| scenario3 | RW3 | 0.2 | 0.5 | 4 | 0.106688 | 0.045 |
| scenario3 | RW3 | 0.2 | 0.5 | 5 | 0.127462 | 0.085 |
| scenario3 | RW3 | 0.2 | 0.5 | 6 | 0.148255 | 0.092 |
| scenario3 | RW3 | 0.2 | 0.5 | 7 | 0.169071 | 0.12 |
| scenario3 | RW3 | 0.2 | 0.5 | 8 | 0.18989 | 0.133 |
| scenario3 | RW3 | 0.2 | 0.5 | 9 | 0.210684 | 0.151 |
| scenario3 | RW3 | 0.2 | 0.5 | 10 | 0.231418 | 0.169 |
| scenario3 | RW3 | 0.2 | 0.5 | 11 | 0.252061 | 0.214 |

| | | | | | | |
|---|---|---|---|---|---|---|
| scenario3 | RW3 | 0.2 | 0.5 | 12 | 0.272578 | 0.234 |
| scenario3 | RW3 | 0.2 | 0.5 | 13 | 0.29294 | 0.25 |
| scenario3 | RW3 | 0.2 | 0.5 | 14 | 0.313115 | 0.277 |
| scenario3 | RW3 | 0.2 | 0.5 | 15 | 0.333077 | 0.306 |
| scenario3 | RW3 | 0.2 | 0.5 | 16 | 0.352802 | 0.337 |
| scenario3 | RW3 | 0.2 | 0.5 | 17 | 0.372266 | 0.353 |
| scenario3 | RW3 | 0.2 | 0.5 | 18 | 0.391449 | 0.362 |
| scenario3 | RW3 | 0.2 | 0.5 | 19 | 0.410334 | 0.364 |
| scenario3 | RW3 | 0.2 | 0.5 | 20 | 0.428903 | 0.427 |
| scenario3 | RW3 | 0.2 | 0.8 | 3 | 0.248201 | 0.14 |
| scenario3 | RW3 | 0.2 | 0.8 | 4 | 0.363065 | 0.293 |
| scenario3 | RW3 | 0.2 | 0.8 | 5 | 0.468119 | 0.401 |
| scenario3 | RW3 | 0.2 | 0.8 | 6 | 0.561264 | 0.534 |
| scenario3 | RW3 | 0.2 | 0.8 | 7 | 0.641981 | 0.649 |
| scenario3 | RW3 | 0.2 | 0.8 | 8 | 0.710636 | 0.708 |
| scenario3 | RW3 | 0.2 | 0.8 | 9 | 0.768128 | 0.796 |
| scenario3 | RW3 | 0.2 | 0.8 | 10 | 0.815633 | 0.844 |
| scenario3 | RW3 | 0.2 | 0.8 | 11 | 0.854435 | 0.893 |
| scenario3 | RW3 | 0.2 | 0.8 | 12 | 0.885806 | 0.936 |
| scenario3 | RW3 | 0.2 | 0.8 | 13 | 0.910944 | 0.944 |
| scenario3 | RW3 | 0.2 | 0.8 | 14 | 0.930924 | 0.963 |
| scenario3 | RW3 | 0.2 | 0.8 | 15 | 0.94669 | 0.974 |
| scenario3 | RW3 | 0.2 | 0.8 | 16 | 0.95905 | 0.981 |
| scenario3 | RW3 | 0.2 | 0.8 | 17 | 0.96868 | 0.991 |
| scenario3 | RW3 | 0.2 | 0.8 | 18 | 0.976143 | 0.988 |
| scenario3 | RW3 | 0.2 | 0.8 | 19 | 0.981897 | 0.996 |
| scenario3 | RW3 | 0.2 | 0.8 | 20 | 0.986312 | 0.998 |
| scenario3 | RW3 | 0.5 | 0.8 | 3 | 0.115016 | 0.047 |
| scenario3 | RW3 | 0.5 | 0.8 | 4 | 0.153237 | 0.084 |
| scenario3 | RW3 | 0.5 | 0.8 | 5 | 0.191354 | 0.136 |
| scenario3 | RW3 | 0.5 | 0.8 | 6 | 0.229218 | 0.168 |
| scenario3 | RW3 | 0.5 | 0.8 | 7 | 0.266678 | 0.22 |
| scenario3 | RW3 | 0.5 | 0.8 | 8 | 0.303577 | 0.251 |
| scenario3 | RW3 | 0.5 | 0.8 | 9 | 0.339766 | 0.292 |
| scenario3 | RW3 | 0.5 | 0.8 | 10 | 0.375113 | 0.326 |
| scenario3 | RW3 | 0.5 | 0.8 | 11 | 0.409506 | 0.39 |
| scenario3 | RW3 | 0.5 | 0.8 | 12 | 0.442852 | 0.426 |
| scenario3 | RW3 | 0.5 | 0.8 | 13 | 0.475076 | 0.458 |
| scenario3 | RW3 | 0.5 | 0.8 | 14 | 0.50612 | 0.498 |
| scenario3 | RW3 | 0.5 | 0.8 | 15 | 0.535944 | 0.531 |
| scenario3 | RW3 | 0.5 | 0.8 | 16 | 0.56452 | 0.596 |
| scenario3 | RW3 | 0.5 | 0.8 | 17 | 0.591833 | 0.618 |
| scenario3 | RW3 | 0.5 | 0.8 | 18 | 0.61788 | 0.634 |
| scenario3 | RW3 | 0.5 | 0.8 | 19 | 0.642665 | 0.684 |
| scenario3 | RW3 | 0.5 | 0.8 | 20 | 0.666202 | 0.719 |
| scenario3 | RW4 | 0.2 | 0.5 | 3 | 0.060185 | 0.003 |
| scenario3 | RW4 | 0.2 | 0.5 | 4 | 0.065952 | 0.007 |
| scenario3 | RW4 | 0.2 | 0.5 | 5 | 0.071675 | 0.016 |
| scenario3 | RW4 | 0.2 | 0.5 | 6 | 0.077382 | 0.021 |
| scenario3 | RW4 | 0.2 | 0.5 | 7 | 0.083089 | 0.022 |

| | | | | | | |
|---|---|---|---|---|---|---|
| scenario3 | RW4 | 0.2 | 0.5 | 8 | 0.088804 | 0.036 |
| scenario3 | RW4 | 0.2 | 0.5 | 9 | 0.094532 | 0.04 |
| scenario3 | RW4 | 0.2 | 0.5 | 10 | 0.100275 | 0.049 |
| scenario3 | RW4 | 0.2 | 0.5 | 11 | 0.106032 | 0.044 |
| scenario3 | RW4 | 0.2 | 0.5 | 12 | 0.111806 | 0.053 |
| scenario3 | RW4 | 0.2 | 0.5 | 13 | 0.117594 | 0.059 |
| scenario3 | RW4 | 0.2 | 0.5 | 14 | 0.123397 | 0.059 |
| scenario3 | RW4 | 0.2 | 0.5 | 15 | 0.129213 | 0.069 |
| scenario3 | RW4 | 0.2 | 0.5 | 16 | 0.135041 | 0.069 |
| scenario3 | RW4 | 0.2 | 0.5 | 17 | 0.140881 | 0.077 |
| scenario3 | RW4 | 0.2 | 0.5 | 18 | 0.146732 | 0.095 |
| scenario3 | RW4 | 0.2 | 0.5 | 19 | 0.152592 | 0.093 |
| scenario3 | RW4 | 0.2 | 0.5 | 20 | 0.15846 | 0.11 |
| scenario3 | RW4 | 0.2 | 0.8 | 3 | 0.104575 | 0.018 |
| scenario3 | RW4 | 0.2 | 0.8 | 4 | 0.136513 | 0.046 |
| scenario3 | RW4 | 0.2 | 0.8 | 5 | 0.168422 | 0.082 |
| scenario3 | RW4 | 0.2 | 0.8 | 6 | 0.200242 | 0.109 |
| scenario3 | RW4 | 0.2 | 0.8 | 7 | 0.231894 | 0.145 |
| scenario3 | RW4 | 0.2 | 0.8 | 8 | 0.263281 | 0.189 |
| scenario3 | RW4 | 0.2 | 0.8 | 9 | 0.294303 | 0.226 |
| scenario3 | RW4 | 0.2 | 0.8 | 10 | 0.324867 | 0.275 |
| scenario3 | RW4 | 0.2 | 0.8 | 11 | 0.354887 | 0.331 |
| scenario3 | RW4 | 0.2 | 0.8 | 12 | 0.384287 | 0.348 |
| scenario3 | RW4 | 0.2 | 0.8 | 13 | 0.413003 | 0.408 |
| scenario3 | RW4 | 0.2 | 0.8 | 14 | 0.44098 | 0.401 |
| scenario3 | RW4 | 0.2 | 0.8 | 15 | 0.468173 | 0.467 |
| scenario3 | RW4 | 0.2 | 0.8 | 16 | 0.494545 | 0.499 |
| scenario3 | RW4 | 0.2 | 0.8 | 17 | 0.520069 | 0.533 |
| scenario3 | RW4 | 0.2 | 0.8 | 18 | 0.544723 | 0.549 |
| scenario3 | RW4 | 0.2 | 0.8 | 19 | 0.568494 | 0.573 |
| scenario3 | RW4 | 0.2 | 0.8 | 20 | 0.591375 | 0.616 |
| scenario3 | RW4 | 0.5 | 0.8 | 3 | 0.067378 | 0.003 |
| scenario3 | RW4 | 0.5 | 0.8 | 4 | 0.077288 | 0.014 |
| scenario3 | RW4 | 0.5 | 0.8 | 5 | 0.087152 | 0.022 |
| scenario3 | RW4 | 0.5 | 0.8 | 6 | 0.097012 | 0.03 |
| scenario3 | RW4 | 0.5 | 0.8 | 7 | 0.106889 | 0.038 |
| scenario3 | RW4 | 0.5 | 0.8 | 8 | 0.116793 | 0.058 |
| scenario3 | RW4 | 0.5 | 0.8 | 9 | 0.126726 | 0.059 |
| scenario3 | RW4 | 0.5 | 0.8 | 10 | 0.136688 | 0.087 |
| scenario3 | RW4 | 0.5 | 0.8 | 11 | 0.146676 | 0.083 |
| scenario3 | RW4 | 0.5 | 0.8 | 12 | 0.156686 | 0.096 |
| scenario3 | RW4 | 0.5 | 0.8 | 13 | 0.166714 | 0.095 |
| scenario3 | RW4 | 0.5 | 0.8 | 14 | 0.176754 | 0.103 |
| scenario3 | RW4 | 0.5 | 0.8 | 15 | 0.186802 | 0.132 |
| scenario3 | RW4 | 0.5 | 0.8 | 16 | 0.196853 | 0.126 |
| scenario3 | RW4 | 0.5 | 0.8 | 17 | 0.206903 | 0.121 |
| scenario3 | RW4 | 0.5 | 0.8 | 18 | 0.216946 | 0.156 |
| scenario3 | RW4 | 0.5 | 0.8 | 19 | 0.226978 | 0.154 |
| scenario3 | RW4 | 0.5 | 0.8 | 20 | 0.236995 | 0.176 |

Percentage Bin Legend - ATI in TIV:
1 = 0-10 %
2 = 11-25%
3 = 26-50%
4 = 51+ %
